# Ultra-Efficient Integration of Gene Libraries onto Yeast Cytosolic Plasmids

**DOI:** 10.1101/2024.11.29.626108

**Authors:** Alexander (Olek) Pisera, Yutong Yu, Rory L. Williams, Chang C. Liu

**Affiliations:** Department of Biomedical Engineering, University of California; Irvine, CA, 92617, USA; Center for Synthetic Biology, University of California; Irvine, CA, 92617, USA; Department of Pharmaceutical Sciences, University of California; Irvine, CA, 92617, USA; Department of Chemistry, University of California; Irvine, CA, 92617, USA; Department of Molecular Biology & Biochemistry, University of California; Irvine, CA, 92617, USA

## Abstract

Efficient methods for diversifying genes of interest (GOIs) are essential in protein engineering. For example, OrthoRep, a yeast-based orthogonal DNA replication system that achieves the rapid *in vivo* diversification of GOIs encoded on a cytosolic plasmid (p1), has been successfully used to drive numerous protein engineering campaigns. However, OrthoRep-based GOI evolution has almost always started from single GOI sequences, limiting the number of locations on a fitness landscape from where evolutionary search begins. Here, we present a simple approach for the high-efficiency integration of GOI libraries onto OrthoRep. By leveraging integrases, we demonstrate recombination of donor DNA onto the cytosolic p1 plasmid at exceptionally high transformation efficiencies, even surpassing the transformation efficiency of standard circular plasmids into yeast. We demonstrate our method’s utility through the straightforward construction of mock nanobody libraries encoded on OrthoRep, from which rare binders were reliably enriched. Overall, integrase-assisted manipulation of yeast cytosolic plasmids should enhance the versatility of OrthoRep in continuous evolution experiments and support the routine construction of large GOI libraries in yeast in general.

## Introduction

Orthogonal DNA replication is an architecture that enables the continuous hypermutation and evolution of user-selected genes *in vivo*^1-6^. In the yeast instantiation of orthogonal DNA replication called OrthoRep, genes of interest (GOIs) are encoded onto a cytosolic DNA plasmid (p1) that is durably and exclusively replicated by an error-prone orthogonal DNA polymerase (DNAP)^2^. The DNAP only replicates p1 such that GOIs experience a high mutation rate, while the *Saccharomyces cerevisiae* host genome is maintained at its normal and necessarily low mutation rates. When selection pressures are imposed, the hypermutating GOI can rapidly adapt to achieve new or improved biomolecular function. OrthoRep has been applied to biosynthetic enzyme engineering^7-9^, antibody generation^10,11^, transcription factor evolution^12^, gene editor evolution^13^, plant enzyme engineering^14-16^, and other biomolecular engineering problems^17^. It has also been applied to the broad exploration of fitness landscapes in the interest of understanding mutational pathways to drug resistance^2^, detecting meaningful patterns of conservation and change during protein evolution^18,19^, and producing large synthetic evolutionary datasets that may be useful as probes of and training sets for ML models^19,20^. However, OrthoRep-driven GOI evolution campaigns have almost always started from a single or few GOI sequences encoded on p1, representing evolutionary search from limited locations on a fitness landscape. To expand the search space that OrthoRep-driven GOI evolution experiments access, it would be desirable to start from a large diversity of GOI sequences (*i*.*e*., libraries) rather than a few. To enable this, we present a strategy for high-efficiency integration of GOIs onto p1 and demonstrate the advantages of this strategy in both routine OrthoRep workflows and through the straightforward generation of mock antibody libraries from which productive binders can be selected. Notably, the efficiency of integrating GOIs onto p1 with our strategy is exceptionally high and exceeds even the efficiency of transforming standard DNA plasmids into yeast^21,22^. This finding predicts the extension of our strategy’s value beyond OrthoRep-driven continuous evolution into yeast-based protein engineering and yeast genetics at large.

## Results

### Rationale

Currently, installing GOIs onto OrthoRep’s p1 plasmid is done by transforming *S. cerevisiae* cells with a linear donor DNA cassette containing the GOI^1,23^. This cassette is flanked by homologous sequences that support recombination onto a landing pad p1 already present inside cells. Selection for successful recombinants results in the isolation of the desired OrthoRep strain encoding the GOI on p1. Our reliance on transformation and recombination to install donor DNA onto p1, versus *ex vivo* cloning of GOIs onto p1 followed by transformation into cells, stems from the fact that replication-active p1s have terminal proteins (TPs) covalently linked to their 5’ ends^1,24,25^. These TPs make it challenging to work with p1 in a test tube^26^, since standard molecular cloning procedures and ingredients are designed for DNA, not DNA-protein conjugates. However, the need for recombination onto a landing pad p1 comes with its own challenge. The p1 plasmid is cytosolic and likely has little access to yeast’s endogenous recombination machinery localized in the nuclear compartment^27,28^. Thus, the efficiency with which GOIs are installed onto p1 is generally low. We reasoned that by deliberately expressing DNA recombinases in the cytosol where p1 resides, we could achieve high levels of donor DNA integration onto p1 (Fig. 1c).

**Figure 1.**
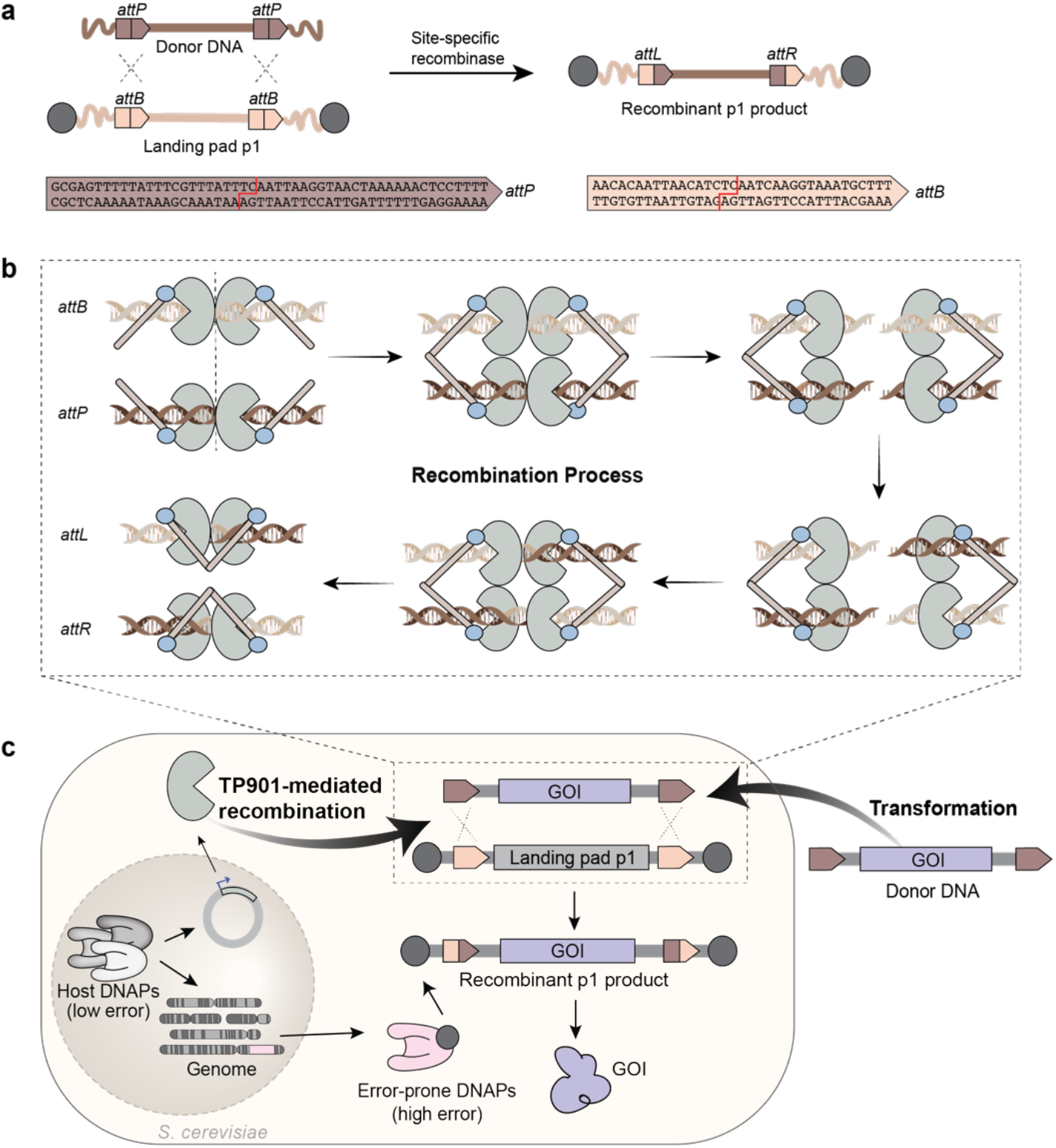
The design of TP901-mediated integration of donor DNA onto the p1 of the OrthoRep system. **a**, Schematic of the recombination process between the landing pad p1 and donor DNA utilizing a site-specific recombinase. The landing pad p1 with *attB* recombination sites and terminal proteins (indicated by dark gray filled circles) recombines with the donor DNA containing *attP* sites at a specific dinucleotide (indicated in red). This process, facilitated by TP901 integrase, results in a recombined p1 plasmid with newly formed *attL* and *attR* sites. **b**, Mechanism of TP901-dependent recombination process. TP901 initially binds to the *attB* and *attP* sites as dimers, then forms a tetramer to align the sites. The *attB* and *attP* sites are cleaved, generating 3’ overhangs. Subsequently, two integrase subunits rotate, swapping the *attB* and *attP* half-sites, which are then re-ligated to form *attL* and *attR* sites. **c**, TP901-mediated integration in the context of the OrthoRep system in *S. cerevisiae*. Transformation allows the donor DNA containing GOI to enter the cell, but replication of the donor DNA only occurs if it is integrated onto the cytosolic landing pad p1. The resulting recombinant p1 product is durably and exclusively replicated by an error-prone orthogonal DNAP at a high mutation rate to drive hypermutation of the GOI while the host genome is maintained with a low mutation rate.

### Specific design and testing

Serine integrases are site-specific recombinases that mediate the exchange and rejoining of DNA at attachment sites^29^ (Fig. 1a and Fig. 1b). We wished to test whether the highly active serine integrase, TP901, could achieve recombination between attachment sites in donor DNA and landing pad p1s in the yeast cytosol. To do so, we 1) cloned donor DNA constructs where *attP* directional recombination sites flanked our GOI and a selectable auxotrophic marker (*e*.*g*., *LEU2*), 2) created yeast strains that maintain a landing pad p1 with complementary *attB* recombination sites flanking a *URA3* selectable auxotrophic marker, and 3) made a nuclear 2*µ* plasmid that encodes TP901 for cytosolic expression. When *attP*-containing donor DNA constructs were transformed into chemically competent TP901-expressing strains with *attB*-containing landing pad p1s, we observed efficient integration. Specifically, the number of colonies that successfully integrated the donor DNA was 20-to 60-fold higher when TP901 was expressed (Fig. 2a). It is important to note that our *attP*-containing donor DNA was flanked by the same homologous sequences we historically used for TP901-independent recombination onto p1^1^. Therefore, the large TP901-dependent increase in the number of successful integrants (Fig. 2b) should be interpreted as improvement over the previous state-of-the-art (SOTA) onto p1 where successful integration was already substantial. We conclude that TP901 supports the efficient cytosolic recombination of donor DNA onto p1.

**Figure 2.**
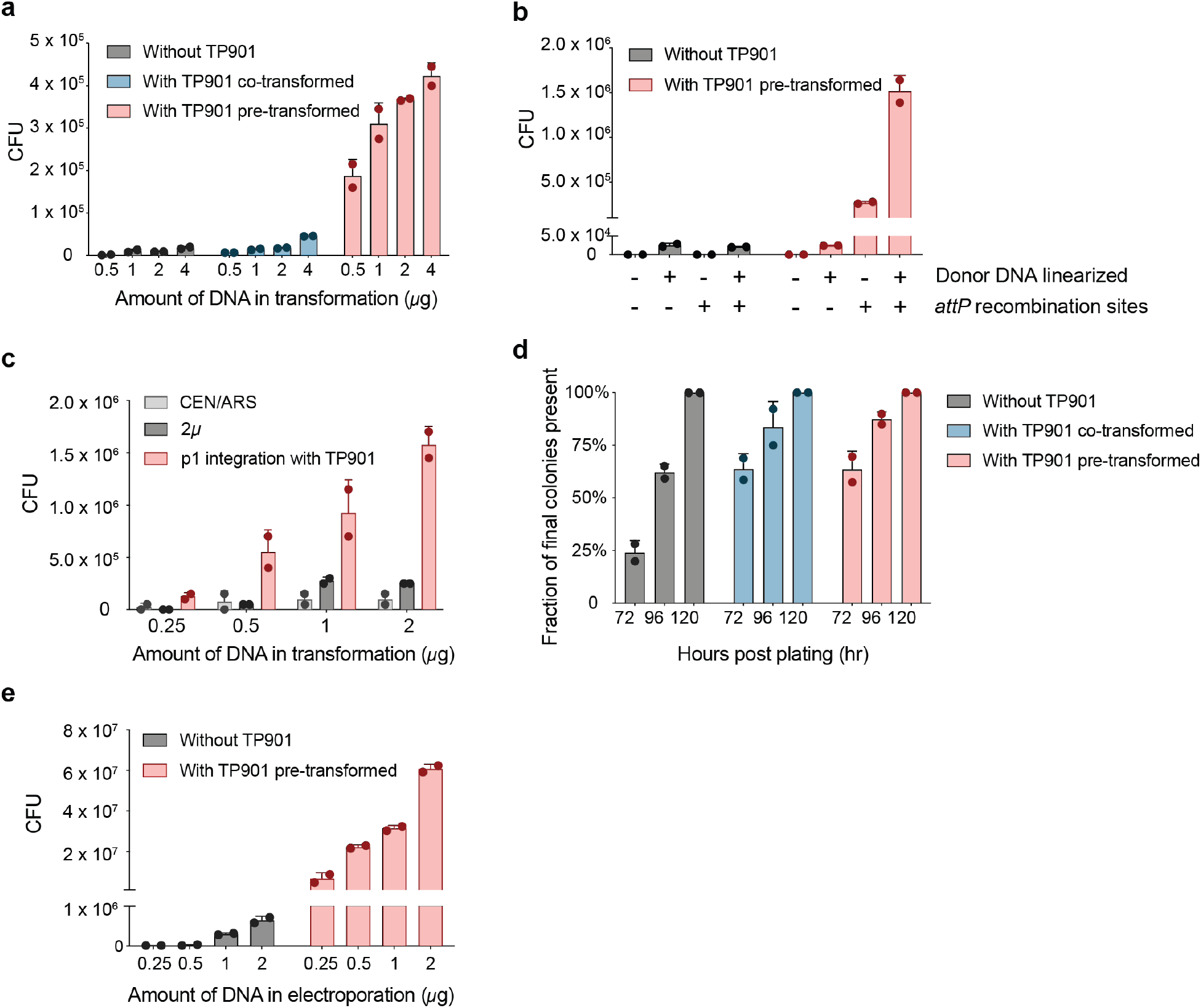
TP901 significantly enhances the efficiency of integrating GOIs onto the OrthoRep system. **a**, Comparison of transformation efficiency with or without TP901. Cells containing both the TP901 integrase and TP901-compatible landing pad p1 yielded 25-to 60-times more colonies than comparable conditions not utilizing TP901. **b**, Comparison of transformation efficiency across different conditions. Conditions tested include the use of linearized donor DNA and the presence of *attP* recombination sites on the donor DNA. The highest transformation efficiency was observed with the combination of linearized donor DNA and *attP* recombination sites. **c**, Efficiency of TP901-dependent integration onto p1 compared to standard transformation with circular plasmid (CEN/ARS and 2*µ* plasmids). The TP901 system yielded higher CFUs across different DNA amounts, outperforming conventional plasmid transformation. **d**, Accelerated colony formation with TP901 integrase. Colonies appeared more rapidly in the presence of TP901 integrase, with most colonies observed within 72 hours, in contrast to slower colony formation without integrase. **e**, Comparison of electroporation efficiency with or without TP901 integrase. By utilizing the TP901 system and electroporation, the transformation of GOI onto OrthoRep was highly efficient. Each condition was measured in duplicate biological replicates. The mean and range (error bars) are shown alongside individual measurements indicated as points.

We next investigated practical variations for implementing TP901-assisted integration onto p1. A simple variation is to introduce the TP901 expression plasmid at the same time as donor DNA is transformed (co-transformation). This variation resulted in a modest boost in efficiency (∼2.5-fold over the previous SOTA) than if the TP901 plasmid was already expressed in all the recipient cells (Fig. 2a). A second variation we explored was whether we could transform donor DNA as a circular plasmid rather than as a linear cassette. As shown in Fig. 2b, when the donor DNA contained *attP* sites and the recipient cell expressed TP901, circular form donor DNA resulted in efficient integration. However, when the donor DNA did not contain *attP* sites, no productive integration occurred for the circular form even though homology flanks for recombination onto the landing pad p1 were present. This is presumably because the homology flanks need to be revealed at the ends of a linear cassette to support TP901-independent recombination. In short, circular or linear donor DNA with *attP* sites were both suitable for TP901-dependent integration of GOIs onto landing pad p1s containing *attB* sites whereas only linear donor DNA with homologous arms achieved integration in the absence of TP901. We also observed that the efficiency of TP901-dependent recombination was 6-fold lower for *attP*-containing donor DNA in the circular *versus* linear form, but both forms supported much higher integration efficiencies than our previous TP901-independent SOTA method.

### Efficiency over standard transformation of nuclear plasmids into yeast

We wished to test TP901-dependent integration of GOIs onto p1 against the transformation efficiency of standard nuclear plasmids, which is typically how gene libraries are introduced into yeast in traditional directed evolution, library screening, and yeast genetics experiments that don’t use OrthoRep. As shown in Fig. 2c representing side-by-side transformation experiments, TP901-dependent integration of a selectable marker onto landing pad p1s led to the appearance of 3.3-fold or 6.3-fold more colonies than the transformation of a 2*µ* plasmid containing the same marker at the 1 *µ*g and 2 *µ*g concentrations, respectively. And compared to the transformation of a CEN/ARS plasmid, TP901-dependent integration onto p1 led to the appearance of 9-fold and 15-fold more colonies. We therefore suggest that TP901-dependent integration onto p1 may be the most efficient way of introducing GOIs into yeast.

### Time to colony formation

When transforming library DNA into cells, colonies of successful transformants should ideally be the same size to ensure an even representation of library members. However, this is not typically the case with GOI integration onto p1 using our previous TP901-independent integration strategy. What we often observe is the first appearance of colonies (*i*.*e*., successful integrants) on day 3-4 after transformation followed by new colonies continuing to appear until day 7, resulting in large colony size variation that creates undesired biases in library composition. Large variation in the waiting times to colonies may be explained by the inefficiency of recombination onto p1. Assume that it is only after the first integration of donor DNA onto p1 that the cell can exponentially expand to form a colony. If recombination is inefficient, the average waiting time to the crucial first integration event may be similar to or even exceed yeast cell division times. The expected variation in waiting times to integration, a Poisson process, will then be characterized on cell division timescales, causing large variation in colony sizes, as those are also governed by cell division time. This is consistent with our previous observations.

Since TP901-mediated recombination is highly efficient, we reasoned that TP901-dependent integration may have short average waiting times. Should the expected waiting time be much shorter than cell division timescales governing colony formation, then waiting time variation will have little impact on colony formation time, colonies should appear faster within a smaller time window, and less variation in colony sizes should result. To test these predictions, we assessed the timing of colony formation for TP901-dependent transformation (Fig. 2d). With our previous TP901-independent SOTA method for GOI integration onto p1, only 25% of eventual colonies appeared on the first day of colony formation (day 3). With TP901-dependent GOI integration, 64% of eventual colonies appeared on the first day of colony formation, supporting our model (Fig. 2d). One notable detail to modify our model stems from the observation that transformation efficiency using different selection markers, for example, HIS3 versus LEU2, results in different number of colonies after selection, for example, donor DNA marked with HIS3 yielded approximately 4x more colonies than with LEU2 (Fig. S1). We suspect that this is due to differential sensitivity of yeast to amino acid deprivation for different amino acids, which renders inviable certain waiting times to integration in an amino acid dependent manner. Overall, our data suggest that TP901-dependent GOI integration supports the creation of evenly represented libraries while also improving the practical ease of encoding GOIs onto OrthoRep for evolution campaigns.

### Multiple integrations per cell

Since TP901-mediated recombination is highly efficient and since the landing pad p1 is maintained at high copy number, we suspected that multiple integration events could occur in the same cell, providing an extra boost in the number of library members installed onto OrthoRep, all else equal. To assess this, we transformed collections of different *attP*-containing donor DNA whose sizes are distinguishable on an agarose gel. We observed that single colonies usually contained ∼3 and up to 7 different p1s after TP901-mediated integration (Fig. S2a). When these single colonies containing multiple p1s were diluted and plated for single colonies again, we found that each subsequent colony only contained one p1 size (Fig. S2b). To further confirm that cells receiving DNA tend to integrate more than one p1 on average, we tested co-transformation of two p1 integration donor marked by two different selection markers, *HIS3* and *LEU2*. We found that number of colonies surviving selection for both markers was only slightly lower than the number of colonies surviving selection for only one or the other marker, suggesting that transformed cells commonly integrate more than one molecule of donor DNA (Fig. S1). Therefore, our overall model is that single cells uptake many molecules of donor DNA, the recombination efficiency of TP901 integrates them onto different copies of the landing pad p1s, and then early cell division events segregate those integrated copies into different cells. While the parameters governing how many different p1’s are generated per cell and how they segregate into sublineages have not been fully worked out, it is easy to imagine how this “multiplier” effect might be an advantage for building large libraries.

### Electroporation

The experiments described so far utilize chemical transformation to deliver donor DNA into strains for integration. However, it is known that electroporation can lead to higher transformation efficiency. We therefore wished to ensure that electroporation was compatible with TP901-dependent integration. As shown in Fig. 2e, integration efficiency was increased around 100-fold in the presence of TP901 such that in a routine single electroporation with 2 *µ*g of donor DNA, we can obtain 6 x 10^7^ colony-forming units (CFUs). As each CFU represents an average of ∼3 unique integrants (Fig. S2a), we should be able to construct libraries nearing 2 x 10^8^ members in only a single transformation. For comparison, the highest published transformation efficiencies for yeast claim ∼1 x 10^8^ CFU/*µ*g DNA^30^ and likely involve detailed optimization of electroporation conditions that our experiments lacked.

### Other integrases

The TP901 serine recombinase was chosen from a list of relatively equivalent options^29^. To test whether other integrases also support recombination onto p1, we constructed recipient strains with four other landing pads, donor DNA constructs containing the corresponding appropriate attachment sites, and 2*µ* plasmids expressing recombinases Bxb1, PhiBT1, PhiC31, and R4^31-34^. The R4 recombinase landing pad p1 consistently failed to receive donor DNA, likely driven by self-circularization of p1 owing to specific idiosyncrasies of R4 attachment sites. For the other three integrases, the efficiencies of integration were consistently higher than without integrase expression (Fig. 3).

**Figure 3.**
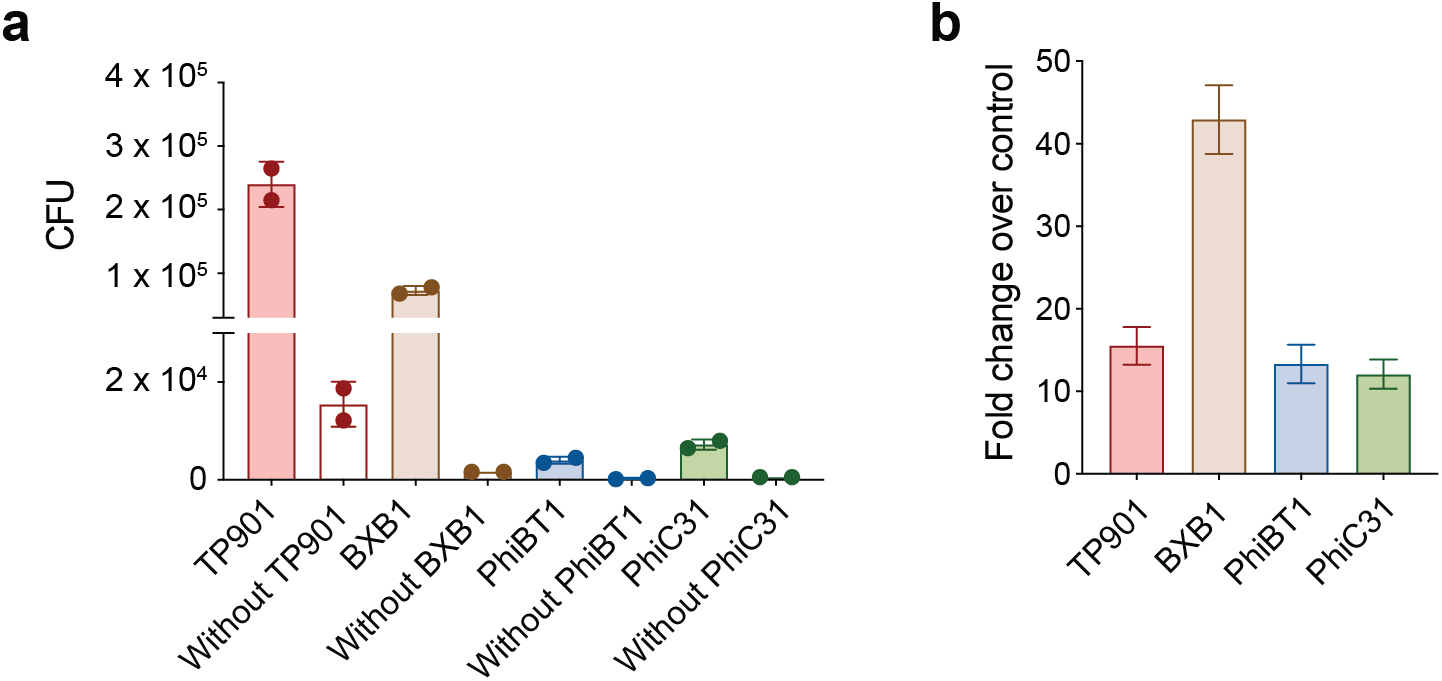
Comparison of integrase-mediated integration onto p1 across different integrases. **a**, Transformation efficiency resulting from the use of TP901, BXB1, PhiBT1, and PhiC31 integrases compared to without integrase. TP901 is the most effective integrase among those tested. **b**, Fold change in transformation efficiency over control for each integrase. Each condition was measured in duplicate biological replicates, with mean and range (error bars) shown.

### Library construction and selection

As a proof-of-concept for using TP901-dependent integration to generate GOI libraries that can be subject to functional selection, we carried out a mock nanobody (Nb) evolution experiment. The experiment mimics the loading of large nanobody libraries onto p1 followed by the enrichment of binders for further OrthoRep-driven evolution. Specifically, we mixed DNA encoding a previously described high-affinity Nb (RBD10i14)^10^ against the SARS-CoV-2 spike protein’s receptor binding domain (RBD) with DNA encoding a Nb that does not bind RBD (Nb.b201)^35^. 4 *µ*g of total DNA representing molar ratios of 1:10, 1:1,000, and 1:100,000 of RBD10i14:Nb.b201 were transformed into strains harboring landing pad p1s. Nbs were encoded such that they would display on the cell surface when they are expressed from p1, as previously described^11^. The resulting populations were then subjected to fluorescence-activated cell sorting (FACS) for binding to fluorescently labeled RBD. Even for the 1:100,000 RBD10i14:Nb.b201 mixture, clones encoding RBD10i14 were detectably enriched after just one round of sorting. After the second round of sorting, RBD10i14 clones dominated the population (Fig. 4 and Fig. S3). High-throughput sequencing revealed that after the third round of sorting for the 1:100,000 RBD10i14:Nb.b201 mixture, the population was almost exclusively composed of RBD10i14 (94.8%). This mock evolution experiment demonstrates that TP901-assisted generation of p1-encoded GOI libraries is reliable for the enrichment of rare binders. Current work in our lab is focusing on the construction of large antibody libraries encoded on p1 for the rapid generation of potent antibodies.

**Figure 4.**
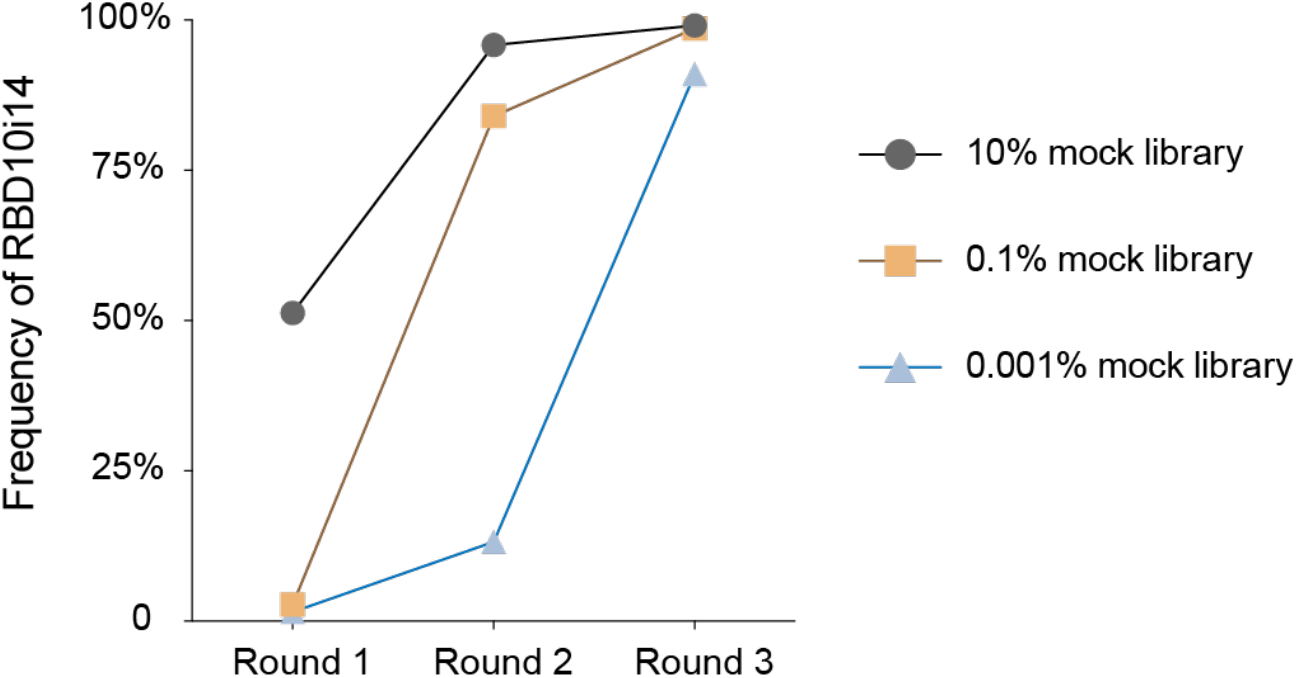
Enrichment of an RBD-binding nanobody (RBD10i14) from a mock yeast surface display nanobody library made through TP901-assisted integration onto p1. Three mock libraries with initial RBD10i14 frequencies of 10%, 0.1%, and 0.001% were efficiently loaded onto OrthoRep using TP901-assisted integration. Enrichment of RBD10i14 was successfully achieved through FACS over a total of 3 rounds of outgrowth and sorting performed. The data show progressive enrichment of RBD10i14 across all three libraries, demonstrating the capability to load large libraries onto OrthoRep and enrich target binders.

## Discussion

TP901-dependent high-efficiency integration of GOIs onto p1 should be of immediate utility to those using OrthoRep by increasing the speed and consistency of installing single genes onto OrthoRep for evolution. It also accesses the ability to install large gene libraries onto OrthoRep, which should further the power of continuous evolution in exploring and navigating fitness landscapes. Moreover, the combination of high-efficiency integration with our recent BadBoy OrthoRep DNAPs that mutate p1 at 10^−4^ substitutions per base^19,20^ should allow for the generation of unprecedented synthetic gene diversity in yeast. For example, if one were to encode ∼10^8^ diverse variants of a 1 kb GOI onto p1, which can be done in a single routine transformation, and then expand cells for 10 generations (2 days), one could yield a gene library of ∼10^11^ variants under realistic assumptions. It will be interesting to explore what such extreme gene diversity can yield.

Beyond the OrthoRep-specific utility of TP901-dependent integration of gene libraries, we emphasize that the transformation efficiencies we achieved exceed those of standard plasmid transformation and gene library construction methods in yeast. Therefore, our reported strategy should find broad utility in yeast genetics and protein engineering. We hypothesize that our high efficiencies stem from that fact that it is easier for donor DNA to enter the cytoplasm, where p1 resides, versus the nucleus, where standard plasmids and DNA constructs need to be for replication and expression. It will be interesting to explore other genetic engineering benefits of p1’s cytosolic localization (*e*.*g*., its potential protection from epigenetic modification and chromitinization) that make it orthogonal in ways beyond its orthogonal replication^36^.

## Author Contributions

A.P. conceived of the project idea with input from Y.Y., R.L.W., and C.C.L. A.P., Y.Y., R.L.W., and C.C.L. designed the experiments. A.P., Y.Y., and R.L.W. conducted the experiments. A.P., Y.Y., R.L.W., and C.C.L. analyzed the data. A.P., Y.Y., and C.C.L. wrote the manuscript with input from all authors.

## Acknowledgements

This work was funded by NIH R35GM136297 and NIH R01CA260415 to C.C.L. A.P. is supported by a Paul & Daisy Soros Fellowship for New Americans. Training grant support to Y.Y. is provided by NIH T32EY032448. R.L.W. is supported by a Hewitt Foundation for Medical Research Postdoctoral Fellowship. We thank the Institute for Rapid Antibody Engineering and Evolution, part of the Engineering+Health Initiative of the UCI Samueli School of Engineering, for additional support.

## Supplementary Information

Materials and Methods

Figure S1. Transformation efficiency comparison based on auxotrophic markers.

Figure S2. Agarose gel of p1 minipreps after transformation with donor DNA mixtures.

Figure S3. FACS plots for the enrichment of an RBD-binding nanobody (RBD10i14) from three mock libraries.

Table S1. Key plasmids used in this study.

Table S2. Key strains used in this study.

Supplementary References 36-38.

