## Supplementary Information for "Ultra-Efficient Integration of Gene Libraries onto Yeast Cytosolic Plasmids"

**Table of Contents:**

**Materials and Methods**

**Supplementary Figures**

Figure S1. Transfromation efficiency comparison based on auxotrophic markers.

Figure S2. Agarose gel of p1 minipreps after transformation with donor DNA mixtures.

Figure S3. FACS plots for the enrichment of an RBD-binding nanobody (RBD10i14) from three mock libraries.

**Supplementary Tables**

Table S1. Key plasmids used in this study.

Table S2. Key strains used in this study.

**Supplementary References 36-38**

### Materials and Methods

#### DNA plasmid construction

Key plasmids used in this study are listed in Table S1. All DNA templates for PCR were obtained from previous studies or ordered from gBlocks from IDT, and all primers used in this study were ordered from IDT. PCR was performed to generate amplicons by PrimeSTAR GXL DNA polymerase (TaKaRa). Plasmid construction was done *via* Gibson Assembly or Golden Gate Assembly (all enzymes ordered from NEB) and transformed into chemically competent *Escherichia coli* strain TOP10 (ThermoFisher). Clonal plasmids were sequence-verified through whole-plasmid sequencing (Plasmidsaurus).

#### Yeast strain construction and media

All yeast strains used in this study are listed in Table S2. The parent strain for this study, yOP109, has landing pad p1 that contains a *MET15* selectable auxotrophic marker but lacks the *attB* recombination sites. To engineer a strain suitable for TP901-assisted integration, a ScaI-linearized donor DNA from pYY13, which contains the *attB* sites, a *URA3* selectable auxotrophic marker, and homology flanks matching the landing pad p1 in yOP109 was transformed into yOP109. After selection for successful integrants, the deletion of the original landing pad p1 containing the *MET15* marker was performed as previously described[36], using the spacer sequence GCTAAGAAGTATCTATCTAA. This yielded the yeast strain yYY20. A 2*μ* plasmid encoding the gene of TP901 integrase (pYY10) was then transformed into yYY20, resulting in yeast strain yYY22.

For the construction of strain yYY235, we began with strain BJ5465 (ATCC: 208289). First, the selectable auxotrophic marker, *TRP1*, was fully deleted from the genome as previously described[36], using the spacer sequence ATGTCTGTTATTAATTTCAC. Next, a linearized donor DNA encoding wild-type TP-DNAP1, along with *URA3* and *CAN1* markers, was transformed and integrated into the Lyp1 locus. Finally, a linearized donor DNA containing β-estradiol inducible promoter driving Aga1p, together with the synthetic transcription factor that is responsible for induction, and a Hygromycin resistance marker, was transformed and integrated into the Aga1 locus, following the protocol described by Paulk *et al* [11]. The landing pad p1 with *attP* sites and *TRP1* selectable auxotrophic marker was transported into this engineered strain by abortive mating[37] with p1 donor strain yYY64 to create strain yYY206. Briefly, equal numbers of p1-containing cells (yYY64) and recipient cells were mixed, spun down, and plated on YPD for 6 hours at 30°C. After 6 hours, the mated cells were restreaked on an SC-HUWK plate supplemented with 50 *μ*g/mL S-Aminoethyl-l-cysteine (Sigma Aldrich). After 3 days, a single colony was picked into SC-HUW media for growth. Finally, a 2*μ* plasmid encoding TP901 integrase (pYY10) was transformed into yYY206, resulting in yeast strain yYY235. All genomic integration and the p1 transport were validated by a full yeast miniprep, PCR, agarose gel electrophoresis, and sequencing (Plasmidsaurus).

Yeast strains were grown in standard media, including yeast extract peptone dextrose (YPD) (10 g/L bacto yeast extract; 20 g/L bacto peptone; 20 g/L dextrose) and appropriate synthetic drop-out media (yeast nitrogen base w/o amino acids (US Biological), drop-out mix synthetic minus the appropriate nutrients w/o yeast nitrogen base (US Biological), and dextrose).

### Yeast p1 miniprep

Yeast minipreps to isolate p1 were performed as previously described[1]. 1.5 mL of saturated yeast culture was centrifuged, and the supernatant was discarded. The pellet was washed with 1 mL of 0.9% NaCl and resuspened in 250 $\mu$L of Zymolyase solution (0.9M sorbitol (Sigma Aldrich), 0.1 M EDTA (Sigma Aldrich), 10U/mL Zymolyase (US Biological)) before incubating at 37 °C for 1 hour with rotation. After incubation, the tube was centrifuged, the supernatant was discarded, and the pellet was resuspended in 280.5 $\mu$L of proteinase K solution (250 $\mu$L of TE buffer(50 mM Tris-HCl pH 7.5, 20 mM EDTA), 25 $\mu$L 10% SDS (Sigma Aldrich), and 5.5 $\mu$L 10mg/mL proteinase K (ThermoFisher)). The sample was incubated at 65 °C for 30 minutes, followed by the addition of 75 $\mu$L of 5 M potassium acetate (ThermoFisher) and incubated on ice for 30 minutes. The tube was centrifuged at 12,000 xg for 10 minutes, and the supernatant was collected and mixed with two volumes of 100% ethanol. After another centrifugation, the ethanol was removed, and the pellet was air-dried at room temperature. The pellet was then resuspended in 150 $\mu$L of TE buffer (50 mM Tris-HCl pH 7.5, 20 mM EDTA), followed by the addition of 8 $\mu$L of 1 mg/mL RNase A and incubated at 37 °C for 30 minutes. Next, 1 volume of isopropanol was added, and the tube was centrifuged at 12,000 xg for 15 minutes. The supernatant was discarded, and the pellet was air-dried at room temperature. The dry pellet was then resuspended in 30 $\mu$L of water, 20 uL of which was loaded onto a 0.9% agarose gel and run at 80V for 2 hours.

### Frozen competent cell preparation[38]

All yeast chemical transformations for testing efficiency in this study used cells prepared in this manner.

Yeast strains for transformation were grown in selective media at 30 °C with shaking at 200 rpm until near saturation. The $OD_{600}$ of the yeast culture was measured using a hemacytomer. Approximately 2.5 x $10^9$ cells were inoculated into 500 mL of YPD, and the culture was grown in the shaking incubator at 30 °C for 4-6 hours until reaching a cell density of 2 x $10^7$ cells/mL. Yeast cells were harvested by centrifugation at 3,000 x g for 5 minutes and washed with 0.5 volumes of sterile water, followed by a second wash with 0.01 volumes of sterile water. The cell pellet was resuspended in 0.01 volumes of filter sterile frozen competent cell solution (5% glycerol, 10% DMSO, both from ThermoFisher), and 50 $\mu$L aliquots were dispensed into 1.5 mL microcentrifuge tubes. These tubes were placed in a styrofoam container and stored at -80 °C.

### Transformation efficiency assessment

To evaluate transformation efficiency, we employed a chemical transformation protocol using frozen competent cells prepared as mentioned above. Frozen competent cells were thawed in a 37 °C water bath for 30 seconds and pelleted by centrifugation at 13,000 x g for 2 minutes. Frozen competent cell transformation mix (260 $\mu$L of 50% w/v PEG 3350 (ThermoFisher), 36 $\mu$L of 1 M LiAc (ThermoFisher), 50 $\mu$L of 2.0 mg/mL single-stranded carrier DNA (ThermoFisher), 14 $\mu$L of DNA plus sterile water) was added to the cell pellet, and the mixture was vortexed vigorously to resuspend the cells. The tube was then incubated in a 42 °C water bath for 30 minutes. After incubation, the cells were pelleted by centrifugation at 13,000 x g for 30 seconds, and the supernatant was removed. 1 mL of sterile water was added to resuspend the cells. Serial dilutions of the transformed cells were plated on appropriate SC drop-out plates, selecting for p1 integration to measure the transformation efficiency. Formed colonies on plates were counted at 72 hr, 96 hr, and 120 hr.

For specific assessments: (1) we tested the efficiency of different amounts of DNA with or without the use of TP901 integrase. 0.5, 1, 2, and 4 $\mu$g of pYY12 plasmid was digested by ScaI and validated by agarose gel electrophoresis. Those donor DNA were transformed into three yeast strains: yOP109 (without TP901), yYY22 (contained the TP901 plasmid, pYY10), and yYY20, which was co-transformed with 1 $\mu$g of TP901 plasmid, pYY10. Serial dilutions of the transformed cells were plated on SC-L plates, and colonies on the plates were counted after 5 days. (2) We also compared p1 integration efficiencies using linearized versus circular donor DNA from pYY12 (with *attP* sites) and pOP96 (without *attP* sites) into strains yOP109 (without TP901) and yYY22 (with TP901 plasmid pre-transformed). Serial dilutions of the transformed cells were plated on SC-L plates and colonies on the plates were counted after 5 days. (3) We assessed TP901-mediated p1 integration efficiency using ScaI-digested pYY12 donor DNA in yYY22, comparing it to transformations with standard circular CEN/ARS and 2$\mu$ plasmids that have the same selectable marker into the same yeast strain. All experiments were conducted in duplicate biological replicates, with mean values and ranges (error bars) presented alongside individual measurements.

Note that for transformations into strains with TP901 (either co-transformed or pre-transformed), we did not select for G418 resistance. Since TP901 facilitates the recombination process, its continued presence in the cell was not necessary post-recombination. By omitting G418 selection, we allowed for the natural loss (or curing) of the TP901 plasmid from the cells.

**Multi-p1 integration test**

To test the phenomena of multiple p1 integrations occurring in one cell, we cloned a mock library composed of random length DNA fragments ranging from 0.1 kb to 4 kb, into the pYY12 plasmid in place of the mKate gene using Gibson Assembly. The mock library plasmid was linearized with ScaI and 2 $\mu$g that linearized donor DNA was transformed into TP901 pre-transformed strain yYY22, and the product of transformation was streaked on an SC-L plate to get single colonies. 16 colonies were picked from a plate and grown to saturation in SC-L media, and DNA was prepared by a full yeast p1 miniprep for agarose gel electrophoresis. One colony with many visible p1's was selected for further characterization, and the liquid culture of that colony was re-streaked on a SC-L plate. Six colonies were grown up in liquid SC-L media and DNA was prepared by a full p1 miniprep followed by agarose gel electrophoresis.

**Electroporation**

A semi-saturated yeast culture of yYY22 was diluted into 100 mL of YPD to reach an $OD_{600}$ of 0.3 and grown at 30 °C for around 8 hours to reach an $OD_{600}$ of 1.6. Cells were harvested by centrifugation at 3000 rpm for 3 minutes at 4 °C, and the supernatant was discarded. The cell pellet was washed twice with 50 mL of ice-cold sterile water and once with 50 mL of ice-cold electroporation buffer (1 M sorbitol (Sigma Aldrich), 1 mM $CaCl_2$ (Sigma Aldrich)). The pellet was then resuspended in 20 mL of condition buffer (0.1 M LiAc (Sigma Aldrich), 10 mM DTT (Sigma Aldrich)) and incubated at 30 °C with shaking at 200 rpm. After conditioning, the cells were collected by centrifugation and washed again with 50 mL of ice-cold electroporation buffer. Finally, the cell pellet was resuspended in 1.2 mL of electroporation buffer, resulting in a cell density of approximately 2 x $10^9$ cells/mL, which was sufficient for 6 electroporation reactions of 200 $\mu$L each. The cells were kept on ice until electroporation.

For each electroporation, 8 mL of a 1:1 mix of 1 M sorbitol and 2x YPD was prepared and kept on ice. The p1 donor DNA was linearized from pYY12 by ScaI (NEB), and the digestion was confirmed by agarose gel electrophoresis. For each electroporation sample, 200 μL of electrocompetent cells, linearized p1 donor DNA, and 25 μg salmon sperm DNA (ThermoFisher) were gently mixed and incubated on ice for 5 minutes. The mixture was transferred to a pre-chilled BioRad GenePulser 0.2 cm cuvette and electroporated at 2.5 kV, 25 μF. 1 mL of pre-chilled sorbitol/YPD was added immediately to the cuvette after electroporation, and the contents were transferred to the prepared 1:1 sorbitol/YPD mix. The cells were incubated on a platform shaker at 200 rpm at 30 °C for 1 hour to recover. Following recovery, the cells were collected by centrifugation and washed with 0.9% NaCl. Serial dilutions of the cells were plated SC-L plates selected for p1 integration to measure the electroporation efficiency.

**Alternate integrase tests**

Strains with new landing pad p1s containing different *attB* sites from corresponding integrases (Bxb1, PhiBT1, PhiC31, and R4) were made by using an appropriate donor cassette with *URA3* markers. Specifically, strains with new landing pad p1s were transformed with a ScaI-lineared donor cassette with homology flanks streaked onto an SC-U plate from which colonies were picked, grown up, and screened by p1 miniprep and sequencing (Plasmidsaurus). For each integrase, a colony containing the landing pad p1 with corresponding *attB* sites and *URA3* marker was then transformed with a 2μ plasmid constitutively expressing the integrase with the same RNR2 promoter as used for TP901. One colony for each integrase was grown up and passaged 3x in SC-U + G418 before being used for a frozen competent cell preparation as mentioned above. Transformations of donor DNA using appropriate attachment sites were done in duplicate, side by side with control that lacked the attachment sites but contained flanking homology to the landing pad p1. 2 μg of linearized donor DNA was transformed per transformation. Colonies were counted after 5 days.

**Mock library construction and selection by fluorescence-activated cell sorting (FACS)**

The sequences for nanobody fragments, RBD10i14 and Nb.b201, were amplified by PCR using primers designed with overhangs compatible with Golden Gate Assembly. A Golden Gate Assembly reaction was then performed with the p1 donor vector plasmid (pYY62) and the nanobody sequence, along with BsaI-HF-V2 (NEB), T4 ligase (NEB), and T4 ligation buffer (NEB). The reaction was incubated at 50 °C for 1 hour. The assembled products were transformed into chemically competent *E. coli* strain TOP10 (ThermoFisher). Clonal plasmids (pYY82 for RBD10i14 and pYY83 for Nb.b201) were verified through whole plasmid sequencing (Plasmidsaurus). Plasmid concentrations were measured using Qubit Assays (ThermoFisher). Three plasmid libraries were created by mixing pYY82 and pYY83 in molar ratios of 1:10, 1:1000, and 1:100,000 for subsequent transformation.

4 μg of each mock library was linearized by ScaI and integrated into yeast strain yYY235 with *LEU2* as the selectable marker for recombinant p1. The transformants were directly inoculated into liquid SC-L media after transformation and grown for 2-3 days until the culture approached saturation. The yeast culture was then passaged at a 1:100 dilution into the induction media (SC-L with 200 nM β-estradiol) and grown at 30 °C with shaking at 200 rpm overnight. Approximately $5 \times 10^7$ induced yeast cells were harvested from the induction culture, washed twice with ice-cold HBSBM buffer (20 mM Tris-HCl pH 7.5, 100 mM NaCl, 0.1% BSA, and 5 mM maltose), and stained in 250 μL of a primary staining solution containing HBSBM buffer with 10 nM biotinylated-

RBD (SinoBiological, 40592-V08B-B) at 4 °C for 1 hour with rotation. After primary staining, the cells were washed once with HBSBM buffer and stained with 250 $\mu$L of secondary staining solution containing HBSBM buffer with 0.5 $\mu$L of 1 mg/mL streptavidin-AF647 (ThermoFisher) and 20 nM of anti-HA-AF488 (R&D system) for 20 minutes at 4 °C with rotation. Next, cells were washed twice with HBSBM buffer and resuspended in 4 mL of HBSBM buffer for sorting. FACS (Sony SH800) was used to sort 300 cells from the gated population of each sample (Fig. S3, red gate) into 3 mL of SC-L media. Sorted cells were grown at 30 °C with shaking at 200 rpm until reaching near-saturation (approximately 3 days). This process of induction, staining, sorting, and growth was repeated for a total of 3 rounds.

### Supplementary Figures

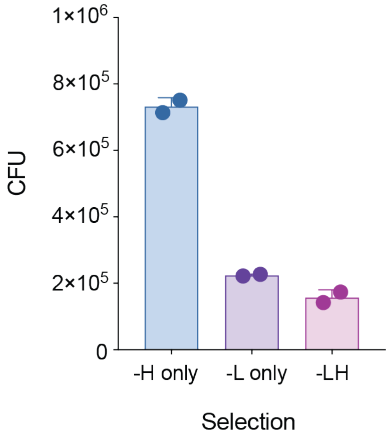


**Figure S1. Transfromation efficiency comparison based on auxotrophic markers.** Transformation efficiency is compared between single and dual selection conditions: His3-only (-H only), Leu2-only (-L only), and combined His3 and Leu2 (-LH) auxotrophic markers. Single-marker selection conditions (-H only and -L only) showed higher transformation efficiency compared to dual-marker selection (-LH), specifically, with the His3 marker (-H only) yielding a higher efficiency than the Leu2 marker (-L only).

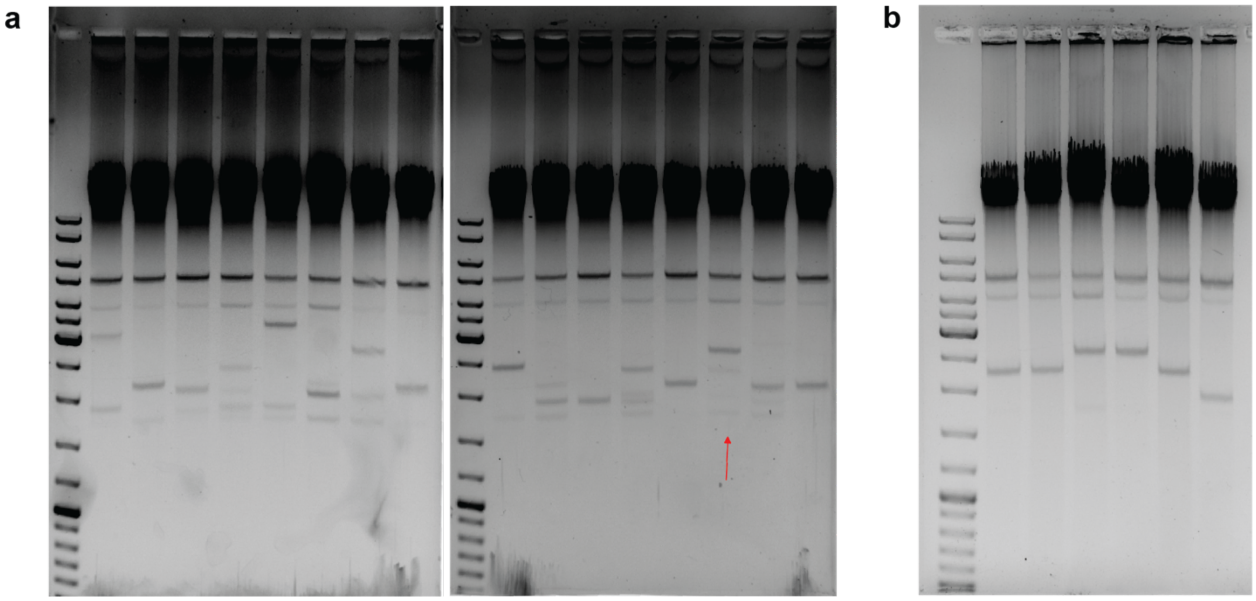


**Figure S2. Agarose gels of p1 minipreps after transformation with donor DNA mixtures. a**, The agarose gel of p1 minipreps post-integration. Agarose gel electrophoresis of p1 minipreps from cells outgrown after integrating a gene library with variable sequence lengths. On average, 2.75 distinct integrations per cell were achieved. **b**, Agarose gel of p1 minipreps from restreaked colonies. Six restreaked colonies are derived from one of the original colonies (indicated by a red arrow in panel **a**).

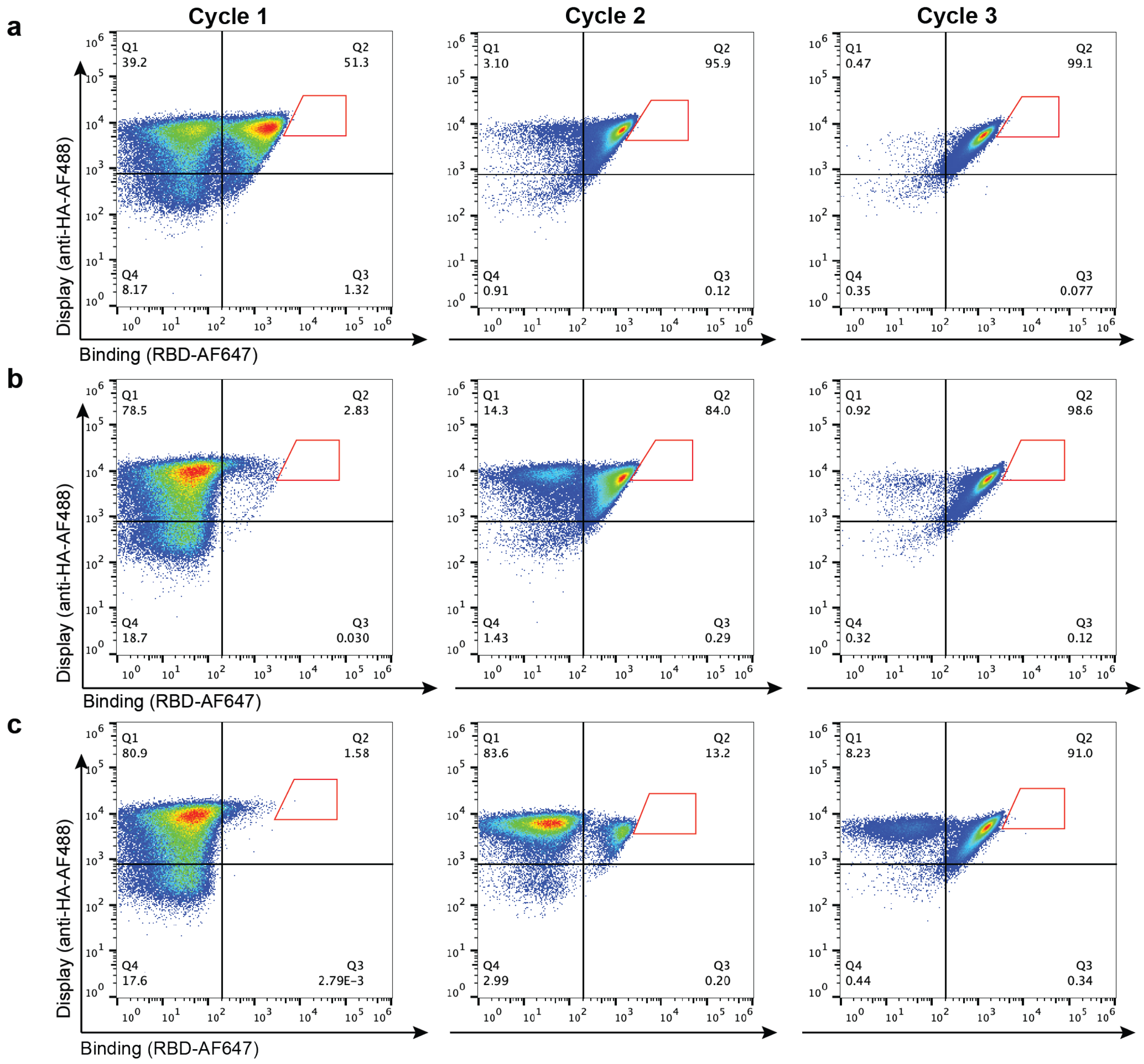


**Figure S3. FACS plots for the enrichment of an RBD-binding nanobody (RBD10i14) from three mock libraries.** Three mock libraries were successfully loaded onto OrthoRep, and the RBD10i14 nanobody was enriched by FACS. Red gates indicate cells sorted to seed the next cycle. **a**, 10% RBD10i14 library. **b**, 0.1% RBD10i14 library. **c**, 0.001% RBD10i14 library.

### Supplementary Tables

#### Table S1. Key plasmids used in this study.

| Name | Description | Sequence |
|---|---|---|
| pYY10 | 2*μ* plasmid for constitutive expression of TP901 under RNR2 promoter. The plasmid includes a kanMX marker for selection in yeast and propagation in *E. coli*. | GAAAATATTGTTGATGCGCTGGCAGTGTTCCTGCGCCGGTTGCATTCGATTCCTGTTTGTAATTGTCCTTTTAACAGCGA<br>TCGCGTATTTCGTCTCGCTCAGGCGCAATCACGAATGAATAACGGTTTGGTTGATGCGAGTGATTTTGATGACGAGCGT<br>AATGGCTGGCCTGTTGAACAAGTCTGGAAAGAAATGCATAAGCTTTTGCCATTCTCACCGGATTCAGTCGTCACTCATGG<br>TGATTTCTCACTTGATAACCTTATTTTTGACGAGGGGAAATTAATAGGTTGTATTGATGTTGGACGAGTCGGAATCGCAG<br>ACCGATACCAGGATCTTGCCATCCTATGGAACTGCCTCGGTGAGTTTTCTCCTTCATTACAGAAACGGCTTTTTCAAAAA<br>TATGGTATTGATAATCCTGATATGAATAAATTGCAGTTTCATTTGATGCTCGATGAGTTTTTCTAATCAGTACTGACAATAA<br>AAAGATTCTTGTTTTCAAGAACTTGTCATTTGTATAGTTTTTTTATATTGTAGTTGTTCTATTTTAATCAAATGTTAGCGTGA<br>TTTATATTTTTTTTCGCCTCGACATCATCTGCCCAGATGCGAAGTTAAGTGCGCAGAAAGTAATATCATGCGTCAATCGTA<br>TGTGAATGCTGGTCGCTATACTGCTGTCGATTCGATACTAACGCCGCCATCCAGTGTCGAAAACGAGCTCTCGAGAACC<br>CTTAATGTCGACAGTCGAACAAGAAGCAGGCAAAGTTTAGAGCACTGCCCCTCCGCACTCAAAAAAGAAAAAACTAGGA<br>GGAAAATAAAATTCTCAACCACACAAACACATAAACACATACAAATACAAATACAAGCTTATTTACTTGACATCGCGCGAT<br>CTTCCACTATTCAGCGCCGTCCGCCCTCTCTCGTGTTTTTTGTTTACGCGACAACTATGCGAAATCCGGAGCAACGGGC<br>AACCGTTTGGGGAAAGACCACACCCACGCGCGATCGCCATGGCAACGAGGTCGCACACGCCCCACACCCAGACCTCC<br>CTGCGAGCGGGCATGGGTACAATGTCCCCGTTGCCACAGACACCACTTCGTAGCACAGCGCAGAGCGTAGCGTGTTGT<br>TGCTGCTGACAAAAGAAAATTTTTCTTAGCAAAGCAAAGGAGGGGAAGCACGGGCAGATAGCACCGTACCATACCCTTG<br>GAAACTCGAAATGAACGAAGCAGGAAATGAGAGAATGAGAGTTTTGTAGGTATATATAGCGGTAGTGTTTGCGCGTTAC<br>CATCATCTTCTGGATCTATCTATTGTTCTTTTCCTCATCACTTTCCCCTTTTTCGCTCTTCTTCTTGTCTTTTATTTCTTTCT<br>TTTTTTTAATTGTTCCCTCGATTGGCTATCTACCAAAGAATCCAAACTTAATACACGTATTTATTTGTCCAATTACCATGAC<br>GAAAAAGGTCGCAATCTATACTCGTGTTTCCACAACAAATCAAGCCGAAGAAGGATTTAGCATAGATGAACAAATCGACC<br>GTCTGACAAAATATGCTGAGGCGATGGGGTGGCAGGTTAGCGACACGTACACTGACGCAGGATTCTCTGGAGCAAAAC<br>TAGAAAGACCTGCAATGCAGCGTCTTATTAATGACATAGAAAACAAGGCGTTCGATACAGTTCTAGTCTACAAATTGGAT<br>AGGCTATCTCGTTCTGTGAGAGACACACTATACCTAGTTAAAGATGTCTTTACCAAAAACAAGATAGACTTCATATCTCTG<br>AACGAAAGCATAGATACATCCAGCGCAATGGGATCATTGTTTCTTACAATTTTATCAGCTATAAACGAATTTGAAAGAGAA<br>AACATCAAGGAAAGGATGACGATGGGCAAGCTGGGCAGAGCCAAAAGCGGGAAGTCAATGATGTGGACAAAGACCGCT<br>TTCGGCTACTACCACAACAGAAAGACGGGGATCTTGGAGATAGTTCCTTTGCAGGCTACCATTGTAGAACAGATTTTTAC<br>GGATTACCTTTCCGGGATCTCCCTGACAAAACTTCGTGATAAATTGAACGAATCAGGCCATATCGGCAAAGATATACCCT<br>GGAGCTATAGAACTCTTCGTCAGACGCTGGATAATCCGGTTTATTGTGGGTATATAAAATTCAAAGACAGTTTGTTCGAA<br>GGTATGCACAAGCCCATCATTCCATATGAAACGTATCTGAAAGTGCAGAAAGAGCTGGAAGAAAGGCAGCAGCAGACCT<br>ACGAAAGGAATAACAATCCTAGACCCTTCCAAGCGAAGTATATGTTGAGCGGGATGGCTAGGTGTGGGTACTGCGGTG<br>CTCCTCTGAAAATAGTATTGGGGCATAAGAGGAAAGACGGCAGTAGGACAATGAAATATCATTGTGCCAATAGGTTCCC<br>GCGTAAAACAAAAGGGATTACCGTTTATAATGATAATAAGAAGTGCGACAGCGGCACATATGATTTAAGCAACCTGGAAA<br>ATACTGTGATCGACAACCTGATCGGTTTCCAAGAGAATAATGATTCCCTTTTAAAAATCATCAATGGTAATAACCAGCCAA<br>TACTAGACACAAGCTCATTCAAGAAGCAAATTTCCCAAATAGACAAGAAAATCCAAAAAAATAGCGACCTATACTTAAATG<br>ACTTTATCACAATGGATGAGTTAAAAGACAGAACTGATAGCTTACAGGCAGAAAAGAAATTACTGAAGGCGAAGATATCA<br>GAGAACAAGTTCAACGACTCAACTGACGTATTCGAACTAGTAAAAACCCAACTGGGCAGCATACCAATAAATGAACTGTC<br>ATACGACAACAAGAAGAAGATCGTGAACAATCTTGTAAGTAAGGTGGACGTGACAGCAGATAATGTTGATATAATCTTCA<br>AGTTCCAGTTAGCATAATAAATCCGCTCTAACCGAAAAGGAAGGAGTTAGACAACCTGAAGTCTAGGTCCCTATTTATTT<br>TTTTATAGTTATGTTAGTATTAAGAACGTTATTTATATTTCAAATTTTTCTTTTTTTTCTGTACAGACGCGTGTACGCATGTA<br>ACATTATACTGAAAACCTTGCTTGAGAAGGTTTTGGGACGCTCGAAGCCGCGGTGAGCAAAAGGCCAGCAAAAGGCCA<br>GGAACCGTAAAAAGGCCGCGTTGCTGGCGTTTTTCCATAGGCTCCGCCCCCCTGACGAGCATCACAAAAATCGACGCT<br>CAAGTCAGAGGTGGCGAAACCCGACAGGACTATAAAGATACCAGGCGTTTCCCCCTGGAAGCTCCCTCGTGCGCTCTC<br>CTGTTCCGACCCTGCCGCTTACCGGATACCTGTCCGCCTTTCTCCCTTCGGGAAGCGTGGCGCTTTCTCATAGCTCACG<br>CTGTAGGTATCTCAGTTCGGTGTAGGTCGTTCGCTCCAAGCTGGGCTGTGTGCACGAACCCCCCGTTCAGCCCGACCG<br>CTGCGCCTTATCCGGTAACTATCGTCTTGAGTCCAACCCGGTAAGACACGACTTATCGCCACTGGCAGCAGCCACTGGT<br>AACAGGATTAGCAGAGCGAGGTATGTAGGCGGTGCTACAGAGTTCTTGAAGTGGTGGCCTAACTACGGCTACACTAGA<br>AGGACAGTATTTGGTATCTGCGCTCTGCTGAAGCCAGTTACCTTCGGAAAAAGAGTTGGTAGCTCTTGATCCGGCAAAC<br>AAACCACCGCTGGTAGCGGTGGTTTTTTTGTTTGCAAGCAGCAGATTACGCGCAGAAAAAAAGGATCTCAAGAAGATCC<br>TTTGATCTTTTCTACACTAGTCGAAGCATCTGTGCTTCATTTTGTAGAACAAAAATGCAACGCGAGAGCGCTAATTTTTCA<br>AACAAAGAATCTGAGCTGCATTTTTACAGAACAGAAATGCAACGCGAAAGCGCTATTTTACCAACGAAGAATCTGTGCTT<br>CATTTTTGTAAAACAAAAATGCAACGCGAGAGCGCTAATTTTTCAAACAAAGAATCTGAGCTGCATTTTTACAGAACAGAA<br>ATGCAACGCGAGAGCGCTATTTTACCAACAAAGAATCTATACTTCTTTTTTGTTCTACAAAAATGCATCCCGAGAGCGCTA<br>TTTTTCTAACAAAGCATCTTAGATTACTTTTTTTCTCCTTTGTGCGCTCTATAATGCAGTCTCTTGATAACTTTTTGCACTGT<br>AGGTCCGTTAAGGTTAGAAGAAGGCTACTTTGGTGTCTATTTTCTCTTCCATAAAAAAAGCCTGACTCCACTTCCCGCGT<br>TTACTGATTACTAGCGAAGCTGCGGGTGCATTTTTTCAAGATAAAGGCATCCCCGATTATATTCTATACCGATGTGGATT<br>GCGCATACTTTGTGAACAGAAAGTGATAGCGTTGATGATTCTTCATTGGTCAGAAAATTATGAACGGTTTCTTCTATTTTG<br>TCTCTATATACTACGTATAGGAAATGTTTACATTTTCGTATTGTTTTCGATTCACTCTATGAATAGTTCTTACTACAATTTTT<br>TTGTCTAAAGAGTAATACTAGAGATAAACATAAAAAATGTAGAGGTCGAGTTTAGATGCAAGTTCAAGGAGCGAAAGGTG<br>GATGGGTAGGTTATATAGGGATATAGCACAGAGATATATAGCAAAGAGATACTTTTGAGCAATGTTTGTGGAAGCGGTAT<br>TCGCAATATTTTAGTAGCTCGTTACAGTCCGGTGCGTTTTTGGTTTTTTGAAAGTGCGTCTTCAGAGCGCTTTTGGTTTTC<br>AAAAGCGCTCTGAAGTTCCTATACTTTCTAGAGAATAGGAACTTCGGAATAGGAACTTCAAAGCGTTTCCGAAAACGAGC<br>GCTTCCGAAAATGCAACGCGAGCTGCGCACATACAGCTCACTGTTCACGTCGCACCTATATCTGCGTGTTGCCTGTATA<br>TATATATACATGAGAAGAACGGCATAGTGCGTGTTTATGCTTAAATGCGTATATGTGTTATGTATAGGTCTAGAGATCTGT<br>TTAGCTTGCCTCGTCCCCGCCGGGTCACCCGGCCAGCGACATGGAGGCCCAGAATACCCTCCTTGACAGTCTTGACGT<br>GCGCAGCTCAGGGGCATGATGTGACTGTCGCCCGTACATTTAGCCCATACATCCCCATGTATAATCATTTGCATCCATA<br>CATTTTGATGGCCGCACGGCGCGAAGCAAAAATTACGGCTCCTCGCTGCAGACCTGCGAGCAGGGAAACGCTCCCCTC<br>ACAGACGCGTTGAATTGTCCCCACGCCGCGCCCCTGTAGAGAAATATAAAAGGTTAGGATTTGCCACTGAGGTTCTTCT<br>TTCATATACTTCCTTTTAAAATCTTGCTAGGATACAGTTCTCACATCACATCCGAACATAAACAACCATGGGTAAGGAAAA<br>GACTCACGTTTCGAGGCCGCGATTAAATTCCAACATGGATGCTGATTTATATGGGTATAAATGGGCTCGCGATAATGTCG<br>GGCAATCAGGTGCGACAATCTATCGATTGTATGGGAAGCCCGATGCGCCAGAGTTGTTTCTGAAACATGGCAAAGGTAG<br>CGTTGCCAATGATGTTACAGATGAGATGGTCAGACTAAACTGGCTGACGGAATTTATGCCTCTTCCGACCATCAAGCATT<br>TTATCCGTACTCCTGATGATGCATGGTTACTCACCACTGCGATCCCCGGCAAAACAGCATTCCAGGTATTAGAAGAATAT<br>CCTGATTCAGGT |
| pYY12 | Plasmid for genertaion of donor DNA for TP901-mediated p1 integration in yeast. After linearization with ScaI, this plasmid produces | TATCAGCTCACTCAAAGGCGGTAATACGGTTATCCACAGAATCAGGGGATAACGCAGGAAAGAACATGTGAGCAAAAGG<br>CCAGCAAAAGGCCAGGAACCGTAAAAAGGCCGCGTTGCTGGCGTTTTTCCATAGGCTCCGCCCCCCTGACGAGCATCA<br>CAAAAATCGACGCTCAAGTCAGAGGTGGCGAAACCCGACAGGACTATAAAGATACCAGGCGTTTCCCCCTGGAAGCTC<br>CCTCGTGCGCTCTCCTGTTCCGACCCTGCCGCTTACCGGATACCTGTCCGCCTTTCTCCCTTCGGGAAGCGTGGCGCT<br>TTCTCATAGCTCACGCTGTAGGTATCTCAGTTCGGTGTAGGTCGTTCGCTCCAAGCTGGGCTGTGTGCACGAACCCCCC<br>GTTCAGCCCGACCGCTGCGCCTTATCCGGTAACTATCGTCTTGAGTCCAACCCGGTAAGACACGACTTATCGCCACTGG<br>CAGCAGCCACTGGTAACAGGATTAGCAGAGCGAGGTATGTAGGCGGTGCTACAGAGTTCTTGAAGTGGTGGCCTAACT<br>ACGGCTACACTAGAAGAACAGTATTTGGTATCTGCGCTCTGCTGAAGCCAGTTACCTTCGGAAAAAGAGTTGGTAGCTC<br>TTGATCCGGCAAACAAACCACCGCTGGTAGCGGTGGTTTTTTTGTTTGCAAGCAGCAGATTACGCGCAGAAAAAAAGGA<br>TCTCAAGAAGATCCTTTGATCTTTTCTACGGGGTCTGACGCTCAGTGGAACGAAAACTCACGTTAAGGGATTTTGGTCAT<br>GAGATTATCAAAAAGGATCTTCACCTAGATCCTTTTAAATTAAAAATGAAGTTTTAAATCAATCTAAAGTATATATGAGTAA<br>ACTTGGTCTGACAGTTACCAATGCTTAATCAGTGAGGCACCTATCTCAGCGATCTGTCTATTTCGTTCATCCATAGTTGC<br>CTGACTCCCCGTCGTGTAGATAACTACGATACGGGAGGGCTTACCATCTGGCCCCAGTGCTGCAATGATACCGCGAGA<br>CCCACGCTCACCGGCTCCAGATTTATCAGCAATAAACCAGCCAGCCGGAAGGGCCGAGCGCAGAAGTGGTCCTGCAAC<br>TTTATCCGCCTCCATCCAGTCTATTAATTGTTGCCGGGAAGCTAGAGTAAGTAGTTCGCCAGTTAATAGTTTGCGCAACG |

| | | |
|---|---|---|
| | donor DNA encoding the fluorescent protein mKate as a reporter with a *LEU2* auxotrophic selection marker, flanked by *attP* recombination sites for p1 integration. This plasmid contains AmpR marker for propagation in *E. coli.* | TTGTTGCCATTGCTACAGGCATCGTGGTGTCACGCTCGTCGTTTGGTATGGCTTCATTCAGCTCCGGTTCCCAACGATC<br>AAGGCGAGTTACATGATCCCCCATGTTGTGCAAAAAAGCGGTTAGCTCCTTCGGTCCTCCGATCGTTGTCAGAAGTAAG<br>TTGGCCGCAGTGTTATCACTCATGGTTATGGCAGCACTGCATAATTCTCTTACTGTCATGCCATCCGTAAGATGCTTTTC<br>TGTGACTGGTGAGTACTCAACCAAGTCATTCTGAGAATAGTGTATGCGGCGACCGAGTTGCTCTTGCCCGGCGTCAATA<br>CGGGATAATACCGCGCCACATAGCAGAACTTTAAAAGTGCTCATCATTGGAAAACGTTCTTCGGGGCGAAAACTCTCAA<br>GGATCTTACCGCTGTTGAGATCCAGTTCGATGTAACCCACTCGTGCACCCAACTGATCTTCAGCATCTTTTACTTTCACC<br>AGCGTTTCTGGGTGAGCAAAAACAGGAAGGCAAAATGCCGCAAAAAAGGGAATAAGGGCGACACGGAAATGTTGAATA<br>CTCATACTCTTCCTTTTTCAATATTATTGAAGCATTTATCAGGGTTATTGTCTCATGAGCGGATACATATTTGAATGTATTT<br>AGAAAAATAAACAAATAGGGGTTCCGCGCACATTTCCCCGAAAAGTGCCACCTGACGTCTAAGAAACCATTATTATCATG<br>ACATTAACCTATAAAAATAGGCGTATCACGAGGCCCTTTCGTCTCACGCGTTTCGGTGATGACGGTGAAAACCTCTGACA<br>CATGCAGCTCCCGGAGACGAGTACTATAATATATGAATTACATTATTAATTTAAAAGATACTAGATGGTGCAGCAGCAGC<br>AGCAGCAGTAGTAGGGCCAGTATTGGCGTTTCCCCCTCCTGACTTGAACAACCCTTTTAACGACTTTGAAATAGATAGAG<br>ACCTCTTCAAAGAGCTTTCTCTGGAAATACGGGGCTCAGAAGAGTCCACATTAGATGAAGCCATTATGGATTGCGAGTTT<br>TTATTTCGTTTATTCAAATTAAGGTAACTAAAAAACTCCTTTTTTAAGCAAGGATTTTCTTAACTTCTTCGGCGACAGCATC<br>ACCGACTTCGGTGGTACTGTTGGAACCACCTAAATCACCAGTTCTGATACCTGCATCCAAAACCTTTTTAACTGCATCTT<br>CAATGGCCTTACCTTCTTCAGGCAAGTTCAATGACAATTTCAACATCATTGCAGCAGACAAGATAGTGGCGATAGGGTTG<br>ACCTTATTCTTTGGCAAATCTGGAGCAGAACCGTGGCATGGTTCGTACAAACCAAATGCGGTGTTCTTGTCTGGCAAAG<br>AGGCCAAGGACGCAGATGGCAACAAACCCAAGGAACCTGGGATAACGGAGGCTTCATCGGAGATGATATCACCAAACA<br>TGTTGCTGGTGATTATAATACCATTTAGGTGGGTTGGGTTCTTAACTAGGATCATGGCGGCAGAATCAATCAATTGATGT<br>TGAACCTTCAATGTAGGGAATTCGTTCTTGATGGTTTCCTCCACAGTTTTTCTCCATAATCTTGAAGAGGCCAAAACATTA<br>GCTTTATCCAAGGACCAAATAGGCAATGGTGGCTCATGTTGTAGGGCCATGAAAGCGGCCATTCTTGTGATTCTTTGCA<br>CTTCTGGAACGGTGTATTGTTCACTATCCCAAGCGACACCATCACCATCGTCTTCCTTTCTCTTACCAAAGTAAATACCTC<br>CCACTAATTCTCTGACAACAACGAAGTCAGTACCTTTAGCAAATTGTGGCTTGATTGGAGATAAGTCTAAAAGAGAGTCG<br>GATGCAAAGTTACATGGTCTTAAGTTGGCGTACAATTGAAGTTCTTTACGGATTTTTAGTAAACCTTGTTCAGGTCTAACA<br>CTACCGGTACCCCATTTAGGACCACCCACAGCACCTAACAAAACGGCATCAGCCTTCTTGGAGGCTTCCAGCGCCTCAT<br>CTGGAAGTGGAACACCTGTAGCATCGATAGCAGCACCACCAATTAAATGATTTTCGAAATCGAACTTGACATTGGAACGA<br>ACATCAGAAATAGCTTTAAGAACCTTAATGGCTTCGGCTGTGATTTCTTGACCAACGTGGTCACCTGGCAAAACGACGAT<br>CTTCTTAGGAACGTGCATTTATAATCATAAACTATTAAATCATATTAAGAATCATTCAATTGATTTTTGGGATTTGGATTATA<br>TAGAAAAGTATGACCTAATTGACTCCGGCGAAAAAGCATGCTTATCTGTGCCCCAGTTTGCTAGGGAGGTCGCAGTATC<br>TGGCCACAGCCACCTCGTGCTGCTCGACGTAGGTCTCTTTGTCGGCCTCCTTGATTCTTTCCAGTCTTCTGTCCACATA<br>GTAGACGCCGGGCATCTTGAGGTTCTTAGCGGGTTTCTTGGATCTGTATGTGGTCTTCAAGTTGCAGATCAGGTGGCCC<br>CCGCCCACGAGCTTCAGGGCCATGTCGGCTCTGCCTTCCAGGCCGCCGTCAGCGGGGTACAGGGTCTCGGTGGAGGC<br>CTCCCAGCCGAGTGTTTTCTTCTGCATCACAGGGCCGTTGGATGGGAAGTTCACCCCTCTGATCTTGACGTTGTAGATG<br>AGGCAGCCGTCCTGGAGGCTGGTGTCCTGGGTAGCGGTCAGCACGCCCCCGTCTTCGTATGTGGTGACTCTCTCCCAT<br>GTGAAGCCCTCGGGGAAGGACTGCTTAAAGAAGTCGGGGATGCCCTGGGTGTGGTTGATGAAGGTTTTGCTGCCGTAC<br>ATGAAGCTGGTAGCCAGGATGTCGAAGGCGAAGGGGAGAGGGCCGCCCTCGACCGCCTTGATTCTCATGGTCTGGGT<br>GCCCTCGTAGGGCTTGCCTTCGCCCTCGGATGTGCACTTGAAGTGGTGGTTGTTCACGGTGCCCTCCATGTACAGCTT<br>CATGTGCATGTTCTCCTTAATCAGCTCGCTCACATGCATTTTACATGTCTATGAGCTTATCACATGTTTCTACATTTTTCTG<br>ATATTTGAAAGTTATTAATCTTTTTGTTTCTATAGATCTTCCTATGTATAGGTCATCAAAAGGAGTTTTTTAGTTACCTTAAT<br>TGAAATAAACGAAATAAAAACTCGCCCTCAGACTCAGATTGGTGTTTCAAAGATCCTCAAGGCAATAGAATAACAGATTTT<br>GATAGTATTAATAAAGAATTAGGTCTTGGTAGAAGAGATGTAAAATTAGATAAAGGTCATGATGATTTAATTAAATTATGTA<br>CTGAAAAAATAGATAGTACTATTAATTGCGTTGCGCTCACTGCCCGCTTTCCAGTCGGGAAACCTGTCGTGCCAGCTGC<br>ATTAATGAATCGGCCAACGCGCGGGGAGAGGCGGTTTGCGTATTGGGCGCTCTTCCGCTTCCTCGCTCACTGACTCGC<br>TGCGCTCGGTCGTTCGGCTGCGGCGAGCGG |
| pYY13 | Plamids for generation of donor DNA for construction of a TP901-compatible landing pad p1. After linearization with ScaI, this plasmid produces donor DNA with the *URA3* selectable marker, *attB* recombination sites, and homology flanks matches the original landing pad p1 used in the lab[1]. This plasmid contains AmpR marker for propagation in *E. coli.* | CAGTAAGAGAATTATGCAGTGCTGCCATAACCATGAGTGATAACACTGCGGCCAACTTACTTCTGACAACGATCGGAGG<br>ACCGAAGGAGCTAACCGCTTTTTTGCACAACATGGGGGATCATGTAACTCGCCTTGATCGTTGGGAACCGGAGCTGAAT<br>GAAGCCATACCAAACGACGAGCGTGACACCACGATGCCTGTAGCAATGGCAACAACGTTGCGCAAACTATTAACTGGC<br>GAACTACTTACTCTAGCTTCCCGGCAACAATTAATAGACTGGATGGAGGCGGATAAAGTTGCAGGACCACTTCTGCGCT<br>CGGCCCTTCCGGCTGGCTGGTTTATTGCTGATAAATCTGGAGCCGGTGAGCGTGGGTCTCGCGGTATCATTGCAGCAC<br>TGGGGCCAGATGGTAAGCCCTCCCGTATCGTAGTTATCTACACGACGGGGAGTCAGGCAACTATGGATGAACGAAATA<br>GACAGATCGCTGAGATAGGTGCCTCACTGATTAAGCATTGGTAACTGTCAGACCAAGTTTACTCATATATACTTTAGATT<br>GATTTAAAACTTCATTTTTAATTTAAAAGGATCTAGGTGAAGATCCTTTTTGATAATCTCATGACCAAAATCCCTTAACGTG<br>AGTTTTCGTTCCACTGAGCGTCAGACCCCGTAGAAAAGATCAAAGGATCTTCTTGAGATCCTTTTTTTCTGCGCGTAATC<br>TGCTGCTTGCAAACAAAAAAACCACCGCTACCAGCGGTGGTTTGTTTGCCGGATCAAGAGCTACCAACTCTTTTTCCGAA<br>GGTAACTGGCTTCAGCAGAGCGCAGATACCAAATACTGTTCTTCTAGTGTAGCCGTAGTTAGGCCACCACTTCAAGAAC<br>TCTGTAGCACCGCCTACATACCTCGCTCTGCTAATCCTGTTACCAGTGGCTGCTGCCAGTGGCGATAAGTCGTGTCTTA<br>CCGGGTTGGACTCAAGACGATAGTTACCGGATAAGGCGCAGCGGTCGGGCTGAACGGGGGGTTCGTGCACACAGCCC<br>AGCTTGGAGCGAACGACCTACACCGAACTGAGATACCTACAGCGTGAGCTATGAGAAAGCGCCACGCTTCCCGAAGGG<br>AGAAAGGCGGACAGGTATCCGGTAAGCGGCAGGGTCGGAACAGGAGAGCGCACGAGGGAGCTTCCAGGGGGAAACG<br>CCTGGTATCTTTATAGTCCTGTCGGGTTTCGCCACCTCTGACTTGAGCGTCGATTTTTGTGATGCTCGTCAGGGGGGCG<br>GAGCCTATGGAAAAACGCCAGCAACGCGGCCTTTTTACGGTTCCTGGCCTTTTGCTGGCCTTTTGCTCACATGTTCTTTC<br>CTGCGTTATCCCCTGATTCTGTGGATAACCGTATTACCGCCTTTGAGTGAGCTGATACCGCTCGCCGCAGCCGAACGAC<br>CGAGCGCAGCGAGTCAGTGAGCGAGGAAGCGGAAGAGCGCCCAATACGCAAACCGCCTCTCCCCGCGCGTTGGCCG<br>ATTCATTAATGCAGCTGGCACGACAGGTTTCCCGACTGGAAAGCGGGCAGTGAGCGCAACGCAATTAATAGTACTATCT<br>ATTTTTTCAGTACATAATTTAATTAAATCATCATGACCTTTATCTAATTTTACATCTCTTCTACCAAGACCTAATTCTTTATTA<br>ATACTATCAAAATCTGTTATTCTATTGCCTTGAGGATCTTTGAAACACCAATCTGAGTCTGAGGGCATGCAACACAATTAA<br>CATCTCAATCAAGGTAAATGCTTTTTTTTCGCCGGAGTCAATTAGGTCATACTTTTCTATATAATCCAAATCCCAAAAATCA<br>ATTGAATGATTCTTAATATGATTTAATAGTTTATGATTATAAATGTCGAAAGCTACATATAAGGAACGTGCTGCTACTCATC<br>CTAGTCCTGTTGCTGCCAAGCTATTTAATATCATGCACGAAAAGCAAACAAACTTGTGTGCTTCATTGGATGTTCGTACC<br>ACCAAGGAATTACTGGAGTTAGTTGAAGCATTAGGTCCCAAAATTTGTTTACTAAAAACACATGTGGATATCTTGACTGAT<br>TTTTCCATGGAGGGCACAGTTAAGCCGCTAAAGGCATTATCCGCCAAGTACAATTTTTTACTCTTCGAGGACAGAAAATT<br>TGCTGACATTGGTAATACAGTCAAATTGCAGTACTCTGCGGGTGTATACAGAATAGCAGAATGGGCAGACATTACGAAT<br>GCACACGGTGTGGTGGGCCCAGGTATTGTTAGCGGTTTGAAGCAGGCGGCAGAAGAAGTAACAAAGGAACCTAGAGG<br>CCTTTTGATGTTAGCAGAATTGTCATGCAAGGGCTCCCTATCTACTGGAGAATATACTAAGGGTACTGTTGACATTGCGA<br>AGAGCGACAAAGATTTTGTTATCGGCTTTATTGCTCAAAGAGACATGGGTGGAAGAGATGAAGGTTACGATTGGTTGATT<br>ATGACACCCGGTGTGGGTTTAGATGACAAGGGAGATGCATTGGGTCAACAGTATAGAACCGTGGATGATGTGGTTTCTA<br>CAGGATCTGACATTATTATTGTTGGAAGAGGACTATTTGCAAAGGGAAGGGATGCTAAGGTAGAGGGTGAACGTTACAG<br>AAAAGCAGGCTGGGAAGCATATTTGAGAAGATGCGGCCAGCAAAACTAAAAAGCATTTACCTTGATTTGGATGTTAATTG<br>TGTTAATCCATAATGGCTTCATCTAATGTGGACTCTTCTGAGCCCCGTATTTCCAGAGAAAGCTCTTTGAAGAGGTCTCT<br>ATCTATTTCAAAGTCGTTAAAAGGGTTGTTCAAGTCAGGAGGGGGAAACGCCAATACTGGCCCTACTACTGCTGCTGCT<br>GCTGCTGCACCATCTAGTATCTTTTAAATTAATAATGTAATTCATATATTATAGTACTCGTCTCCGGGAGCTGCATGTGTC<br>AGAGGTTTTCACCGTCATCACCGAAACGCGTGAGACGAAAGGGCCTCGTGATACGCCTATTTTTATAGGTTAATGTCAT<br>GATAATAATGGTTTCTTAGACGTCAGGTGGCACTTTTCGGGGAAATGTGCGCGGAACCCCTATTTGTTTATTTTTCTAAAT<br>ACATTCAAATATGTATCCGCTCATGAGACAATAACCCTGATAAATGCTTCAATAATATTGAAAAAGGAAGAGTATGAGTAT<br>TCAACATTTCCGTGTCGCCCTTATTCCCTTTTTTGCGGCATTTTGCCTTCCTGTTTTTGCTCACCCAGAAACGCTGGTGAA<br>AGTAAAAGATGCTGAAGATCAGTTGGGTGCACGAGTGGGTTACATCGAACTGGATCTCAACAGCGGTAAGATCCTTGAG<br>AGTTTTCGCCCCGAAGAACGTTTTCCAATGATGAGCACTTTTAAAGTTCTGCTATGTGGCGCGGTATTATCCCGTATTGA<br>CGCCGGGCAAGAGCAACTCGGTCGCCGCATACACTATTCTCAGAATGACTTGGTTGAGTACTCACCAGTCACAGAAAAG<br>CATCTTACGGATGGCATGA |
| pYY33 | Plamids for generation of donor DNA for construction of a | CAGTAAGAGAATTATGCAGTGCTGCCATAACCATGAGTGATAACACTGCGGCCAACTTACTTCTGACAACGATCGGAGG<br>ACCGAAGGAGCTAACCGCTTTTTTGCACAACATGGGGGATCATGTAACTCGCCTTGATCGTTGGGAACCGGAGCTGAAT<br>GAAGCCATACCAAACGACGAGCGTGACACCACGATGCCTGTAGCAATGGCAACAACGTTGCGCAAACTATTAACTGGC<br>GAACTACTTACTCTAGCTTCCCGGCAACAATTAATAGACTGGATGGAGGCGGATAAAGTTGCAGGACCACTTCTGCGCT<br>CGGCCCTTCCGGCTGGCTGGTTTATTGCTGATAAATCTGGAGCCGGTGAGCGTGGGTCTCGCGGTATCATTGCAGCAC<br>TGGGGCCAGATGGTAAGCCCTCCCGTATCGTAGTTATCTACACGACGGGGAGTCAGGCAACTATGGATGAACGAAATA<br>GACAGATCGCTGAGATAGGTGCCTCACTGATTAAGCATTGGTAACTGTCAGACCAAGTTTACTCATATATACTTTAGATT<br>GATTTAAAACTTCATTTTTAATTTAAAAGGATCTAGGTGAAGATCCTTTTTGATAATCTCATGACCAAAATCCCTTAACGTG |

| | | |
|---|---|---|
| | TP901-compatible landing pad p1. After linearization with ScaI, this plasmid produces donor DNA with the *TRP1* selectable marker, *attB* recombination sites, and homology flanks matches the original landing pad p1 used in the lab[1]. This plasmid contains AmpR marker for propagation in *E. coli*.. | AGTTTTCGTTCCACTGAGCGTCAGACCCCGTAGAAAAGATCAAAGGATCTTCTTGAGATCCTTTTTTTCTGCGCGTAATC<br>TGCTGCTTGCAAACAAAAAAACCACCGCTACCAGCGGTGGTTTGTTTGCCGGATCAAGAGCTACCAACTCTTTTTCCGAA<br>GGTAACTGGCTTCAGCAGAGCGCAGATACCAAATACTGTTCTTCTAGTGTAGCCGTAGTTAGGCCACCACTTCAAGAAC<br>TCTGTAGCACCGCCTACATACCTCGCTCTGCTAATCCTGTTACCAGTGGCTGCTGCCAGTGGCGATAAGTCGTGTCTTA<br>CCGGGTTGGACTCAAGACGATAGTTACCGGATAAGGCGCAGCGGTCGGGCTGAACGGGGGGTTCGTGCACACAGCCC<br>AGCTTGGAGCGAACGACCTACACCGAACTGAGATACCTACAGCGTGAGCTATGAGAAAGCGCCACGCTTCCCGAAGGG<br>AGAAAGGCGGACAGGTATCCGGTAAGCGGCAGGGTCGGAACAGGAGAGCGCACGAGGGAGCTTCCAGGGGGAAACG<br>CCTGGTATCTTTATAGTCCTGTCGGGTTTCGCCACCTCTGACTTGAGCGTCGATTTTTGTGATGCTCGTCAGGGGGGCG<br>GAGCCTATGGAAAAACGCCAGCAACGCGGCCTTTTTACGGTTCCTGGCCTTTTGCTGGCCTTTTGCTCACATGTTCTTTC<br>CTGCGTTATCCCCTGATTCTGTGGATAACCGTATTACCGCCTTTGAGTGAGCTGATACCGCTCGCCGCAGCCGAACGAC<br>CGAGCGCAGCGAGTCAGTGAGCGAGGAAGCGGAAGAGCGCCCAATACGCAAACCGCCTCTCCCCGCGCGTTGGCCG<br>ATTCATTAATGCAGCTGGCACGACAGGTTTCCCGACTGGAAAGCGGGCAGTGAGCGCAACGCAATTAATAGTACTATCT<br>ATTTTTTCAGTACATAATTTAATTAAATCATCATGACCTTTATCTAATTTTACATCTCTTCTACCAAGACCTAATTCTTTATTA<br>ATACTATCAAAATCTGTTATTCTATTGCCTTGAGGATCTTTGAAACACCAATCTGAGTCTGAGGGCATGCAACACAATTAA<br>CATCTCAATCAAGGTAAATGCTTTTATATGAAAGTTTTTATAATAATTATAAAATGTCTGTTATTAATTTCACAGGTAGTTCT<br>GGTCCATTGGTGAAAGTTTGCGGCTTGCAGAGCACAGAGGCCGCAGAATGTGCTCTAGATTCCGATGCTGACTTGCTG<br>GGTATTATATGTGTGCCCAATAGAAAGAGAACAATTGACCCGGTTATTGCAAGGAAAATTTCAAGTCTTGTAAAAGCATAT<br>AAAAATAGTTCAGGCACTCCGAAATACTTGGTTGGCGTGTTTCGTAATCAACCTAAGGAGGATGTTTTGGCTCTGGTCAA<br>TGATTACGGCATTGATATCGTCCAACTGCATGGAGATGAGTCGTGGCAAGAATACCAAGAGTTCCTCGGTTTGCCAGTT<br>ATTAAAAGACTCGTATTTCCAAAAGACTGCAACATACTACTCAGTGCAGCTTCACAGAAACCTCATTCGTTTATTCCCTTG<br>TTTGATTCAGAAGCAGGTGGGACAGGTGAACTTTTGGATTGGAACTCGATTTCTGACTGGGTTGGAAGGCAAGAGAGCC<br>CCGAAAGCTTACATTTTATGTTAGCTGGTGGACTGACGCCAGAAAATGTTGGTGATGCGCTTAGATTAAATGGCGTTATT<br>GGTGTTGATGTAAGCGGAGGTGTGGAGACAAATGGTGTAAAAGACTCTAACAAAATAGCAAATTTCGTCAAAAATGCTAA<br>GAAATGAAAATCAGCAAATGAAGATATCAAATATGAAATAAAACAACAAGGAAACGTTTAAAACTCTAGAAATCTGATATC<br>AAAATGATCGTTAAACCTATCGCTTCCCTAAGTGTTGTCAACGTACGAAAGGGTGCCTTGCTTATATGTTTTTAAAAAAC<br>CCTCTTTGATATGATATTTTACCCTTTGATTTAGAAATCGCTTATTGATCTTATTAGATAAAATAAACGCAAAACACTTGTT<br>GATTAAGCAGAATAATGCCAGAATATCAGAATTGCAACTATCGAAACTGCAATCGCTAGTGTAAATCCTAGTATTTTCCTT<br>CCCTTTCTCTAAGTAAGATGGGTTTTCCCGGAAAATACGATATTGTTTTCCCTTTAAAAGTTTAGAACAGAAGCAAACTAA<br>AAGCGAGAACTTGAAATCAGACCCAAAAAAACACGTTTAAGTTTATGCACAACCTAATAGCATCAACCAGCGATGCTAAT<br>CTAGCCAAAAAACGTAATGCAGCAACTGAACTGGTAGACACAAGCCTCTAAACTGTTTCAAGTTGTTGTTATATATGGGA<br>AAAAAACCTTTAAAAATTTGTTTAAATGATCTTCTTTGAAAAATAAGGCTTGTAATGTTGCAATTACGTATTGTGAGGCTGC<br>AATTTATAGATCAGGGAAAAGGAAAAGGAAAGGGTTCTTCCATGCGTCAGCGAGGGGCCCAGGGGCTGGGCAGTGGCT<br>GAACAAAGCTCGTATGAGTTGAGAAGGAATGCTTGTGGGAATATTGAATATGTGTATATGTGTTGTATATGTGAGAGAAT<br>TGTGTAAAGTGGGCTATCAGTAGTTGGATCATTTGGTTATAGCGATACGATCTGATGCACGAAGTCTTTTATGGGCCCTA<br>TCGAACTTAGGATAACCATTATTGCACCGTATAAGACGAAGGAAAGGGAGCTGATAGCAGCAGCTCTTCTATCTACTGTT<br>AAGAGGTAGCTGGTGGCGATGGCACCGCAGTTACAGCCTAGAGTTAGGCCTCTGGTGACGAAATCATCGCGAGGTACT<br>CTGAAGAAGTCAAGCATGGGACCTCCGGATAGTGCACCTACGATACCACTTAGCACTGTAATGACAGCCATTACGGTAA<br>CGGATCCTTCGAAATTTTCGATGACGGGCATCCCCAAGGCTAAAGTGATGCTTCTTCCAATGAATCCAATGGAGTTATAT<br>GAGCTGATACCGATGTTGTAGCATATAAAGGGATTTAACATTAGAGTGGCAGCACATAAAGAGATGGTTGGGATAAGAA<br>GCACGACAAAATGTTTCTTTAGATCGCTTCTGTAAGTGTACATGGGCATCGATAATGCGACAATAGAGACGTCCATACAG<br>GAGGAAAAAAATATCACCGGCGCCAGGCCAGTTCTTGTTTGGACCTGATACACTAGTATTTGTGAATAATGTTAAGTATGT<br>TGTACCAACCTTGTATTCACGAAGATCCGAGAGGAAATATTTGATTTGGCGATGTTTAATGAGTGATGAGATCAATTCAA<br>CGATCCAGAATAGAGCCACTGAACAAATAACAGGATGCAAGAATTTCTTATACTTAGGATTTGGCAATAATGGTGCATCG<br>GTGACGATGAAGAACATGCAAATGGCGGTAAATAGTTGGAAAGGCATGATGTACCATGAAAAATAGTATGTGAATATAGT<br>TGCGAAAAACCCCAATCCGTATAGAACATGATGCAAGTGATCGTGCCACATGTTTACGGTGAAAAGATGTTCCATTCTGT<br>GAGAGAACTGCCTGGTGATGGCTCTTTCACACTCTACCTCTCTATTTCTATCAGAAAATTGATCATGAGAGGTAGATCCA<br>CAGAAACGTTTCGATGCCATTGGAACCATGGCTGCATTATTACGAGAATGTATCAGAGCAGGCTGTAATAACAACTGATT<br>TGTTTCAGCAGTTGAGTCAGATCCATGAGTGTTATCCTCGTCATAAATGTCCTGCACATTTTGAGTGATCCCTAATCCCTT<br>GTCGTTAGGAAGTCCAATTTCACTACCTCTTAGAATGTCAGCATCTAAATTAGCAGTATCACCCATTGCCAAATGATGAGT<br>AAGATTTGTACGAGTTCTTTGCAATGGGAACCCACTATTTGAAGCAACGTTGACTAATTGGAAATCTTCAACATCAGGAT<br>CCGAGAAAATGTCAGCAAAACTATGAATATCATCATCGTCTTCGTTTATGTCCAGTGTACTTTCTTCTTCAGTAGCATCTT<br>CATCCGCAGAACTGTATCCATTCTCAGCATCACGAAGAATATCCTCTTTACTGCCGAGTAAAGAAGTCCGTTTCCTTCCA<br>TTCGACCTACCAAGGTGGCCAGTAGCCAGTGCAAACAGCTTTTGACCAAATATAGTTACGTATGCCAAACAAAACCCGG<br>TGATTATGTACATGAACACGAACGAGGCAGCTATTAACATAGCTTCTCTAAAAGATATCCAGGGACTCAAAGGTAAAGTG<br>ACAAACGCTGGTGTAAAGAATAAATTCATCCATCTCAAAGCCCAACTTAAAGGAACATCGATGACACCAACAATCTTCTG<br>AGTTAAGTTCTCATTAATCATAGATAAAACACATAAGAAACAGAAATTTACCAACATTACAGCAACTGATGCAGGAAATTT<br>GATCCCTATCACATCTTTAATTAATATATTAACTCCATATAAGATCATCATGATCAATATCACACCGATTGGAACGAAAACG<br>TACTGTTTCAACAAACTGTACTTATAAATATGCAAAAAATCAACGAATGATCCGGTATGAAAGCATTTACCTTGATTTGGA<br>TGTTAATTGTGTTAATCCATAATGGCTTCATCTAATGTGGACTCTTCTGAGCCCCGTATTTCCAGAGAAAGCTCTTTGAAG<br>AGGTCTCTATCTATTTCAAAGTCGTTAAAAGGGTTGTTCAAGTCAGGAGGGGGAAACGCCAATACTGGCCCTACTACTG<br>CTGCTGCTGCTGCTGCACCATCTAGTATCTTTTAAATTAATAATGTAATTCATATATTATAGTACTCGTCTCCGGGAGCTG<br>CATGTGTCAGAGGTTTTCACCGTCATCACCGAAACGCGTGAGACGAAAGGGCCTCGTGATACGCCTATTTTTATAGGTT<br>AATGTCATGATAATAATGGTTTCTTAGACGTCAGGTGGCACTTTTCGGGGAAATGTGCGCGGAACCCCTATTTGTTTATT<br>TTTCTAAATACATTCAAATATGTATCCGCTCATGAGACAATAACCCTGATAAATGCTTCAATAATATTGAAAAAGGAAGAG<br>TATGAGTATTCAACATTTCCGTGTCGCCCTTATTCCCTTTTTTGCGGCATTTTGCCTTCCTGTTTTTGCTCACCCAGAAAC<br>GCTGGTGAAAGTAAAAGATGCTGAAGATCAGTTGGGTGCACGAGTGGGTTACATCGAACTGGATCTCAACAGCGGTAA<br>GATCCTTGAGAGTTTTCGCCCCGAAGAACGTTTTCCAATGATGAGCACTTTTAAAGTTCTGCTATGTGGCGCGGTATTAT<br>CCCGTATTGACGCCGGGCAAGAGCAACTCGGTCGCCGCATACACTATTCTCAGAATGACTTGGTTGAGTACTCACCAGT<br>CACAGAAAAGCATCTTACGGATGGCATGA |
| pYY62 | Plasmid used for constructing donor DNA for TP901-compatible p1 integration. This plasmid is a sfGFP dropout plasmid where sfGFP can be replaced by GOIs using Golden Gate Assembly. This allows the p1 donor to be customized to encode a GOI-HA- | CACATTTCCCCGAAAAGTGCCACCTGACGTCTAAGAAACCATTATTATCATGACATTAACCTATAAAAATAGGCGTATCAC<br>GAGGCCCTTTCGTCTCACGCGTTTCGGTGATGACGGTGAAAACCTCTGACACATGCAGCTCCCGGAGACGAGTACTATA<br>ATATATGAATTACATTATTAATTTAAAAGATACTAGATGGTGCAGCAGCAGCAGCAGCAGTAGTAGGGCCAGTATTGGCG<br>TTTCCCCCTCCTGACTTGAACAACCCTTTTAACGACTTTGAAATAGATAGTGACCTCTTCAAAGAGCTTTCTCTGGAAATA<br>CGGGGCTCAGAAGAGTCCACATTAGATGAAGCCATTATGGATTGCGAGTTTTTATTTCGTTTATTCAAATTAAGGTAACTA<br>AAAAACTCCTTTTGATGACCTATACATAGGAAGATCTATAGAAACAAAAAGATTAATAACTTTCAAATATCAGAAAAATATA<br>GAAACATGTGATAAGCTCATAGACATATAAAAATATGAGATTCCCATCTATCTTCACCGCTGTTTTGTTCGCTGCTTCTTC<br>TGCTTTGGCTGCTCCAGCTAACACCACCACCGAAGACGAAACCGCTCAAATCCCAGCTGAAGCTGTTATCGACTACTCT<br>GACTTGGAAGGTGACTTCGACGCTGCTGCTTTGCCATTGTCTAACTCTACCAACAACGGTTTGTCTTCTACCAACACCAC<br>CATCGCTTCTATCGCTGCTAAGGAAGAAGGTGTTCAATTGGACAAGAGAGAAGCTGACGCAGAGACCGAAAGTGAAAC<br>GTGATTTCATGCGTCATTTTGAACATTTTGTAAATCTTATTTAATAATGTGTGCGGCAATTCACATTTAATTTATGAATGTTT<br>TCTTAACATCGCGGCAACTCAAGAAACGGCAGGTTCGGATCTTAGCTACTAGAGAAAGAGGAGAAATACTAGATGCGTA<br>AAGGCGAAGAGCTGTTCACTGGTGTCGTCCCTATTCTGGTGGAACTGGATGGTGATGTCAACGGTCATAAGTTTTCCGT<br>GCGTGGCGAGGGTGAAGGTGACGCAACTAATGGTAAACTGACGCTGAAGTTCATCTGTACTACTGGTAAACTGCCGGTT<br>CCTTGGCCGACTCTGGTAACGACGCTGACTTATGGTGTTCAGTGCTTTGCTCGTTATCCGGACCATATGAAGCAGCATG<br>ACTTCTTCAAGTCCGCCATGCCGGAAGGCTATGTGCAGGAACGCACGATTTCCTTTAAGGATGACGGCACGTACAAAAC<br>GCGTGCGGAAGTGAAATTTGAAGGCGATACCCTGGTAAACCGCATTGAGCTGAAAGGCATTGACTTTAAAGAGGACGG<br>CAATATCCTGGGCCATAAGCTGGAATACAATTTTAACAGCCACAATGTTTACATCACCGCCGATAAACAAAAAAATGGCA<br>TTAAAGCGAATTTTAAAATTCGCCACAACGTGGAGGATGGCAGCGTGCAGCTGGCTGATCACTACCAGCAAAACACTCC<br>AATCGGTGATGGTCCTGTTCTGCTGCCAGACAATCACTATCTGAGCACGCAAAGCGTTCTGTCTAAAGATCCGAACGAG<br>AAACGCGATCATATGGTTCTGCTGGAGTTCGTAACCGCAGCGGGCATCACGCATGGTATGGATGAACTGTACAAATGAC<br>CAGGCATCAAATAAAACGAAAGGCTCAGTCGAAAGACTGGGCCTTTCGTTTTATCTGTTGTTTGTCGGTGAACGCTCTCT<br>ACTAGAGTCACACTGGCTCACCTTCGGGTGGGCCTTTCTGCGTTTATAGGTCTCGGTTCCAAGGACAATAGCTCGACGA<br>TTGAAGGTAGATACCCATACGACGTTCCAGACTACGCTCTGCAGGCTAGTGGTGGTGGTGGTTCTGGTGGTGGTGGTT<br>CTGGTGGTGGTGGTTCTGCTAGCGACGTCCAGGAACTGACAACTATATGCGAGCAAATCCCCTCACCAACTTTAGAATC<br>GACGCCGTACTCTTTGTCAACGACTACTATTTTGGCCAACGGGAAGGCAATGCAAGGAGTTTTTGAATATTACAAATCAG<br>TAACGTTTGTCAGTAATTGCGGTTCTCACCCCTCAACAACTAGCAAAGGCAGCCCCATAAACACACAGTATGTTTTTTGA<br>ATCCTAATAATGGCCGCAAAAAAAAAAAAAAAAAAAAAAAAAAAAAAAAAAAAAAAAAAAAAAAAAAAAAAAAAAAAAAA |

| | | |
|---|---|---|
| | Aga2 fusion protein with a LEU2 auxotrophic selection marker, flanked by attP recombination sites. The donor DNA for p1 integration will be generated by linearization with ScaI. This plasmid contains AmpR marker for propagation in *E. coli.* | AAAAAAAAAACCTGTCACCGGATGTGTTTTCCGGTCTGATGAGTCCGTGAGGACGAAACAGGACTAGGCTGCGAATTTC<br>TTATGATTTATGATTTTTATTATTAAATAAGTTATAAAAAAAATAAGTGTATACAAATTTTAAAGTGACTCTTAGGTTTTAAAA<br>CGAAAATTCTTATTCTTGAGTAACTCTTTCCTGTAGGTCAGGTTGCTTTCTCAGGTATAGCATGAGGTCGCTCTTAAGCAA<br>GGATTTTCTTAACTTCTTCGGCGACAGCATCACCGACTTCGGTGGTACTGTTGGAACCACCTAAATCACCAGTTCTGATA<br>CCTGCATCCAAAACCTTTTTAACTGCATCTTCAATGGCCTTACCTTCTTCAGGCAAGTTCAATGACAATTTCAACATCATT<br>GCAGCAGACAAGATAGTGGCGATAGGGTTGACCTTATTCTTTGGCAAATCTGGAGCAGAACCGTGGCATGGTTCGTACA<br>AACCAAATGCGGTGTTCTTGTCTGGCAAAGAGGCCAAGGACGCAGATGGCAACAAACCCAAGGAACCTGGGATAACGG<br>AGGCTTCATCGGAGATGATATCACCAAACATGTTGCTGGTGATTATAATACCATTTAGGTGGGTTGGGTTCTTAACTAGG<br>ATCATGGCGGCAGAATCAATCAATTGATGTTGAACCTTCAATGTAGGAAATTCGTTCTTGATGGTTTCCTCCACAGTTTTT<br>CTCCATAATCTTGAAGAGGCCAAAACATTAGCTTTATCCAAGGACCAAATAGGCAATGGTGGCTCATGTTGTAGGGCCAT<br>GAAAGCGGCCATTCTTGTGATTCTTTGCACTTCTGGAACGGTGTATTGTTCACTATCCCAAGCGACACCATCACCATCGT<br>CCTCCTTTCTCTTACCAAAGTAAATACCTCCCACTAATTCTCTGACAACAACGAAGTCAGTACCTTTAGCAAATTGTGGCT<br>TGATTGGAGATAAGTCTAAAAGAGAGTCGGATGCAAAGTTACATGGTCTTAAGTTGGCGTACAATTGAAGTTCTTTACGG<br>ATTTTTAGTAAACCTTGTTCAGGTCTAACACTACCGGTTCCCCATTTAGGACCACCCACAGCACCTAACAAAACGGCATC<br>AGCCTTCTTGGAGGCTTCCAGCGCCTCATCTGGAAGTGGAACACCTGTAGCATCGATAGCAGCACCACCAATTAAATGA<br>TTTTCGAAATCGAACTTGACATTGGAACGAACATCAGAAATAGCTTTAAGAACCTTAATGGCTTCGGCTGTGATTTCTTGA<br>CCAACGTGGTCACCTGGCAAAACGACGATCTTCTTAGGGGCAGACATTTTATAATTATTATAAAAACTTTCATATAAAAAG<br>GAGTTTTTTAGTTACCTTAATTGAAATAAACGAAATAAAAACTCGCCCTCAGACTCAGATTGGTGTTTCAAAGATCCTCAA<br>GGCAATAGAATAACAGATTTTGATAGTATTAATAAAGAATTAGGTCTTGGTAGAAGAGATGTAAAATTAGATAAAGGTCAT<br>GATGATTTAATTAAATTATGTACTGAAAAAATAGATAGTACTATTAATTGCGTTGCGCTCACTGCCCGCTTTCCAGTCGGG<br>AAACCTGTCGTGCCAGCTGCATTAATGAATCGGCCAACGCGCGGGGAGAGGCGGTTTGCGTATTGGGCGCTCTTCCGC<br>TTCCTCGCTCACTGACTCGCTGCGCTCGGTCGTTCGGCTGCGGCGAGCGGTATCAGCTCACTCAAAGGCGGTAATACG<br>GTTATCCACAGAATCAGGGGATAACGCAGGAAAGAACATGTGAGCAAAAGGCCAGCAAAAGGCCAGGAACCGTAAAAA<br>GGCCGCGTTGCTGGCGTTTTTCCATAGGCTCCGCCCCCCTGACGAGCATCACAAAAATCGACGCTCAAGTCAGAGGTG<br>GCGAAACCCGACAGGACTATAAAGATACCAGGCGTTTCCCCCTGGAAGCTCCCTCGTGCGCTCTCCTGTTCCGACCCT<br>GCCGCTTACCGGATACCTGTCCGCCTTTCTCCCTTCGGGAAGCGTGGCGCTTTCTCATAGCTCACGCTGTAGGTATCTC<br>AGTTCGGTGTAGGTCGTTCGCTCCAAGCTGGGCTGTGTGCACGAACCCCCCGTTCAGCCCGACCGCTGCGCCTTATCC<br>GGTAACTATCGTCTTGAGTCCAACCCGGTAAGACACGACTTATCGCCACTGGCAGCAGCCACTGGTAACAGGATTAGCA<br>GAGCGAGGTATGTAGGCGGTGCTACAGAGTTCTTGAAGTGGTGGCCTAACTACGGCTACACTAGAAGAACAGTATTTGG<br>TATCTGCGCTCTGCTGAAGCCAGTTACCTTCGGAAAAAGAGTTGGTAGCTCTTGATCCGGCAAACAAACCACCGCTGGT<br>AGCGGTGGTTTTTTTGTTTGCAAGCAGCAGATTACGCGCAGAAAAAAAGGATCTCAAGAAGATCCTTTGATCTTTTCTAC<br>GGGGTCTGACGCTCAGTGGAACGAAAACTCACGTTAAGGGATTTTGGTCATGAGATTATCAAAAAGGATCTTCACCTAG<br>ATCCTTTTAAATTAAAAATGAAGTTTTAAATCAATCTAAAGTATATATGAGTAAACTTGGTCTGACAGTTACCAATGCTTAA<br>TCAGTGAGGCACCTATCTCAGCGATCTGTCTATTTCGTTCATCCATAGTTGCCTGACTCCCCGTCGTGTAGATAACTACG<br>ATACGGGAGGGCTTACCATCTGGCCCCAGTGCTGCAATGATACCGCGTGACCCACGCTCACCGGCTCCAGATTTATCA<br>GCAATAAACCAGCCAGCCGGAAGGGCCGAGCGCAGAAGTGGTCCTGCAACTTTATCCGCCTCCATCCAGTCTATTAATT<br>GTTGCCGGGAAGCTAGAGTAAGTAGTTCGCCAGTTAATAGTTTGCGCAACGTTGTTGCCATTGCTACAGGCATCGTGGT<br>GTCACGCTCGTCGTTTGGTATGGCTTCATTCAGCTCCGGTTCCCAACGATCAAGGCGAGTTACATGATCCCCCATGTTG<br>TGCAAAAAAGCGGTTAGCTCCTTCGGTCCTCCGATCGTTGTCAGAAGTAAGTTGGCCGCAGTGTTATCACTCATGGTTAT<br>GGCAGCACTGCATAATTCTCTTACTGTCATGCCATCCGTAAGATGCTTTTCTGTGACTGGTGAGTACTCAACCAAGTCAT<br>TCTGAGAATAGTGTATGCGGCGACCGAGTTGCTCTTGCCCGGCGTCAATACGGGATAATACCGCGCCACATAGCAGAA<br>CTTTAAAAGTGCTCATCATTGGAAAACGTTCTTCGGGGCGAAAACTCTCAAGGATCTTACCGCTGTTGAGATCCAGTTCG<br>ATGTAACCCACTCGTGCACCCAACTGATCTTCAGCATCTTTTACTTTCACCAGCGTTTCTGGGTGAGCAAAAACAGGAAG<br>GCAAAATGCCGCAAAAAAGGGAATAAGGGCGACACGGAAATGTTGAATACTCATACTCTTCCTTTTTCAATATTATTGAA<br>GCATTTATCAGGGTTATTGTCTCATGAGCGGATACATATTTGAATGTATTTAGAAAAATAAACAAATAGGGGTTCCGCG |
| pYY82 | Plasmid that replaced sfGFP in pYY62 with RBD10i14. After linearization with ScaI, this plasmid produces donor DNA encoding the RBD10i14-HA-Aga2 fusion protein with a *LEU2* auxotrophic selection marker, flanked by *attP* recombination sites for p1 integration. This plasmid contains AmpR marker for propagation in *E. coli.* | CACATTTCCCCGAAAAGTGCCACCTGACGTCTAAGAAACCATTATTATCATGACATTAACCTATAAAAATAGGCGTATCAC<br>GAGGCCCTTTCGTCTCACGCGTTTCGGTGATGACGGTGAAAACCTCTGACACATGCAGCTCCCGGAGACGAGTACTATA<br>ATATATGAATTACATTATTAATTTAAAAGATACTAGATGGTGCAGCAGCAGCAGCAGCAGTAGTAGGGCCAGTATTGGCG<br>TTTCCCCCTCCTGACTTGAACAACCCTTTTAACGACTTTGAAATAGATAGTGACCTCTTCAAAGAGCTTTCTCTGGAAATA<br>CGGGGCTCAGAAGAGTCCACATTAGATGAAGCCATTATGGATTGCGAGTTTTTATTTCGTTTATTCAAATTAAGGTAACTA<br>AAAAACTCCTTTTGATGACCTATACATAGGAAGATCTATAGAAACAAAAAGATTAATAACTTTCAAATATCAGAAAAATATA<br>GAAACATGTGATAAGCTCATAGACATATAAAAATATGAGATTCCCATCTATCTTCACCGCTGTTTTGTTCGCTGCTTCTTC<br>TGCTTTGGCTGCTCCAGCTAACACCACCACCGAAGACGAAACCGCTCAAATCCCAGCTGAAGCTGTTATCGACTACTCT<br>GACTTGGAAGGTGACTTCGACGCTGCTGCTTTGCCATTGTCTAACTCTACCAACAACGGTTTGTCTTCTACCAACACCAC<br>CATCGCTTCTATCGCTGCTAAGGAAGAAGGTGTTCAATTGGACAAGAGAGAAGCTGACGCACAGGTGCAGCTGCAGGA<br>AAGCGGCGGCGGCCTGGTGCAGGCGGGCGGCAGCCTGCGCCTGAGCTGCGCGGCGAGCGGCACTATTTTTCAGGTT<br>GGTTCTGTGGGCTGGTATCGCCAGGCGCCGGGCAAAGGACGCAAATTTGTTGCCACTATTGCTGACGGTAGTAGTACC<br>AATTATGCGGGTAGCGTGAAAGGCCGCTTTACCATTAGCCGCGATAACGCGAAAAACACCGTGTATCTGCAGATGAACA<br>GCCTGAAACCGGAAGATACCGCGGTGTATTATTGCGCGGCTCTGGGTCAGGTTTCTGAATACAACTCTGCTTCTTACGA<br>ATGGACTTATCCGTATTGGGGCCAGGGCACCCAGGTGACCGTGAGCAGCGGTTCCAAGGACAATAGCTCGACGATTGA<br>AGGTAGATACCCATACGACGTTCCAGACTACGCTCTGCAGGCTAGTGGTGGTGGTGGTTCTGGTGGTGGTGGTTCTGG<br>TGGTGGTGGTTCTGCTAGCGACGTCCAGGAACTGACAACTATATGCGAGCAAATCCCCTCACCAACTTTAGAATCGACG<br>CCGTACTCTTTGTCAACGACTACTATTTTGGCCAACGGGAAGGCAATGCAAGGAGTTTTTGAATATTACAAATCAGTAAC<br>GTTTGTCAGTAATTGCGGTTCTCACCCCTCAACAACTAGCAAAGGCAGCCCCATAAACACACAGTATGTTTTTTGAATCC<br>TAATAATGGCCGCAAAAAAAAAAAAAAAAAAAAAAAAAAAAAAAAAAAAAAAAAAAAAAAAAAAAAAAAAAAAAAAAAA<br>AAAAAACCTGTCACCGGATGTGTTTTCCGGTCTGATGAGTCCGTGAGGACGAAACAGGACTAGGCTGCGAATTTCTTAT<br>GATTTATGATTTTTATTATTAAATAAGTTATAAAAAAAATAAGTGTATACAAATTTTAAAGTGACTCTTAGGTTTTAAAACGA<br>AAATTCTTATTCTTGAGTAACTCTTTCCTGTAGGTCAGGTTGCTTTCTCAGGTATAGCATGAGGTCGCTCTTAAGCAAGGA<br>TTTTCTTAACTTCTTCGGCGACAGCATCACCGACTTCGGTGGTACTGTTGGAACCACCTAAATCACCAGTTCTGATACCT<br>GCATCCAAAACCTTTTTAACTGCATCTTCAATGGCCTTACCTTCTTCAGGCAAGTTCAATGACAATTTCAACATCATTGCA<br>GCAGACAAGATAGTGGCGATAGGGTTGACCTTATTCTTTGGCAAATCTGGAGCAGAACCGTGGCATGGTTCGTACAAAC<br>CAAATGCGGTGTTCTTGTCTGGCAAAGAGGCCAAGGACGCAGATGGCAACAAACCCAAGGAACCTGGGATAACGGAGG<br>CTTCATCGGAGATGATATCACCAAACATGTTGCTGGTGATTATAATACCATTTAGGTGGGTTGGGTTCTTAACTAGGATC<br>ATGGCGGCAGAATCAATCAATTGATGTTGAACCTTCAATGTAGGAAATTCGTTCTTGATGGTTTCCTCCACAGTTTTTCTC<br>CATAATCTTGAAGAGGCCAAAACATTAGCTTTATCCAAGGACCAAATAGGCAATGGTGGCTCATGTTGTAGGGCCATGAA<br>AGCGGCCATTCTTGTGATTCTTTGCACTTCTGGAACGGTGTATTGTTCACTATCCCAAGCGACACCATCACCATCGTCCT<br>CCTTTCTCTTACCAAAGTAAATACCTCCCACTAATTCTCTGACAACAACGAAGTCAGTACCTTTAGCAAATTGTGGCTTGA<br>TTGGAGATAAGTCTAAAAGAGAGTCGGATGCAAAGTTACATGGTCTTAAGTTGGCGTACAATTGAAGTTCTTTACGGATT<br>TTTAGTAAACCTTGTTCAGGTCTAACACTACCGGTTCCCCATTTAGGACCACCCACAGCACCTAACAAAACGGCATCAGC<br>CTTCTTGGAGGCTTCCAGCGCCTCATCTGGAAGTGGAACACCTGTAGCATCGATAGCAGCACCACCAATTAAATGATTTT<br>CGAAATCGAACTTGACATTGGAACGAACATCAGAAATAGCTTTAAGAACCTTAATGGCTTCGGCTGTGATTTCTTGACCA<br>ACGTGGTCACCTGGCAAAACGACGATCTTCTTAGGGGCAGACATTTTATAATTATTATAAAAACTTTCATATAAAAAGGAG<br>TTTTTTAGTTACCTTAATTGAAATAAACGAAATAAAAACTCGCCCTCAGACTCAGATTGGTGTTTCAAAGATCCTCAAGGC<br>AATAGAATAACAGATTTTGATAGTATTAATAAAGAATTAGGTCTTGGTAGAAGAGATGTAAAATTAGATAAAGGTCATGAT<br>GATTTAATTAAATTATGTACTGAAAAAATAGATAGTACTATTAATTGCGTTGCGCTCACTGCCCGCTTTCCAGTCGGGAAA<br>CCTGTCGTGCCAGCTGCATTAATGAATCGGCCAACGCGCGGGGAGAGGCGGTTTGCGTATTGGGCGCTCTTCCGCTTC<br>CTCGCTCACTGACTCGCTGCGCTCGGTCGTTCGGCTGCGGCGAGCGGTATCAGCTCACTCAAAGGCGGTAATACGGTT<br>ATCCACAGAATCAGGGGATAACGCAGGAAAGAACATGTGAGCAAAAGGCCAGCAAAAGGCCAGGAACCGTAAAAAGGC<br>CGCGTTGCTGGCGTTTTTCCATAGGCTCCGCCCCCCTGACGAGCATCACAAAAATCGACGCTCAAGTCAGAGGTGGCG<br>AAACCCGACAGGACTATAAAGATACCAGGCGTTTCCCCCTGGAAGCTCCCTCGTGCGCTCTCCTGTTCCGACCCTGCC<br>GCTTACCGGATACCTGTCCGCCTTTCTCCCTTCGGGAAGCGTGGCGCTTTCTCATAGCTCACGCTGTAGGTATCTCAGT<br>TCGGTGTAGGTCGTTCGCTCCAAGCTGGGCTGTGTGCACGAACCCCCCGTTCAGCCCGACCGCTGCGCCTTATCCGGT<br>AACTATCGTCTTGAGTCCAACCCGGTAAGACACGACTTATCGCCACTGGCAGCAGCCACTGGTAACAGGATTAGCAGAG<br>CGAGGTATGTAGGCGGTGCTACAGAGTTCTTGAAGTGGTGGCCTAACTACGGCTACACTAGAAGAACAGTATTTGGTAT<br>CTGCGCTCTGCTGAAGCCAGTTACCTTCGGAAAAAGAGTTGGTAGCTCTTGATCCGGCAAACAAACCACCGCTGGTAGC<br>GGTGGTTTTTTTGTTTGCAAGCAGCAGATTACGCGCAGAAAAAAAGGATCTCAAGAAGATCCTTTGATCTTTTCTACGGG<br>GTCTGACGCTCAGTGGAACGAAAACTCACGTTAAGGGATTTTGGTCATGAGATTATCAAAAAGGATCTTCACCTAGATCC |

| | | |
|---|---|---|
| | | TTTTAAATTAAAAATGAAGTTTTAAATCAATCTAAAGTATATATGAGTAAACTTGGTCTGACAGTTACCAATGCTTAATCAG<br>TGAGGCACCTATCTCAGCGATCTGTCTATTTCGTTCATCCATAGTTGCCTGACTCCCCGTCGTGTAGATAACTACGATAC<br>GGGAGGGCTTACCATCTGGCCCCAGTGCTGCAATGATACCGCGTGACCCACGCTCACCGGCTCCAGATTTATCAGCAA<br>TAAACCAGCCAGCCGGAAGGGCCGAGCGCAGAAGTGGTCCTGCAACTTTATCCGCCTCCATCCAGTCTATTAATTGTTG<br>CCGGGAAGCTAGAGTAAGTAGTTCGCCAGTTAATAGTTTGCGCAACGTTGTTGCCATTGCTACAGGCATCGTGGTGTCA<br>CGCTCGTCGTTTGGTATGGCTTCATTCAGCTCCGGTTCCCAACGATCAAGGCGAGTTACATGATCCCCCATGTTGTGCA<br>AAAAAGCGGTTAGCTCCTTCGGTCCTCCGATCGTTGTCAGAAGTAAGTTGGCCGCAGTGTTATCACTCATGGTTATGGC<br>AGCACTGCATAATTCTCTTACTGTCATGCCATCCGTAAGATGCTTTTCTGTGACTGGTGAGTACTCAACCAAGTCATTCT<br>GAGAATAGTGTATGCGGCGACCGAGTTGCTCTTGCCCGGCGTCAATACGGGATAATACCGCGCCACATAGCAGAACTT<br>TAAAAGTGCTCATCATTGGAAAACGTTCTTCGGGGCGAAAACTCTCAAGGATCTTACCGCTGTTGAGATCCAGTTCGATG<br>TAACCCACTCGTGCACCCAACTGATCTTCAGCATCTTTTACTTTCACCAGCGTTTCTGGGTGAGCAAAAACAGGAAGGCA<br>AAATGCCGCAAAAAAGGGAATAAGGGCGACACGGAAATGTTGAATACTCATACTCTTCCTTTTTCAATATTATTGAAGCAT<br>TTATCAGGGTTATTGTCTCATGAGCGGATACATATTTGAATGTATTTAGAAAAATAAACAAATAGGGGTTCCGCG |
| pYY83 | Plasmid that replaced sfGFP in pYY62 with Nb.b201. After linearization with ScaI, this plasmid produces donor DNA encoding the Nb.b201-HA-Aga2 fusion protein with a *LEU2* auxotrophic selection marker, flanked by *attP* recombination sites for p1 integration. This plasmid contains AmpR marker for propagation in *E. coli.* | CACATTTCCCCGAAAAGTGCCACCTGACGTCTAAGAAACCATTATTATCATGACATTAACCTATAAAAATAGGCGTATCAC<br>GAGGCCCTTTCGTCTCACGCGTTTCGGTGATGACGGTGAAAACCTCTGACACATGCAGCTCCCGGAGACGAGTACTATA<br>ATATATGAATTACATTATTAATTTAAAAGATACTAGATGGTGCAGCAGCAGCAGCAGCAGTAGTAGGGCCAGTATTGGCG<br>TTTCCCCCTCCTGACTTGAACAACCCTTTTAACGACTTTGAAATAGATAGTGACCTCTTCAAAGAGCTTTCTCTGGAAATA<br>CGGGGCTCAGAAGAGTCCACATTAGATGAAGCCATTATGGATTGCGAGTTTTTATTTCGTTTATTCAAATTAAGGTAACTA<br>AAAAACTCCTTTTGATGACCTATACATAGGAAGATCTATAGAAACAAAAAGATTAATAACTTTCAAATATCAGAAAAATATA<br>GAAACATGTGATAAGCTCATAGACATATAAAAATATGAGATTCCCATCTATCTTCACCGCTGTTTTGTTCGCTGCTTCTTC<br>TGCTTTGGCTGCTCCAGCTAACACCACCACCGAAGACGAAACCGCTCAAATCCCAGCTGAAGCTGTTATCGACTACTCT<br>GACTTGGAAGGTGACTTCGACGCTGCTGCTTTGCCATTGTCTAACTCTACCAACAACGGTTTGTCTTCTACCAACACCAC<br>CATCGCTTCTATCGCTGCTAAGGAAGAAGGTGTTCAATTGGACAAGAGAGAAGCTGACGCACAGGTGCAGCTGCAGGA<br>AAGCGGCGGCGGCCTGGTGCAGGCGGGCGGCAGCCTGCGCCTGAGCTGCGCGGCGAGCGGCTATATTTCTGACGCT<br>TACTACATGGGCTGGTATCGCCAGGCGCCGGGCAAAGAACGCGAATTTGTTGCCACTATTACTCATGGTACTAATACCT<br>ATTATGCGGATAGCGTGAAAGGCCGCTTTACCATTAGCCGCGATAACGCGAAAAACACCGTGTATCTGCAGATGAACAG<br>CCTGAAACCGGAAGATACCGCGGTGTATTATTGCGCGGTTCTGGAAACTCGTTCTTACTCTTTTCGTTATTGGGGCCAG<br>GGCACCCAGGTGACCGTGAGCAGCGGTTCCAAGGACAATAGCTCGACGATTGAAGGTAGATACCCATACGACGTTCCA<br>GACTACGCTCTGCAGGCTAGTGGTGGTGGTGGTTCTGGTGGTGGTGGTTCTGGTGGTGGTGGTTCTGCTAGCGACGTC<br>CAGGAACTGACAACTATATGCGAGCAAATCCCCTCACCAACTTTAGAATCGACGCCGTACTCTTTGTCAACGACTACTAT<br>TTTGGCCAACGGGAAGGCAATGCAAGGAGTTTTTGAATATTACAAATCAGTAACGTTTGTCAGTAATTGCGGTTCTCACC<br>CCTCAACAACTAGCAAAGGCAGCCCCATAAACACACAGTATGTTTTTTGAATCCTAATAATGGCCGCAAAAAAAAAAAAA<br>AAAAAAAAAAAAAAAAAAAAAAAAAAAAAAAAAAAAAAAAAAAAAAAAAAAAAAAAAACCTGTCACCGGATGTGTTTT<br>CCGGTCTGATGAGTCCGTGAGGACGAAACAGGACTAGGCTGCGAATTTCTTATGATTTATGATTTTTATTATTAAATAAGT<br>TATAAAAAAAATAAGTGTATACAAATTTTAAAGTGACTCTTAGGTTTTAAAACGAAAATTCTTATTCTTGAGTAACTCTTTCC<br>TGTAGGTCAGGTTGCTTTCTCAGGTATAGCATGAGGTCGCTCTTAAGCAAGGATTTTCTTAACTTCTTCGGCGACAGCAT<br>CACCGACTTCGGTGGTACTGTTGGAACCACCTAAATCACCAGTTCTGATACCTGCATCCAAAACCTTTTTAACTGCATCT<br>TCAATGGCCTTACCTTCTTCAGGCAAGTTCAATGACAATTTCAACATCATTGCAGCAGACAAGATAGTGGCGATAGGGTT<br>GACCTTATTCTTTGGCAAATCTGGAGCAGAACCGTGGCATGGTTCGTACAAACCAAATGCGGTGTTCTTGTCTGGCAAA<br>GAGGCCAAGGACGCAGATGGCAACAAACCCAAGGAACCTGGGATAACGGAGGCTTCATCGGAGATGATATCACCAAAC<br>ATGTTGCTGGTGATTATAATACCATTTAGGTGGGTTGGGTTCTTAACTAGGATCATGGCGGCAGAATCAATCAATTGATG<br>TTGAACCTTCAATGTAGGAAATTCGTTCTTGATGGTTTCCTCCACAGTTTTTCTCCATAATCTTGAAGAGGCCAAAACATT<br>AGCTTTATCCAAGGACCAAATAGGCAATGGTGGCTCATGTTGTAGGGCCATGAAAGCGGCCATTCTTGTGATTCTTTGC<br>ACTTCTGGAACGGTGTATTGTTCACTATCCCAAGCGACACCATCACCATCGTCCTCCTTTCTCTTACCAAAGTAAATACCT<br>CCCACTAATTCTCTGACAACAACGAAGTCAGTACCTTTAGCAAATTGTGGCTTGATTGGAGATAAGTCTAAAAGAGAGTC<br>GGATGCAAAGTTACATGGTCTTAAGTTGGCGTACAATTGAAGTTCTTTACGGATTTTTAGTAAACCTTGTTCAGGTCTAAC<br>ACTACCGGTTCCCCATTTAGGACCACCCACAGCACCTAACAAAACGGCATCAGCCTTCTTGGAGGCTTCCAGCGCCTCA<br>TCTGGAAGTGGAACACCTGTAGCATCGATAGCAGCACCACCAATTAAATGATTTTCGAAATCGAACTTGACATTGGAACG<br>AACATCAGAAATAGCTTTAAGAACCTTAATGGCTTCGGCTGTGATTTCTTGACCAACGTGGTCACCTGGCAAAACGACGA<br>TCTTCTTAGGGGCAGACATTTTATAATTATTATAAAAACTTTCATATAAAAAGGAGTTTTTTAGTTACCTTAATTGAAATAAA<br>CGAAATAAAAACTCGCCCTCAGACTCAGATTGGTGTTTCAAAGATCCTCAAGGCAATAGAATAACAGATTTTGATAGTATT<br>AATAAAGAATTAGGTCTTGGTAGAAGAGATGTAAAATTAGATAAAGGTCATGATGATTTAATTAAATTATGTACTGAAAAAA<br>TAGATAGTACTATTAATTGCGTTGCGCTCACTGCCCGCTTTCCAGTCGGGAAACCTGTCGTGCCAGCTGCATTAATGAAT<br>CGGCCAACGCGCGGGGAGAGGCGGTTTGCGTATTGGGCGCTCTTCCGCTTCCTCGCTCACTGACTCGCTGCGCTCGG<br>TCGTTCGGCTGCGGCGAGCGGTATCAGCTCACTCAAAGGCGGTAATACGGTTATCCACAGAATCAGGGGATAACGCAG<br>GAAAGAACATGTGAGCAAAAGGCCAGCAAAAGGCCAGGAACCGTAAAAAGGCCGCGTTGCTGGCGTTTTTCCATAGGC<br>TCCGCCCCCCTGACGAGCATCACAAAAATCGACGCTCAAGTCAGAGGTGGCGAAACCCGACAGGACTATAAAGATACC<br>AGGCGTTTCCCCCTGGAAGCTCCCTCGTGCGCTCTCCTGTTCCGACCCTGCCGCTTACCGGATACCTGTCCGCCTTTCT<br>CCCTTCGGGAAGCGTGGCGCTTTCTCATAGCTCACGCTGTAGGTATCTCAGTTCGGTGTAGGTCGTTCGCTCCAAGCTG<br>GGCTGTGTGCACGAACCCCCCGTTCAGCCCGACCGCTGCGCCTTATCCGGTAACTATCGTCTTGAGTCCAACCCGGTA<br>AGACACGACTTATCGCCACTGGCAGCAGCCACTGGTAACAGGATTAGCAGAGCGAGGTATGTAGGCGGTGCTACAGAG<br>TTCTTGAAGTGGTGGCCTAACTACGGCTACACTAGAAGAACAGTATTTGGTATCTGCGCTCTGCTGAAGCCAGTTACCTT<br>CGGAAAAAGAGTTGGTAGCTCTTGATCCGGCAAACAAACCACCGCTGGTAGCGGTGGTTTTTTTGTTTGCAAGCAGCAG<br>ATTACGCGCAGAAAAAAAGGATCTCAAGAAGATCCTTTGATCTTTTCTACGGGGTCTGACGCTCAGTGGAACGAAAACTC<br>ACGTTAAGGGATTTTGGTCATGAGATTATCAAAAAGGATCTTCACCTAGATCCTTTTAAATTAAAAATGAAGTTTTAAATCA<br>ATCTAAAGTATATATGAGTAAACTTGGTCTGACAGTTACCAATGCTTAATCAGTGAGGCACCTATCTCAGCGATCTGTCTA<br>TTTCGTTCATCCATAGTTGCCTGACTCCCCGTCGTGTAGATAACTACGATACGGGAGGGCTTACCATCTGGCCCCAGTG<br>CTGCAATGATACCGCGTGACCCACGCTCACCGGCTCCAGATTTATCAGCAATAAACCAGCCAGCCGGAAGGGCCGAGC<br>GCAGAAGTGGTCCTGCAACTTTATCCGCCTCCATCCAGTCTATTAATTGTTGCCGGGAAGCTAGAGTAAGTAGTTCGCC<br>AGTTAATAGTTTGCGCAACGTTGTTGCCATTGCTACAGGCATCGTGGTGTCACGCTCGTCGTTTGGTATGGCTTCATTCA<br>GCTCCGGTTCCCAACGATCAAGGCGAGTTACATGATCCCCCATGTTGTGCAAAAAAGCGGTTAGCTCCTTCGGTCCTCC<br>GATCGTTGTCAGAAGTAAGTTGGCCGCAGTGTTATCACTCATGGTTATGGCAGCACTGCATAATTCTCTTACTGTCATGC<br>CATCCGTAAGATGCTTTTCTGTGACTGGTGAGTACTCAACCAAGTCATTCTGAGAATAGTGTATGCGGCGACCGAGTTG<br>CTCTTGCCCGGCGTCAATACGGGATAATACCGCGCCACATAGCAGAACTTTAAAAGTGCTCATCATTGGAAAACGTTCTT<br>CGGGGCGAAAACTCTCAAGGATCTTACCGCTGTTGAGATCCAGTTCGATGTAACCCACTCGTGCACCCAACTGATCTTC<br>AGCATCTTTTACTTTCACCAGCGTTTCTGGGTGAGCAAAAACAGGAAGGCAAAATGCCGCAAAAAAGGGAATAAGGGCG<br>ACACGGAAATGTTGAATACTCATACTCTTCCTTTTTCAATATTATTGAAGCATTTATCAGGGTTATTGTCTCATGAGCGGA<br>TACATATTTGAATGTATTTAGAAAAATAAACAAATAGGGGTTCCGCG |
| pRW397 | Plamids for generation of donor DNA for construction of a PhiC31-compatible landing pad p1. After linearization with ScaI, this plasmid produces donor DNA with the | ACTATCTATTTTTTCAGTACATAATTTAATTAAATCATCATGACCTTTATCTAATTTTACATCTCTTCTACCAAGACCTAATT<br>CTTTATTAATACTATCAAAATCTGTTATTCTATTGCCTTGAGGATCTTTGAAACACCAATCTGAGTCTGAGGGCATGCGTG<br>CCAGGGCGTGCCCCGGGGCTCCCCGGGCGCGTTTTTCGCCGGAGTCAATTAGGTCATACTTTTCTATATAATCCAAATC<br>CCAAAAATCAATTGAATGATTCTTAATATGATTTAATAGTTTATGATTATAAATGTCGAAAGCTACATATAAGGAACGTGCT<br>GCTACTCATCCTAGTCCTGTTGCTGCCAAGCTATTTAATATCATGCACGAAAAGCAAACAAACTTGTGTGCTTCATTGGAT<br>GTTCGTACCACCAAGGAATTACTGGAGTTAGTTGAAGCATTAGGTCCCAAAATTTGTTTACTAAAAACACATGTGGATATC<br>TTGACTGATTTTTCCATGGAGGGCACAGTTAAGCCGCTAAAGGCATTATCCGCCAAGTACAATTTTTTACTCTTCGAGGA<br>CAGAAAATTTGCTGACATTGGTAATACAGTCAAATTGCAGTACTCTGCGGGTGTATACAGAATAGCAGAATGGGCAGACA<br>TTACGAATGCACACGGTGTGGTGGGCCCAGGTATTGTTAGCGGTTTGAAGCAGGCGGCAGAAGAAGTAACAAAGGAAC<br>CTAGAGGCCTTTTGATGTTAGCAGAATTGTCATGCAAGGGCTCCCTATCTACTGGAGAATATACTAAGGGTACTGTTGAC<br>ATTGCGAAGAGCGACAAAGATTTTGTTATCGGCTTTATTGCTCAAAGAGACATGGGTGGAAGAGATGAAGGTTACGATT<br>GGTTGATTATGACACCCGGTGTGGGTTTAGATGACAAGGGAGATGCATTGGGTCAACAGTATAGAACCGTGGATGATGT<br>GGTTTCTACAGGATCTGACATTATTATTGTTGGAAGAGGACTATTTGCAAAGGGAAGGGATGCTAAGGTAGAGGGTGAA<br>CGTTACAGAAAAGCAGGCTGGGAAGCATATTTGAGAAGATGCGGCCAGCAAAACTAACGCGCCCGGGGAGCCCGTGG<br>GCACGCCCTGGCACAATCCATAATGGCTTCATCTAATGTGGACTCTTCTGAGCCCCGTATTTCCAGAGAAAGCTCTTTGA<br>AGAGGTCTCTATCTATTTCAAAGTCGTTAAAAGGGTTGTTCAAGTCAGGAGGGGGAAACGCCAATACTGGCCCTACTAC<br>TGCTGCTGCTGCTGCTGCACCATCTAGTATCTTTTAAATTAATAATGTAATTCATATATTATAGTACTCGTCTCCGGGAGC<br>TGCATGTGTCAGAGGTTTTCACCGTCATCACCGAAACGCGTGAGACGAAAGGGCCTCGTGATACGCCTATTTTTATAGG<br>TTAATGTCATGATAATAATGGTTTCTTAGACGTCAGGTGGCACTTTTCGGGGAAATGTGCGCGGAACCCCTATTTGTTTA |

| | | |
|---|---|---|
| | *URA3* selectable marker, *attB* recombination sites for PhiC31, and homology flanks matches the original landing pad p1 used in the lab[1]. This plasmid contains AmpR marker for propagation in *E. coli.* | TTTTTCTAAATACATTCAAATATGTATCCGCTCATGAGACAATAACCCTGATAAATGCTTCAATAATATTGAAAAAGGAAGA<br>GTATGAGTATTCAACATTTCCGTGTCGCCCTTATTCCCTTTTTTGCGGCATTTTGCCTTCCTGTTTTTGCTCACCCAGAAA<br>CGCTGGTGAAAGTAAAAGATGCTGAAGATCAGTTGGGTGCACGAGTGGGTTACATCGAACTGGATCTCAACAGCGGTAA<br>GATCCTTGAGAGTTTTCGCCCCGAAGAACGTTTTCCAATGATGAGCACTTTTAAAGTTCTGCTATGTGGCGCGGTATTAT<br>CCCGTATTGACGCCGGGCAAGAGCAACTCGGTCGCCGCATACACTATTCTCAGAATGACTTGGTTGAGTACTCACCAGT<br>CACAGAAAAGCATCTTACGGATGGCATGACAGTAAGAGAATTATGCAGTGCTGCCATAACCATGAGTGATAACACTGCG<br>GCCAACTTACTTCTGACAACGATCGGAGGACCGAAGGAGCTAACCGCTTTTTTGCACAACATGGGGGATCATGTAACTC<br>GCCTTGATCGTTGGGAACCGGAGCTGAATGAAGCCATACCAAACGACGAGCGTGACACCACGATGCCTGTAGCAATGG<br>CAACAACGTTGCGCAAACTATTAACTGGCGAACTACTTACTCTAGCTTCCCGGCAACAATTAATAGACTGGATGGAGGC<br>GGATAAAGTTGCAGGACCACTTCTGCGCTCGGCCCTTCCGGCTGGCTGGTTTATTGCTGATAAATCTGGAGCCGGTGA<br>GCGTGGGTCTCGCGGTATCATTGCAGCACTGGGGCCAGATGGTAAGCCCTCCCGTATCGTAGTTATCTACACGACGGG<br>GAGTCAGGCAACTATGGATGAACGAAATAGACAGATCGCTGAGATAGGTGCCTCACTGATTAAGCATTGGTAACTGTCA<br>GACCAAGTTTACTCATATATACTTTAGATTGATTTAAAACTTCATTTTTAATTTAAAAGGATCTAGGTGAAGATCCTTTTTGA<br>TAATCTCATGACCAAAATCCCTTAACGTGAGTTTTCGTTCCACTGAGCGTCAGACCCCGTAGAAAAGATCAAAGGATCTT<br>CTTGAGATCCTTTTTTTCTGCGCGTAATCTGCTGCTTGCAAACAAAAAAACCACCGCTACCAGCGGTGGTTTGTTTGCCG<br>GATCAAGAGCTACCAACTCTTTTTCCGAAGGTAACTGGCTTCAGCAGAGCGCAGATACCAAATACTGTTCTTCTAGTGTA<br>GCCGTAGTTAGGCCACCACTTCAAGAACTCTGTAGCACCGCCTACATACCTCGCTCTGCTAATCCTGTTACCAGTGGCT<br>GCTGCCAGTGGCGATAAGTCGTGTCTTACCGGGTTGGACTCAAGACGATAGTTACCGGATAAGGCGCAGCGGTCGGGC<br>TGAACGGGGGGTTCGTGCACACAGCCCAGCTTGGAGCGAACGACCTACACCGAACTGAGATACCTACAGCGTGAGCTA<br>TGAGAAAGCGCCACGCTTCCCGAAGGGAGAAAGGCGGACAGGTATCCGGTAAGCGGCAGGGTCGGAACAGGAGAGC<br>GCACGAGGGAGCTTCCAGGGGGAAACGCCTGGTATCTTTATAGTCCTGTCGGGTTTCGCCACCTCTGACTTGAGCGTC<br>GATTTTTGTGATGCTCGTCAGGGGGGCGGAGCCTATGGAAAAACGCCAGCAACGCGGCCTTTTTACGGTTCCTGGCCT<br>TTTGCTGGCCTTTTGCTCACATGTTCTTTCCTGCGTTATCCCCTGATTCTGTGGATAACCGTATTACCGCCTTTGAGTGA<br>GCTGATACCGCTCGCCGCAGCCGAACGACCGAGCGCAGCGAGTCAGTGAGCGAGGAAGCGGAAGAGCGCCCAATAC<br>GCAAACCGCCTCTCCCCGCGCGTTGGCCGATTCATTAATGCAGCTGGCACGACAGGTTTCCCGACTGGAAAGCGGGCA<br>GTGAGCGCAACGCAATTAATAGT |
| pRW398 | Plamids for generation of donor DNA for construction of a BXB1-compatible landing pad p1. After linearization with ScaI, this plasmid produces donor DNA with the *URA3* selectable marker, *attB* recombination sites for BXB1, and homology flanks matches the original landing pad p1 used in the lab[1]. This plasmid contains AmpR marker for propagation in *E. coli.* | ACTATCTATTTTTTCAGTACATAATTTAATTAAATCATCATGACCTTTATCTAATTTTACATCTCTTCTACCAAGACCTAATT<br>CTTTATTAATACTATCAAAATCTGTTATTCTATTGCCTTGAGGATCTTTGAAACACCAATCTGAGTCTGAGGGCATGCGGC<br>TTGTCGACGACGGCGTGCTCCGTCGTCAGGATCATTTTTTCGCCGGAGTCAATTAGGTCATACTTTTCTATATAATCCAA<br>ATCCCAAAAATCAATTGAATGATTCTTAATATGATTTAATAGTTTATGATTATAAATGTCGAAAGCTACATATAAGGAACGT<br>GCTGCTACTCATCCTAGTCCTGTTGCTGCCAAGCTATTTAATATCATGCACGAAAAGCAAACAAACTTGTGTGCTTCATT<br>GGATGTTCGTACCACCAAGGAATTACTGGAGTTAGTTGAAGCATTAGGTCCCAAAATTTGTTTACTAAAAACACATGTGG<br>ATATCTTGACTGATTTTTCCATGGAGGGCACAGTTAAGCCGCTAAAGGCATTATCCGCCAAGTACAATTTTTTACTCTTCG<br>AGGACAGAAAATTTGCTGACATTGGTAATACAGTCAAATTGCAGTACTCTGCGGGTGTATACAGAATAGCAGAATGGGC<br>AGACATTACGAATGCACACGGTGTGGTGGGCCCAGGTATTGTTAGCGGTTTGAAGCAGGCGGCAGAAGAAGTAACAAA<br>GGAACCTAGAGGCCTTTTGATGTTAGCAGAATTGTCATGCAAGGGCTCCCTATCTACTGGAGAATATACTAAGGGTACT<br>GTTGACATTGCGAAGAGCGACAAAGATTTTGTTATCGGCTTTATTGCTCAAAGAGACATGGGTGGAAGAGATGAAGGTT<br>ACGATTGGTTGATTATGACACCCGGTGTGGGTTTAGATGACAAGGGAGATGCATTGGGTCAACAGTATAGAACCGTGGA<br>TGATGTGGTTTCTACAGGATCTGACATTATTATTGTTGGAAGAGGACTATTTGCAAAGGGAAGGGATGCTAAGGTAGAG<br>GGTGAACGTTACAGAAAAGCAGGCTGGGAAGCATATTTGAGAAGATGCGGCCAGCAAAACTAAATGATCCTGACGACG<br>GAGAGCGCCGTCGTCGACAAGCCAATCCATAATGGCTTCATCTAATGTGGACTCTTCTGAGCCCCGTATTTCCAGAGAA<br>AGCTCTTTGAAGAGGTCTCTATCTATTTCAAAGTCGTTAAAAGGGTTGTTCAAGTCAGGAGGGGGAAACGCCAATACTG<br>GCCCTACTACTGCTGCTGCTGCTGCTGCACCATCTAGTATCTTTTAAATTAATAATGTAATTCATATATTATAGTACTCGTC<br>TCCGGGAGCTGCATGTGTCAGAGGTTTTCACCGTCATCACCGAAACGCGTGAGACGAAAGGGCCTCGTGATACGCCTA<br>TTTTTATAGGTTAATGTCATGATAATAATGGTTTCTTAGACGTCAGGTGGCACTTTTCGGGGAAATGTGCGCGGAACCCC<br>TATTTGTTTATTTTTCTAAATACATTCAAATATGTATCCGCTCATGAGACAATAACCCTGATAAATGCTTCAATAATATTGAA<br>AAAGGAAGAGTATGAGTATTCAACATTTCCGTGTCGCCCTTATTCCCTTTTTTGCGGCATTTTGCCTTCCTGTTTTTGCTC<br>ACCCAGAAACGCTGGTGAAAGTAAAAGATGCTGAAGATCAGTTGGGTGCACGAGTGGGTTACATCGAACTGGATCTCAA<br>CAGCGGTAAGATCCTTGAGAGTTTTCGCCCCGAAGAACGTTTTCCAATGATGAGCACTTTTAAAGTTCTGCTATGTGGCG<br>CGGTATTATCCCGTATTGACGCCGGGCAAGAGCAACTCGGTCGCCGCATACACTATTCTCAGAATGACTTGGTTGAGTA<br>CTCACCAGTCACAGAAAAGCATCTTACGGATGGCATGACAGTAAGAGAATTATGCAGTGCTGCCATAACCATGAGTGAT<br>AACACTGCGGCCAACTTACTTCTGACAACGATCGGAGGACCGAAGGAGCTAACCGCTTTTTTGCACAACATGGGGGATC<br>ATGTAACTCGCCTTGATCGTTGGGAACCGGAGCTGAATGAAGCCATACCAAACGACGAGCGTGACACCACGATGCCTG<br>TAGCAATGGCAACAACGTTGCGCAAACTATTAACTGGCGAACTACTTACTCTAGCTTCCCGGCAACAATTAATAGACTGG<br>ATGGAGGCGGATAAAGTTGCAGGACCACTTCTGCGCTCGGCCCTTCCGGCTGGCTGGTTTATTGCTGATAAATCTGGA<br>GCCGGTGAGCGTGGGTCTCGCGGTATCATTGCAGCACTGGGGCCAGATGGTAAGCCCTCCCGTATCGTAGTTATCTAC<br>ACGACGGGGAGTCAGGCAACTATGGATGAACGAAATAGACAGATCGCTGAGATAGGTGCCTCACTGATTAAGCATTGGT<br>AACTGTCAGACCAAGTTTACTCATATATACTTTAGATTGATTTAAAACTTCATTTTTAATTTAAAAGGATCTAGGTGAAGAT<br>CCTTTTTGATAATCTCATGACCAAAATCCCTTAACGTGAGTTTTCGTTCCACTGAGCGTCAGACCCCGTAGAAAAGATCA<br>AAGGATCTTCTTGAGATCCTTTTTTTCTGCGCGTAATCTGCTGCTTGCAAACAAAAAAACCACCGCTACCAGCGGTGGTT<br>TGTTTGCCGGATCAAGAGCTACCAACTCTTTTTCCGAAGGTAACTGGCTTCAGCAGAGCGCAGATACCAAATACTGTTCT<br>TCTAGTGTAGCCGTAGTTAGGCCACCACTTCAAGAACTCTGTAGCACCGCCTACATACCTCGCTCTGCTAATCCTGTTAC<br>CAGTGGCTGCTGCCAGTGGCGATAAGTCGTGTCTTACCGGGTTGGACTCAAGACGATAGTTACCGGATAAGGCGCAGC<br>GGTCGGGCTGAACGGGGGGTTCGTGCACACAGCCCAGCTTGGAGCGAACGACCTACACCGAACTGAGATACCTACAG<br>CGTGAGCTATGAGAAAGCGCCACGCTTCCCGAAGGGAGAAAGGCGGACAGGTATCCGGTAAGCGGCAGGGTCGGAAC<br>AGGAGAGCGCACGAGGGAGCTTCCAGGGGGAAACGCCTGGTATCTTTATAGTCCTGTCGGGTTTCGCCACCTCTGACT<br>TGAGCGTCGATTTTTGTGATGCTCGTCAGGGGGGCGGAGCCTATGGAAAAACGCCAGCAACGCGGCCTTTTTACGGTT<br>CCTGGCCTTTTGCTGGCCTTTTGCTCACATGTTCTTTCCTGCGTTATCCCCTGATTCTGTGGATAACCGTATTACCGCCT<br>TTGAGTGAGCTGATACCGCTCGCCGCAGCCGAACGACCGAGCGCAGCGAGTCAGTGAGCGAGGAAGCGGAAGAGCG<br>CCCAATACGCAAACCGCCTCTCCCCGCGCGTTGGCCGATTCATTAATGCAGCTGGCACGACAGGTTTCCCGACTGGAA<br>AGCGGGCAGTGAGCGCAACGCAATTAATAGT |
| pRW402 | Plasmid for genertaion of donor DNA for BXB1-compatible p1 integration in yeast. After linearization with ScaI, this plasmid produces donor DNA encoding the fluorescent protein mScarletI as a reporter with a *LEU2* auxotrophic | ACTATCTATTTTTTCAGTACATAATTTAATTAAATCATCATGACCTTTATCTAATTTTACATCTCTTCTACCAAGACCTAATT<br>CTTTATTAATACTATCAAAATCTGTTATTCTATTGCCTTGAGGATCTTTGAAACACCAATCTGAGTCTGAGGGGTTTGTCT<br>GGTCAACCACCGCGTGCTCAGTGGTGTACGGTACAAACCTCTGATGAGACGGAGAGGTCAGCTTTTCTTTGCTAATATC<br>CAGTGAACTACCAATCATTATAGGAAGTTCTCTGTCATTGTATATAGCACAGTAACTTAGGATTTAAGAACCTCTAGAACC<br>AGTAGAACCAGGACCAGCAGTACCATCAACAGCAGAATTAGAAGCTTGAGCCCTAGATCTCAAACCAGATTTATACAATT<br>CATCCATACCACCAGTAGAATGTCTACCTTCAGATCTTTCATATTGTTCAACAACAGTATAATCCTCGTTATGTGAAGTAA<br>TGTCAAGTTTTCTATCTACATTATAAGCACCCGGCATTTGAACAGGTTTTTTAGCTTTATAAGTAGTTTTAAAATCAGCCAA<br>ATATCTACCACCATCTTTCAATCTCAAAGCCATTTTAATATCACCTTTCAAAACACCATCTTCAGGATACAATCTTTCAGTA<br>GAAGCTTCCCAACCCATAGTTTTTTTTTGCATAACAGGACCATCAGGTGGAAAATTAGTACCTCTCAATTTAACTTTATAA<br>ATCAAAGTACCATCTTCCAAAGAAGTATCTTGAGTAACAGTAACAGCACCACCATCTTCAAAATTCATAACTCTTTCCCAT<br>TTAAAACCTTCAGGAAAAGATTGTTTATAATAATCAGGAATATCAGCAGGATGTTTAATAAAAGCTCTAGAACCATACATA<br>AATTGTGGAGACAAAATATCCCAAGAAAATGGCAATGGACCACCTTTAGTAACTTTCAATTTAGCAGTTTGAGTACCTTCA<br>TAAGGTCTACCTTCACCTTCACCTTCAATTTCAAATTCATGACCATTCATAGAACCTTCCATGTGAACTTTAAATCTCATAA<br>ATTCTTTAATAACAGCTTCACCTTTAGAAACCATATTTTTATATGTCTATGAGCTTATCACATGTTTCTATATTTTTCTGATA<br>TTTGAAAGTTATTAATCTTTTTGTTTCTATAGATCTTCCTATGTATAGGTCATCCGTTATCGTCTCAAGCAGTACAGAAAAT<br>TAATTTCTAATGTGAACACTAGTGAGATAGCTAAAGATCTTAATTGTGAAAACAATATTGAAAGTATAATAAATACATTAAA<br>AGAACAAAATAGATATTTTGACAAACAAATTGAATATATGTCTGCCCCTAAGAAGATCGTCGTTTTGCCAGGTGACCACG<br>TTGGTCAAGAAATCACAGCCGAAGCCATTAAGGTTCTTAAAGCTATTTCTGATGTTCGTTCCAATGTCAAGTTCGATTTCG<br>AAAATCATTTAATTGGTGGTGCTGCTATCGATGCTACAGGTGTTCCACTTCCAGATGAGGCGCTGGAAGCCTCCAAGAA<br>GGCTGATGCCGTTTTGTTAGGTGCTGTGGGTGGTCCTAAATGGGGAACCGGTAGTGTTAGACCTGAACAAGGTTTACTA<br>AAAATCCGTAAAGAACTTCAATTGTACGCCAACTTAAGACCATGTAACTTTGCATCCGACTCTCTTTTAGACTTATCTCCA<br>ATCAAGCCACAATTTGCTAAAGGTACTGACTTCGTTGTTGTCAGAGAATTAGTGGGAGGTATTTACTTTGGTAAGAGAAA<br>GGAGGACGATGGTGATGGTGTCGCTTGGGATAGTGAACAATACACCGTTCCAGAAGTGCAAAGAATCACAAGAATGGC<br>CGCTTTCATGGCCCTACAACATGAGCCACCATTGCCTATTTGGTCCTTGGATAAAGCTAATGTTTTGGCCTCTTCAAGAT<br>TATGGAGAAAAACTGTGGAGGAAACCATCAAGAACGAATTTCCTACATTGAAGGTTCAACATCAATTGATTGATTCTGCC<br>GCCATGATCCTAGTTAAGAACCCAACCCACCTAAATGGTATTATAATCACCAGCAACATGTTTGGTGATATCATCTCCGA |

| | | |
|---|---|---|
| | selection marker, flanked by *attP* recombination sites for BXB1. This plasmid contains AmpR marker for propagation in *E. coli.* | TGAAGCCTCCGTTATCCCAGGTTCCTTGGGTTTGTTGCCATCTGCGTCCTTGGCCTCTTTGCCAGACAAGAACACCGCA<br>TTTGGTTTGTACGAACCATGCCACGGTTCTGCTCCAGATTTGCCAAAGAATAAGGTCAACCCTATCGCCACTATCTTGTC<br>TGCTGCAATGATGTTGAAATTGTCATTGAACTTGCCTGAAGAAGGTAAGGCCATTGAAGATGCAGTTAAAAAGGTTTTGG<br>ATGCAGGTATCAGAACTGGTGATTTAGGTGGTTCCAACAGTACCACCGAAGTCGGTGATGCTGTCGCCGAAGAAGTTAA<br>GAAAATCCTTGCTTAAAAGAGAGTGAGGGTTTGTACCGTACACCACTGAGAGCGCGGTGGTTGACCAGACAAACCTCTA<br>TTTCAAAGTCGTTAAAAGGGTTGTTCAAGTCAGGAGGGGGAAACGCCAATACTGGCCCTACTACTGCTGCTGCTGCTGC<br>TGCACCATCTAGTATCTTTTAAATTAATAATGTAATTCATATATTATAGTGCGGCCGCGTGTTACAACCAATTAACCAATTC<br>TGATTAGAAAAACTCATCGAGCATCAAATGAAACTGCAATTTATTCATATCAGGATTATCAATACCATATTTTTGAAAAAGC<br>CGTTTCTGTAATGAAGGAGAAAACTCACCGAGGCAGTTCCATAGGATGGCAAGATCCTGGTATCGGTCTGCGATTCCGA<br>CTCGTCCAACATCAATACAACCTATTAATTTCCCCTCGTCAAAAATAAGGTTATCAAGTGAGAAATCACCATGAGTGACGA<br>CTGAATCCGGTGAGAATGGCAAAAGCTTATGCATTTCTTTCCAGACTTGTTCAACAGGCCAGCCATTACGCTCGTCATCA<br>AAATCACTCGCATCAACCAAACCGTTATTCATTCGTGATTGCGCCTGAGCGAGGCGAAATACGCGATCGCTGTTAAAAG<br>GACAATTACAAACAGGAATCGAATGCAACCGGCGCAGGAACACTGCCAGCGCATCAACAATATTTTCACCTGAATCAGG<br>ATATTCTTCTAATACCTGGAATGCTGTTTTCCCGGGGATCGCAGTGGTGAGTAACCATGCATCATCAGGAGTACGGATAA<br>AATGCTTGATGGTCGGAAGAGGCATAAATTCCGTCAGCCAGTTTAGTCTGACCATCTCATCTGTAACATCATTGGCAACG<br>CTACCTTTGCCATGTTTCAGAAACAACTCTGGCGCATCGGGCTTCCCATACAATCGATAGATTGTCGCACCTGATTGCCC<br>GACATTATCGCGAGCCCATTTATACCCATATAAATCAGCATCCATGTTGGAATTTAATCGCGGCCTGGAGCAAGACGTTT<br>CCCGTTGAATATGGCTCATAACACCCCTTGTATTACTGTTTATGTAAGCAGACAGTTTTATTGTTCATGATGATATATTTTT<br>ATCTTGTGCAATGTAACATCAGAGATTTTGAGACACAACGTGGCTTTGTTGAATAAATCGAACTTTTGCTGAGTTGAAGG<br>ATCAGTCATGACCAAAATCCCTTAACGTGAGTTTTCGTTCCACTGAGCGTCAGACCCCGTAGAAAAGATCAAAGGATCTT<br>CTTGAGATCCTTTTTTTCTGCGCGTAATCTGCTGCTTGCAAACAAAAAAACCACCGCTACCAGCGGTGGTTTGTTTGCCG<br>GATCAAGAGCTACCAACTCTTTTTCCGAAGGTAACTGGCTTCAGCAGAGCGCAGATACCAAATACTGTTCTTCTAGTGTA<br>GCCGTAGTTAGGCCACCACTTCAAGAACTCTGTAGCACCGCCTACATACCTCGCTCTGCTAATCCTGTTACCAGTGGCT<br>GCTGCCAGTGGCGATAAGTCGTGTCTTACCGGGTTGGACTCAAGACGATAGTTACCGGATAAGGCGCAGCGGTCGGGC<br>TGAACGGGGGGTTCGTGCACACAGCCCAGCTTGGAGCGAACGACCTACACCGAACTGAGATACCTACAGCGTGAGCTA<br>TGAGAAAGCGCCACGCTTCCCGAAGGGAGAAAGGCGGACAGGTATCCGGTAAGCGGCAGGGTCGGAACAGGAGAGC<br>GCACGAGGGAGCTTCCAGGGGGAAACGCCTGGTATCTTTATAGTCCTGTCGGGTTTCGCCACCTCTGACTTGAGCGTC<br>GATTTTTGTGATGCTCGTCAGGGGGGCGGAGCCTATGGAAAAACGCCAGCAACGCGGCCTTTTTACGGTTCCTGGCCT<br>TTTGCTGGCCTTTTGCTCACATGTTCTTTCCTGCGTTATCCCCTGATTCTGTGGATAACCGTGCGGCCGC |
| pRW403 | Plasmid for genertaion of donor DNA for PhiC31-compatible p1 integration in yeast. After linearization with ScaI, this plasmid produces donor DNA encoding the fluorescent protein mScarletI as a reporter with a *LEU2* auxotrophic selection marker, flanked by *attP* recombination sites for PhiC31. This plasmid contains AmpR marker for propagation in *E. coli.* | ACTATCTATTTTTTCAGTACATAATTTAATTAAATCATCATGACCTTTATCTAATTTTACATCTCTTCTACCAAGACCTAATT<br>CTTTATTAATACTATCAAAATCTGTTATTCTATTGCCTTGAGGATCTTTGAAACACCAATCTGAGTCTGAGGGCCCCAACT<br>GGGGTAACCTCGGAGTTCTCTCAGTTGGGGGTCTGATGAGACGGAGAGGTCAGCTTTTCTTTGCTAATATCCAGTGAAC<br>TACCAATCATTATAGGAAGTTCTCTGTCATTGTATATAGCACAGTAACTTAGGATTTAAGAACCTCTAGAACCAGTAGAAC<br>CAGGACCAGCAGTACCATCAACAGCAGAATTAGAAGCTTGAGCCCTAGATCTCAAACCAGATTTATACAATTCATCCATA<br>CCACCAGTAGAATGTCTACCTTCAGATCTTTCATATTGTTCAACAACAGTATAATCCTCGTTATGTGAAGTAATGTCAAGT<br>TTTCTATCTACATTATAAGCACCCGGCATTTGAACAGGTTTTTAGCTTTATAAGTAGTTTTAAAATCAGCCAAATATCTAC<br>CACCATCTTTCAATCTCAAAGCCATTTTAATATCACCTTTCAAAACACCATCTTCAGGATACAATCTTTCAGTAGAAGCTTC<br>CCAACCCATAGTTTTTTTTGCATAACAGGACCATCAGGTGGAAAATTAGTACCTCTCAATTTAACTTTATAAATCAAAGTA<br>CCATCTTCCAAAGAAGTATCTTGAGTAACAGTAACAGCACCACCATCTTCAAAATTCATAACTCTTTCCCATTTAAAACCT<br>TCAGGAAAAGATTGTTTATAATAATCAGGAATATCAGCAGGATGTTTAATAAAAGCTCTAGAACCATACATAAATTGTGGA<br>GACAAAATATCCCAAGAAAATGGCAATGGACCACCTTTAGTAACTTTCAATTTAGCAGTTTGAGTACCTTCATAAGGTCTA<br>CCTTCACCTTCACCTTCAATTTCAAATTCATGACCATTCATAGAACCTTCCATGTGAACTTTAAATCTCATAAATTCTTTAA<br>TAACAGCTTCACCTTTAGAAACCATATTTTTATATGTCTATGAGCTTATCACATGTTTCTATATTTTTCTGATATTTGAAAGT<br>TATTAATCTTTTTGTTTCTATAGATCTTCCTATGTATAGGTCATCCGTTATCGTCTCAAGCAGTACAGAAAATTAATTTCTA<br>ATGTGAACACTAGTGAGATAGCTAAAGATCTTAATTGTGAAAACAATATTGAAAGTATAATAAATACATTAAAAGAACAAAA<br>TAGATATTTTGACAAACAAATTGAATATATGTCTGCCCCTAAGAAGATCGTCGTTTTGCCAGGTGACCACGTTGGTCAAG<br>AAATCACAGCCGAAGCCATTAAGGTTCTTAAAGCTATTTCTGATGTTCGTTCCAATGTCAAGTTCGATTTCGAAAATCATT<br>TAATTGGTGGTGCTGCTATCGATGCTACAGGTGTTCCACTTCCAGATGAGGCGCTGGAAGCCTCCAAGAAGGCTGATGC<br>CGTTTTGTTAGGTGCTGTGGGTGGTCCTAAATGGGGAACCGGTAGTGTTAGACCTGAACAAGGTTTACTAAAAATCCGT<br>AAAGAACTTCAATTGTACGCCAACTTAAGACCATGTAACTTTGCATCCGACTCTCTTTTAGACTTATCTCCAATCAAGCCA<br>CAATTTGCTAAAGGTACTGACTTCGTTGTTGTCAGAGAATTAGTGGGAGGTATTTACTTTGGTAAGAGAAAGGAGGACGA<br>TGGTGATGGTGTCGCTTGGGATAGTGAACAATACACCGTTCCAGAAGTGCAAAGAATCACAAGAATGGCCGCTTTCATG<br>GCCCTACAACATGAGCCACCATTGCCTATTTGGTCCTTGGATAAAGCTAATGTTTTGGCCTCTTCAAGATTATGGAGAAA<br>AACTGTGGAGGAAACCATCAAGAACGAATTTCCTACATTGAAGGTTCAACATCAATTGATTGATTCTGCCGCCATGATCC<br>TAGTTAAGAACCCAACCCACCTAAATGGTATTATAATCACCAGCAACATGTTTGGTGATATCATCTCCGATGAAGCCTCC<br>GTTATCCCAGGTTCCTTGGGTTTGTTGCCATCTGCGTCCTTGGCCTCTTTGCCAGACAAGAACACCGCATTTGGTTTGTA<br>CGAACCATGCCACGGTTCTGCTCCAGATTTGCCAAAGAATAAGGTCAACCCTATCGCCACTATCTTGTCTGCTGCAATGA<br>TGTTGAAATTGTCATTGAACTTGCCTGAAGAAGGTAAGGCCATTGAAGATGCAGTTAAAAAGGTTTTGGATGCAGGTATC<br>AGAACTGGTGATTTAGGTGGTTCCAACAGTACCACCGAAGTCGGTGATGCTGTCGCCGAAGAAGTTAAGAAAATCCTTG<br>CTTAAAAGAGAGTGAGCCCCCAACTGAGAGAACTCGTAGGTTACCCCAGTTGGGGCTCTATTTCAAAGTCGTTAAAAGG<br>GTTGTTCAAGTCAGGAGGGGGAAACGCCAATACTGGCCCTACTACTGCTGCTGCTGCTGCTGCACCATCTAGTATCTTT<br>TAAATTAATAATGTAATTCATATATTATAGTGCGGCCGCGTGTTACAACCAATTAACCAATTCTGATTAGAAAAACTCATCG<br>AGCATCAAATGAAACTGCAATTTATTCATATCAGGATTATCAATACCATATTTTTGAAAAAGCCGTTTCTGTAATGAAGGA<br>GAAAACTCACCGAGGCAGTTCCATAGGATGGCAAGATCCTGGTATCGGTCTGCGATTCCGACTCGTCCAACATCAATAC<br>AACCTATTAATTTCCCCTCGTCAAAAATAAGGTTATCAAGTGAGAAATCACCATGAGTGACGACTGAATCCGGTGAGAAT<br>GGCAAAAGCTTATGCATTTCTTTCCAGACTTGTTCAACAGGCCAGCCATTACGCTCGTCATCAAAATCACTCGCATCAAC<br>CAAACCGTTATTCATTCGTGATTGCGCCTGAGCGAGGCGAAATACGCGATCGCTGTTAAAAGGACAATTACAAACAGGA<br>ATCGAATGCAACCGGCGCAGGAACACTGCCAGCGCATCAACAATATTTTCACCTGAATCAGGATATTCTTCTAATACCTG<br>GAATGCTGTTTTCCCGGGGATCGCAGTGGTGAGTAACCATGCATCATCAGGAGTACGGATAAAATGCTTGATGGTCGGA<br>AGAGGCATAAATTCCGTCAGCCAGTTTAGTCTGACCATCTCATCTGTAACATCATTGGCAACGCTACCTTTGCCATGTTT<br>CAGAAACAACTCTGGCGCATCGGGCTTCCCATACAATCGATAGATTGTCGCACCTGATTGCCCGACATTATCGCGAGCC<br>CATTTATACCCATATAAATCAGCATCCATGTTGGAATTTAATCGCGGCCTGGAGCAAGACGTTTCCCGTTGAATATGGCT<br>CATAACACCCCTTGTATTACTGTTTATGTAAGCAGACAGTTTTATTGTTCATGATGATATATTTTTATCTTGTGCAATGTAA<br>CATCAGAGATTTTGAGACACAACGTGGCTTTGTTGAATAAATCGAACTTTTGCTGAGTTGAAGGATCAGTCATGACCAAA<br>ATCCCTTAACGTGAGTTTTCGTTCCACTGAGCGTCAGACCCCGTAGAAAAGATCAAAGGATCTTCTTGAGATCCTTTTTTT<br>CTGCGCGTAATCTGCTGCTTGCAAACAAAAAAACCACCGCTACCAGCGGTGGTTTGTTTGCCGGATCAAGAGCTACCAA<br>CTCTTTTTCCGAAGGTAACTGGCTTCAGCAGAGCGCAGATACCAAATACTGTTCTTCTAGTGTAGCCGTAGTTAGGCCAC<br>CACTTCAAGAACTCTGTAGCACCGCCTACATACCTCGCTCTGCTAATCCTGTTACCAGTGGCTGCTGCCAGTGGCGATA<br>AGTCGTGTCTTACCGGGTTGGACTCAAGACGATAGTTACCGGATAAGGCGCAGCGGTCGGGCTGAACGGGGGGTTCG<br>TGCACACAGCCCAGCTTGGAGCGAACGACCTACACCGAACTGAGATACCTACAGCGTGAGCTATGAGAAAGCGCCACG<br>CTTCCCGAAGGGAGAAAGGCGGACAGGTATCCGGTAAGCGGCAGGGTCGGAACAGGAGAGCGCACGAGGGAGCTTC<br>CAGGGGGAAACGCCTGGTATCTTTATAGTCCTGTCGGGTTTCGCCACCTCTGACTTGAGCGTCGATTTTTGTGATGCTC<br>GTCAGGGGGGCGGAGCCTATGGAAAAACGCCAGCAACGCGGCCTTTTTACGGTTCCTGGCCTTTTGCTGGCCTTTTGC<br>TCACATGTTCTTTCCTGCGTTATCCCCTGATTCTGTGGATAACCGTGCGGCCGC |
| pRW404 | 2μ plasmid for constitutive expression of BXB1 under RNR2 promoter. The plasmid includes a kanMX marker for | GTTGCTGGCGTTTTTCCATAGGCTCCGCCCCCCTGACGAGCATCACAAAAATCGACGCTCAAGTCAGAGGTGGCGAAA<br>CCCGACAGGACTATAAAGATACCAGGCGTTTCCCCCTGGAAGCTCCCTCGTGCGCTCTCCTGTTCCGACCCTGCCGCT<br>TACCGGATACCTGTCCGCCTTTCTCCCTTCGGGAAGCGTGGCGCTTTCTCATAGCTCACGCTGTAGGTATCTCAGTTCG<br>GTGTAGGTCGTTCGCTCCAAGCTGGGCTGTGTGCACGAACCCCCCGTTCAGCCCGACCGCTGCGCCTTATCCGGTAAC<br>TATCGTCTTGAGTCCAACCCGGTAAGACACGACTTATCGCCACTGGCAGCAGCCACTGGTAACAGGATTAGCAGAGCG<br>AGGTATGTAGGCGGTGCTACAGAGTTCTTGAAGTGGTGGCCTAACTACGGCTACACTAGAAGGACAGTATTTGGTATCT<br>GCGCTCTGCTGAAGCCAGTTACCTTCGGAAAAAGAGTTGGTAGCTCTTGATCCGGCAAACAAACCACCGCTGGTAGCG<br>GTGGTTTTTTTGTTTGCAAGCAGCAGATTACGCGCAGAAAAAAAGGATCTCAAGAAGATCCTTTGATCTTTTCTACACTAG<br>TCGAAGCATCTGTGCTTCATTTTGTAGAACAAAAATGCAACGCGAGAGCGCTAATTTTTCAAACAAAGAATCTGAGCTGC<br>ATTTTTACAGAACAGAAATGCAACGCGAAAGCGCTATTTTACCAACGAAGAATCTGTGCTTCATTTTTGTAAAACAAAAAT<br>GCAACGCGAGAGCGCTAATTTTTCAAACAAAGAATCTGAGCTGCATTTTTACAGAACAGAAATGCAACGCGAGAGCGCT<br>ATTTTACCAACAAAGAATCTATACTTCTTTTTTGTTCTACAAAAATGCATCCCGAGAGCGCTATTTTTCTAACAAAGCATCT<br>TAGATTACTTTTTTTCTCCTTTGTGCGCTCTATAATGCAGTCTCTTGATAACTTTTTGCACTGTAGGTCCGTTAAGGTTAGA |

| | | |
|---|---|---|
| | selection in yeast and propagation in *E.coli*. | AGAAGGCTACTTTGGTGTCTATTTTCTCTTCCATAAAAAAAGCCTGACTCCACTTCCCGCGTTTACTGATTACTAGCGAAG<br>CTGCGGGTGCATTTTTTCAAGATAAAGGCATCCCCGATTATATTCTATACCGATGTGGATTGCGCATACTTTGTGAACAG<br>AAAGTGATAGCGTTGATGATTCTTCATTGGTCAGAAAATTATGAACGGTTTCTTCTATTTTGTCTCTATATACTACGTATAG<br>GAAATGTTTACATTTTCGTATTGTTTTCGATTCACTCTATGAATAGTTCTTACTACAATTTTTTTGTCTAAAGAGTAATACTA<br>GAGATAAACATAAAAAATGTAGAGGTCGAGTTTAGATGCAAGTTCAAGGAGCGAAAGGTGGATGGGTAGGTTATATAGG<br>GATATAGCACAGAGATATATAGCAAAGAGATACTTTTGAGCAATGTTTGTGGAAGCGGTATTCGCAATATTTTAGTAGCT<br>CGTTACAGTCCGGTGCGTTTTTGGTTTTTTGAAAGTGCGTCTTCAGAGCGCTTTTGGTTTTCAAAAGCGCTCTGAAGTTC<br>CTATACTTTCTAGAGAATAGGAACTTCGGAATAGGAACTTCAAAGCGTTTCCGAAAACGAGCGCTTCCGAAAATGCAACG<br>CGAGCTGCGCACATACAGCTCACTGTTCACGTCGCACCTATATCTGCGTGTTGCCTGTATATATATATACATGAGAAGAA<br>CGGCATAGTGCGTGTTTATGCTTAAATGCGTATATGTGTTATGTATAGGTCTAGAGATCTGTTTAGCTTGCCTCGTCCCC<br>GCCGGGTCACCCGGCCAGCGACATGGAGGCCCAGAATACCCTCCTTGACAGTCTTGACGTGCGCAGCTCAGGGGCAT<br>GATGTGACTGTCGCCCGTACATTTAGCCCATACATCCCCATGTATAATCATTTGCATCCATACATTTTGATGGCCGCACG<br>GCGCGAAGCAAAAATTACGGCTCCTCGCTGCAGACCTGCGAGCAGGGAAACGCTCCCCTCACAGACGCGTTGAATTGT<br>CCCCACGCCGCGCCCCTGTAGAGAAATATAAAAGGTTAGGATTTGCCACTGAGGTTCTTCTTTCATATACTTCCTTTTAA<br>AATCTTGCTAGGATACAGTTCTCACATCACATCCGAACATAAACAACCATGGGTAAGGAAAAGACTCACGTTTCGAGGCC<br>GCGATTAAATTCCAACATGGATGCTGATTTATATGGGTATAAATGGGCTCGCGATAATGTCGGGCAATCAGGTGCGACA<br>ATCTATCGATTGTATGGGAAGCCCGATGCGCCAGAGTTGTTTCTGAAACATGGCAAAGGTAGCGTTGCCAATGATGTTA<br>CAGATGAGATGGTCAGACTAAACTGGCTGACGGAATTTATGCCTCTTCCGACCATCAAGCATTTTATCCGTACTCCTGAT<br>GATGCATGGTTACTCACCACTGCGATCCCCGGCAAAACAGCATTCCAGGTATTAGAAGAATATCCTGATTCAGGTGAAA<br>ATATTGTTGATGCGCTGGCAGTGTTCCTGCGCCGGTTGCATTCGATTCCTGTTTGTAATTGTCCTTTTAACAGCGATCGC<br>GTATTTCGTCTCGCTCAGGCGCAATCACGAATGAATAACGGTTTGGTTGATGCGAGTGATTTTGATGACGAGCGTAATG<br>GCTGGCCTGTTGAACAAGTCTGGAAAGAAATGCATAAGCTTTTGCCATTCTCACCGGATTCAGTCGTCACTCATGGTGAT<br>TTCTCACTTGATAACCTTATTTTTGACGAGGGGAAATTAATAGGTTGTATTGATGTTGGACGAGTCGGAATCGCAGACCG<br>ATACCAGGATCTTGCCATCCTATGGAACTGCCTCGGTGAGTTTTCTCCTTCATTACAGAAACGGCTTTTTCAAAAATATG<br>GTATTGATAATCCTGATATGAATAAATTGCAGTTTCATTTGATGCTCGATGAGTTTTTCTAATCAGTACTGACAATAAAAAG<br>ATTCTTGTTTTCAAGAACTTGTCATTTGTATAGTTTTTTTATATTGTAGTTGTTCTATTTTAATCAAATGTTAGCGTGATTTAT<br>ATTTTTTTTCGCCTCGACATCATCTGCCCAGATGCGAAGTTAAGTGCGCAGAAAGTAATATCATGCGTCAATCGTATGTG<br>AATGCTGGTCGCTATACTGCTGTCGATTCGATACTAACGCCGCCATCCAGTGTCGAAAACGAGCTCTCGAGAACCCTTA<br>ATGTCGACAGTCGAACAAGAAGCAGGCAAAGTTTAGAGCACTGCCCCTCCGCACTCAAAAAAGAAAAAACTAGGAGGAA<br>AATAAAATTCTCAACCACACAAACACATAAACACATACAAATACAAATACAAGCTTATTTACTTGACATCGCGCGATCTTC<br>CACTATTCAGCGCCGTCCGCCCTCTCTCGTGTTTTTTGTTTACGCGACAACTATGCGAAATCCGGAGCAACGGGCAACC<br>GTTTGGGGAAAGACCACACCCACGCGCGATCGCCATGGCAACGAGGTCGCACACGCCCCACACCCAGACCTCCCTGC<br>GAGCGGGCATGGGTACAATGTCCCCGTTGCCACAGACACCACTTCGTAGCACAGCGCAGAGCGTAGCGTGTTGTTGCT<br>GCTGACAAAAGAAAATTTTTCTTAGCAAAGCAAAGGAGGGGAAGCACGGGCAGATAGCACCGTACCATACCCTTGGAAA<br>CTCGAAATGAACGAAGCAGGAAATGAGAGAATGAGAGTTTTGTAGGTATATATAGCGGTAGTGTTTGCGCGTTACCATC<br>ATCTTCTGGATCTATCTATTGTTCTTTTCCTCATCACTTTCCCCTTTTTCGCTCTTCTTCTTGTCTTTTATTTCTTTCTTTTTT<br>TTAATTGTTCCCTCGATTGGCTATCTACCAAAGAATCCAAACTTAATACACGTATTTATTTGTCCAATTACCATGGTGAGA<br>GCCCTGGTAGTCATCCGCCTGTCCCGCGTCACCGATGCTACGACTTCACCGGAGCGTCAGCTGGAGTCTTGCCAGCAG<br>CTCTGCGCCCAGCGCGGCTGGGACGTCGTCGGGGTAGCGGAGGATCTGGACGTCTCCGGGGCGGTCGATCCGTTCG<br>ACCGGAAGCGCAGACCGAACCTGGCCCGGTGGCTAGCGTTCGAGGAGCAACCGTTTGACGTGATCGTGGCGTACCGG<br>GTAGATCGGTTGACCCGATCGATCCGGCATCTTCAGCAGCTGGTCCACTGGGCCGAGGACCACAAGAAGCTGGTCGTC<br>TCCGCGACCGAAGCGCACTTCGATACGACGACGCCGTTTGCGGCGGTCGTCATCGCGCTTATGGGAACGGTGGCGCA<br>GATGGAATTAGAAGCGATCAAAGAGCGGAACCGTTCGGCTGCGCATTTCAATATCCGCGCCGGGAAATACCGAGGATC<br>CCTGCCGCCGTGGGGATACCTGCCTACGCGCGTGGACGGGGAGTGGCGGCTGGTGCCGGACCCTGTGCAGCGAGAG<br>CGCATCCTCGAGGTGTATCACCGCGTCGTCGACAACCACGAGCCGCTGCATCTGGTGGCCCACGACCTGAACCGGCG<br>TGGTGTCCTGTCGCCGAAGGACTACTTCGCGCAGCTGCAAGGCCGCGAGCCGCAGGGCCGGGAGTGGTCGGCTACC<br>GCGCTGAAGCGATCGATGATCTCCGGGGCGATGCTCGGGTACGCGACTCTGAACGGTAAGACCGTCCGAGACGACGA<br>CGGAGCCCCGCTGGTGCGGGCTGAGCCGATCCTGACCCGTGAGCAGCTGGAGGCGCTGCGCGCCGAGCTCGTCAAG<br>ACCTCCCGGGCGAAGCCCGCGGTGTCTACCCCGTCGCTGCTGCTGCGGGTGTTGTTCTGCGCGGTGTGCGGGGAGCC<br>CGCGTACAAGTTCGCCGGGGGAGGACGTAAGCACCCGCGCTACCGCTGCCGCTCGATGGGGTTCCCGAAGCACTGCG<br>GGAACGGCACGGTGGCGATGGCCGAGTGGGACGCGTTCTGCGAGGAGCAGGTACTGGATCTGCTCGGGGACGCGGA<br>GCGTCTGGAGAAAGTCTGGGTAGCGGGCTCGGACTCCGCGGTCGAACTCGCGGAGGTGAACGCGGAGCTGGTGGAC<br>CTGACGTCGCTGATCGGCTCCCCGGCCTACCGGGCGGGCTCTCCGCAGCGAGAAGCACTGGATGCCCGTATTGCGGC<br>GCTGGCCGCGCGGCAAGAGGAGCTGGAGGGCCTGGAGGCTCGCCCGTCTGGCTGGGAGTGGCGCGAAACCGGGCA<br>GCGGTTCGGGGACTGGTGGCGGGAGCAGGACACCGCGGCAAAGAACACCTGGCTTCGGTCGATGAACGTTCGGCTGA<br>CGTTCGACGTCCGCGGCGGGCTGACTCGCACGATCGACTTCGGGGATCTTCAGGAGTACGAGCAGCATCTCAGGCTC<br>GGCAGCGTGGTCGAACGGCTACACACCGGGATGTCGTAATAACTCGAGGCGAATTTCTTATGATTTATGATTTTTATTAT<br>TAAATAAGTTATAAAAAAAATAAGTGTATACAAATTTTAAAGTGACTCTTAGGTTTTAAAACGAAAATTCTTATTCTTGAGTA<br>ACTCTTTCCTGTAGGTCAGGTTGCTTTCTCAGGTATAGCATGAGGTCGCTCTTATTGACCACACCTCTACCGGCATGCCG<br>AGCAAATGCCTGCAAATCGCTCCCCATTTCAGCAAAAGGCCAGCAAAAGGCCAGGAACCGTAAAAAGGCCGC |
| pRW405 | 2*μ* plasmid for constitutive expression of PhiC31 under RNR2 promoter. The plasmid includes a kanMX marker for selection in yeast and propagation in *E.coli*. | GTTGCTGGCGTTTTTCCATAGGCTCCGCCCCCCTGACGAGCATCACAAAAATCGACGCTCAAGTCAGAGGTGGCGAAA<br>CCCGACAGGACTATAAAGATACCAGGCGTTTCCCCCTGGAAGCTCCCTCGTGCGCTCTCCTGTTCCGACCCTGCCGCT<br>TACCGGATACCTGTCCGCCTTTCTCCCTTCGGGAAGCGTGGCGCTTTCTCATAGCTCACGCTGTAGGTATCTCAGTTCG<br>GTGTAGGTCGTTCGCTCCAAGCTGGGCTGTGTGCACGAACCCCCCGTTCAGCCCGACCGCTGCGCCTTATCCGGTAAC<br>TATCGTCTTGAGTCCAACCCGGTAAGACACGACTTATCGCCACTGGCAGCAGCCACTGGTAACAGGATTAGCAGAGCG<br>AGGTATGTAGGCGGTGCTACAGAGTTCTTGAAGTGGTGGCCTAACTACGGCTACACTAGAAGGACAGTATTTGGTATCT<br>GCGCTCTGCTGAAGCCAGTTACCTTCGGAAAAAGAGTTGGTAGCTCTTGATCCGGCAAACAAACCACCGCTGGTAGCG<br>GTGGTTTTTTTGTTTGCAAGCAGCAGATTACGCGCAGAAAAAAAGGATCTCAAGAAGATCCTTTGATCTTTTCTACACTAG<br>TCGAAGCATCTGTGCTTCATTTTGTAGAACAAAAATGCAACGCGAGAGCGCTAATTTTTCAAACAAAGAATCTGAGCTGC<br>ATTTTTACAGAACAGAAATGCAACGCGAAAGCGCTATTTTACCAACGAAGAATCTGTGCTTCATTTTTGTAAAACAAAAAT<br>GCAACGCGAGAGCGCTAATTTTTCAAACAAAGAATCTGAGCTGCATTTTTACAGAACAGAAATGCAACGCGAGAGCGCT<br>ATTTTACCAACAAAGAATCTATACTTCTTTTTTGTTCTACAAAAATGCATCCCGAGAGCGCTATTTTTCTAACAAAGCATCT<br>TAGATTACTTTTTTTCTCCTTTGTGCGCTCTATAATGCAGTCTCTTGATAACTTTTTGCACTGTAGGTCCGTTAAGGTTAGA<br>AGAAGGCTACTTTGGTGTCTATTTTCTCTTCCATAAAAAAAGCCTGACTCCACTTCCCGCGTTTACTGATTACTAGCGAAG<br>CTGCGGGTGCATTTTTTCAAGATAAAGGCATCCCCGATTATATTCTATACCGATGTGGATTGCGCATACTTTGTGAACAG<br>AAAGTGATAGCGTTGATGATTCTTCATTGGTCAGAAAATTATGAACGGTTTCTTCTATTTTGTCTCTATATACTACGTATAG<br>GAAATGTTTACATTTTCGTATTGTTTTCGATTCACTCTATGAATAGTTCTTACTACAATTTTTTTGTCTAAAGAGTAATACTA<br>GAGATAAACATAAAAAATGTAGAGGTCGAGTTTAGATGCAAGTTCAAGGAGCGAAAGGTGGATGGGTAGGTTATATAGG<br>GATATAGCACAGAGATATATAGCAAAGAGATACTTTTGAGCAATGTTTGTGGAAGCGGTATTCGCAATATTTTAGTAGCT<br>CGTTACAGTCCGGTGCGTTTTTGGTTTTTTGAAAGTGCGTCTTCAGAGCGCTTTTGGTTTTCAAAAGCGCTCTGAAGTTC<br>CTATACTTTCTAGAGAATAGGAACTTCGGAATAGGAACTTCAAAGCGTTTCCGAAAACGAGCGCTTCCGAAAATGCAACG<br>CGAGCTGCGCACATACAGCTCACTGTTCACGTCGCACCTATATCTGCGTGTTGCCTGTATATATATATACATGAGAAGAA<br>CGGCATAGTGCGTGTTTATGCTTAAATGCGTATATGTGTTATGTATAGGTCTAGAGATCTGTTTAGCTTGCCTCGTCCCC<br>GCCGGGTCACCCGGCCAGCGACATGGAGGCCCAGAATACCCTCCTTGACAGTCTTGACGTGCGCAGCTCAGGGGCAT<br>GATGTGACTGTCGCCCGTACATTTAGCCCATACATCCCCATGTATAATCATTTGCATCCATACATTTTGATGGCCGCACG<br>GCGCGAAGCAAAAATTACGGCTCCTCGCTGCAGACCTGCGAGCAGGGAAACGCTCCCCTCACAGACGCGTTGAATTGT<br>CCCCACGCCGCGCCCCTGTAGAGAAATATAAAAGGTTAGGATTTGCCACTGAGGTTCTTCTTTCATATACTTCCTTTTAA<br>AATCTTGCTAGGATACAGTTCTCACATCACATCCGAACATAAACAACCATGGGTAAGGAAAAGACTCACGTTTCGAGGCC<br>GCGATTAAATTCCAACATGGATGCTGATTTATATGGGTATAAATGGGCTCGCGATAATGTCGGGCAATCAGGTGCGACA<br>ATCTATCGATTGTATGGGAAGCCCGATGCGCCAGAGTTGTTTCTGAAACATGGCAAAGGTAGCGTTGCCAATGATGTTA<br>CAGATGAGATGGTCAGACTAAACTGGCTGACGGAATTTATGCCTCTTCCGACCATCAAGCATTTTATCCGTACTCCTGAT<br>GATGCATGGTTACTCACCACTGCGATCCCCGGCAAAACAGCATTCCAGGTATTAGAAGAATATCCTGATTCAGGTGAAA<br>ATATTGTTGATGCGCTGGCAGTGTTCCTGCGCCGGTTGCATTCGATTCCTGTTTGTAATTGTCCTTTTAACAGCGATCGC<br>GTATTTCGTCTCGCTCAGGCGCAATCACGAATGAATAACGGTTTGGTTGATGCGAGTGATTTTGATGACGAGCGTAATG<br>GCTGGCCTGTTGAACAAGTCTGGAAAGAAATGCATAAGCTTTTGCCATTCTCACCGGATTCAGTCGTCACTCATGGTGAT<br>TTCTCACTTGATAACCTTATTTTTGACGAGGGGAAATTAATAGGTTGTATTGATGTTGGACGAGTCGGAATCGCAGACCG<br>ATACCAGGATCTTGCCATCCTATGGAACTGCCTCGGTGAGTTTTCTCCTTCATTACAGAAACGGCTTTTTCAAAAATATG |

| | | |
|---|---|---|
| | | GTATTGATAATCCTGATATGAATAAATTGCAGTTTCATTTGATGCTCGATGAGTTTTTCTAATCAGTACTGACAATAAAAAG<br>ATTCTTGTTTTCAAGAACTTGTCATTTGTATAGTTTTTTTATATTGTAGTTGTTCTATTTTAATCAAATGTTAGCGTGATTTAT<br>ATTTTTTTTCGCCTCGACATCATCTGCCCAGATGCGAAGTTAAGTGCGCAGAAAGTAATATCATGCGTCAATCGTATGTG<br>AATGCTGGTCGCTATACTGCTGTCGATTCGATACTAACGCCGCCATCCAGTGTCGAAAACGAGCTCTCGAGAACCCTTA<br>ATGTCGACAGTCGAACAAGAAGCAGGCAAAGTTTAGAGCACTGCCCCTCCGCACTCAAAAAAGAAAAAACTAGGAGGAA<br>AATAAAATTCTCAACCACACAAACACATAAACACATACAAATACAAATACAAGCTTATTTACTTGACATCGCGCGATCTTC<br>CACTATTCAGCGCCGTCCGCCCTCTCTCGTGTTTTTTGTTTACGCGACAACTATGCGAAATCCGGAGCAACGGGCAACC<br>GTTTGGGGAAAGACCACACCCACGCGCGATCGCCATGGCAACGAGGTCGCACACGCCCCACACCCAGACCTCCCTGC<br>GAGCGGGCATGGGTACAATGTCCCCGTTGCCACAGACACCACTTCGTAGCACAGCGCAGAGCGTAGCGTGTTGTTGCT<br>GCTGACAAAAGAAAATTTTTCTTAGCAAAGCAAAGGAGGGGAAGCACGGGCAGATAGCACCGTACCATACCCTTGGAAA<br>CTCGAAATGAACGAAGCAGGAAATGAGAGAATGAGAGTTTTGTAGGTATATATAGCGGTAGTGTTTGCGCGTTACCATC<br>ATCTTCTGGATCTATCTATTGTTCTTTTCCTCATCACTTTCCCCTTTTTCGCTCTTCTTCTTGTCTTTTATTTCTTTCTTTTTT<br>TTAATTGTTCCCTCGATTGGCTATCTACCAAAGAATCCAAACTTAATACACGTATTTATTTGTCCAATTACCATGGACACG<br>TACGCGGGTGCTTACGACCGTCAGTCGCGCGAGCGCGAGAATTCGAGCGCAGCAAGCCCAGCGACACAGCGTAGCGC<br>CAACGAAGACAAGGCGGCCGACCTTCAGCGCGAAGTCGAGCGCGACGGGGGCCGGTTCAGGTTCGTCGGGCATTTCA<br>GCGAAGCGCCGGGCACGTCGGCGTTCGGGACGGCGGAGCGCCCGGAGTTCGAACGCATCCTGAACGAATGCCGCGC<br>CGGGCGGCTCAACATGATCATTGTCTATGACGTGTCGCGCTTCTCGCGCCTGAAGGTCATGGACGCGATTCCGATTGT<br>CTCGGAATTGCTCGCCCTGGGCGTGACGATTGTTTCCACTCAGGAAGGCGTCTTCCGGCAGGGAAACGTCATGGACCT<br>GATTCACCTGATTATGCGGCTCGACGCGTCGCACAAAGAATCTTCGCTGAAGTCGGCGAAGATTCTCGACACGAAGAAC<br>CTTCAGCGCGAATTGGGCGGGTACGTCGGCGGGAAGGCGCCTTACGGCTTCGAGCTTGTTTCGGAGACGAAGGAGAT<br>CACGCGCAACGGCCGAATGGTCAATGTCGTCATCAACAAGCTTGCGCACTCGACCACTCCCCTTACCGGACCCTTCGA<br>GTTCGAGCCCGACGTAATCCGGTGGTGGTGGCGTGAGATCAAGACGCACAAACACCTTCCCTTCAAGCCGGGCAGTCA<br>AGCCGCCATTCACCCGGGCAGCATCACGGGGCTTTGTAAGCGCATGGACGCTGACGCCGTGCCGACCCGGGGCGAGA<br>CGATTGGGAAGAAGACCGCTTCAAGCGCCTGGGACCCGGCAACCGTTATGCGAATCCTTCGGGACCCGCGTATTGCG<br>GGCTTCGCCGCTGAGGTGATCTACAAGAAGAAGCCGGACGGCACGCCGACCACGAAGATTGAGGGTTACCGCATTCA<br>GCGCGACCCGATCACGCTCCGGCCGGTCGAGCTTGATTGCGGACCGATCATCGAGCCCGCTGAGTGGTATGAGCTTC<br>AGGCGTGGTTGGACGGCAGGGGGCGCGGCAAGGGGCTTTCCCGGGGGCAAGCCATTCTGTCCGCCATGGACAAGCT<br>GTACTGCGAGTGTGGCGCCGTCATGACTTCGAAGCGCGGGGAAGAATCGATCAAGGACTCTTACCGCTGCCGTCGCC<br>GGAAGGTGGTCGACCCGTCCGCACCTGGGCAGCACGAAGGCACGTGCAACGTCAGCATGGCGGCACTCGACAAGTTC<br>GTTGCGGAACGCATCTTCAACAAGATCAGGCACGCCGAAGGCGACGAAGAGACGTTGGCGCTTCTGTGGGAAGCCGC<br>CCGACGCTTCGGCAAGCTCACTGAGGCGCCTGAGAAGAGCGGCGAACGGGCGAACCTTGTTGCGGAGCGCGCCGAC<br>GCCCTGAACGCCCTTGAAGAGCTGTACGAAGACCGCGCGGCAGGCGCGTACGACGGACCCGTTGGCAGGAAGCACTT<br>CCGGAAGCAACAGGCAGCGCTGACGCTCCGGCAGCAAGGGGCGGAAGAGCGGCTTGCCGAACTTGAAGCCGCCGAA<br>GCCCCGAAGCTTCCCCTTGACCAATGGTTCCCCGAAGACGCCGACGCTGACCCGACCGGCCCTAAGTCGTGGTGGGG<br>GCGCGCGTCAGTAGACGACAAGCGCGTGTTCGTCGGGCTCTTCGTAGACAAGATCGTTGTCACGAAGTCGACTACGGG<br>CAGGGGGCAGGGAACGCCCATCGAGAAGCGCGCTTCGATCACGTGGGCGAAGCCGCCGACCGACGACGACGAAGAC<br>GACGCCCAGGACGGCACGGAAGACGTAGCGGCGTAATAACTCGAGGCGAATTTCTTATGATTTATGATTTTTATTATTAA<br>ATAAGTTATAAAAAAAATAAGTGTATACAAATTTTAAAGTGACTCTTAGGTTTTAAAACGAAAATTCTTATTCTTGAGTAACT<br>CTTTCCTGTAGGTCAGGTTGCTTTCTCAGGTATAGCATGAGGTCGCTCTTATTGACCACACCTCTACCGGCATGCCGAG<br>CAAATGCCTGCAAATCGCTCCCCATTTCAGCAAAAGGCCAGCAAAAGGCCAGGAACCGTAAAAAGGCCGC |
| pOP262 | Plamids for generation of donor DNA for construction of a PhiBT-compatible landing pad p1. After linearization with ScaI, this plasmid produces donor DNA with the *URA3* selectable marker, *attB* recombination sites for PhiBT, and homology flanks matches the original landing pad p1 used in the lab[1]. This plasmid contains AmpR marker for propagation in *E. coli.* | CAGTAAGAGAATTATGCAGTGCTGCCATAACCATGAGTGATAACACTGCGGCCAACTTACTTCTGACAACGATCGGAGG<br>ACCGAAGGAGCTAACCGCTTTTTTGCACAACATGGGGGATCATGTAACTCGCCTTGATCGTTGGGAACCGGAGCTGAAT<br>GAAGCCATACCAAACGACGAGCGTGACACCACGATGCCTGTAGCAATGGCAACAACGTTGCGCAAACTATTAACTGGC<br>GAACTACTTACTCTAGCTTCCCGGCAACAATTAATAGACTGGATGGAGGCGGATAAAGTTGCAGGACCACTTCTGCGCT<br>CGGCCCTTCCGGCTGGCTGGTTTATTGCTGATAAATCTGGAGCCGGTGAGCGTGGGTCTCGCGGTATCATTGCAGCAC<br>TGGGGCCAGATGGTAAGCCCTCCCGTATCGTAGTTATCTACACGACGGGGAGTCAGGCAACTATGGATGAACGAAATA<br>GACAGATCGCTGAGATAGGTGCCTCACTGATTAAGCATTGGTAACTGTCAGACCAAGTTTACTCATATATACTTTAGATT<br>GATTTAAAACTTCATTTTTAATTTAAAAGGATCTAGGTGAAGATCCTTTTTGATAATCTCATGACCAAAATCCCTTAACGTG<br>AGTTTTCGTTCCACTGAGCGTCAGACCCCGTAGAAAAGATCAAAGGATCTTCTTGAGATCCTTTTTTTCTGCGCGTAATC<br>TGCTGCTTGCAAACAAAAAAACCACCGCTACCAGCGGTGGTTTGTTTGCCGGATCAAGAGCTACCAACTCTTTTTCCGAA<br>GGTAACTGGCTTCAGCAGAGCGCAGATACCAAATACTGTTCTTCTAGTGTAGCCGTAGTTAGGCCACCACTTCAAGAAC<br>TCTGTAGCACCGCCTACATACCTCGCTCTGCTAATCCTGTTACCAGTGGCTGCTGCCAGTGGCGATAAGTCGTGTCTTA<br>CCGGGTTGGACTCAAGACGATAGTTACCGGATAAGGCGCAGCGGTCGGGCTGAACGGGGGGTTCGTGCACACAGCCC<br>AGCTTGGAGCGAACGACCTACACCGAACTGAGATACCTACAGCGTGAGCTATGAGAAAGCGCCACGCTTCCCGAAGGG<br>AGAAAGGCGGACAGGTATCCGGTAAGCGGCAGGGTCGGAACAGGAGAGCGCACGAGGGAGCTTCCAGGGGGAAACG<br>CCTGGTATCTTTATAGTCCTGTCGGGTTTCGCCACCTCTGACTTGAGCGTCGATTTTTGTGATGCTCGTCAGGGGGGCG<br>GAGCCTATGGAAAAACGCCAGCAACGCGGCCTTTTTACGGTTCCTGGCCTTTTGCTGGCCTTTTGCTCACATGTTCTTTC<br>CTGCGTTATCCCCTGATTCTGTGGATAACCGTATTACCGCCTTTGAGTGAGCTGATACCGCTCGCCGCAGCCGAACGAC<br>CGAGCGCAGCGAGTCAGTGAGCGAGGAAGCGGAAGAGCGCCCAATACGCAAACCGCCTCTCCCCGCGCGTTGGCCG<br>ATTCATTAATGCAGCTGGCACGACAGGTTTCCCGACTGGAAAGCGGGCAGTGAGCGCAACGCAATTAATAGTACTATCT<br>ATTTTTTCAGTACATAATTTAATTAAATCATCATGACCTTTATCTAATTTTACATCTCTTCTACCAAGACCTAATTCTTTATTA<br>ATACTATCAAAATCTGTTATTCTATTGCCTTGAGGATCTTTGAAACACCAATCTGAGTCTGAGGGCATGCCCAGGTTTTTG<br>ACGAAAGTGATCCAGATGATCCAGCTTTTTCGCCGGAGTCAATTAGGTCATACTTTTCTATATAATCCAAATCCCAAAAAT<br>CAATTGAATGATTCTTAATATGATTTAATAGTTTATGATTATAAATGTCGAAAGCTACATATAAGGAACGTGCTGCTACTCA<br>TCCTAGTCCTGTTGCTGCCAAGCTATTTAATATCATGCACGAAAAGCAAACAAACTTGTGTGCTTCATTGGATGTTCGTAC<br>CACCAAGGAATTACTGGAGTTAGTTGAAGCATTAGGTCCCAAAATTTGTTTACTAAAAACACATGTGGATATCTTGACTGA<br>TTTTTCCATGGAGGGCACAGTTAAGCCGCTAAAGGCATTATCCGCCAAGTACAATTTTTTACTCTTCGAGGACAGAAAAT<br>TTGCTGACATTGGTAATACAGTCAAATTGCAGTACTCTGCGGGTGTATACAGAATAGCAGAATGGGCAGACATTACGAAT<br>GCACACGGTGTGGTGGGCCCAGGTATTGTTAGCGGTTTGAAGCAGGCGGCAGAAGAAGTAACAAAGGAACCTAGAGG<br>CCTTTTGATGTTAGCAGAATTGTCATGCAAGGGCTCCCTATCTACTGGAGAATATACTAAGGGTACTGTTGACATTGCGA<br>AGAGCGACAAAGATTTTGTTATCGGCTTTATTGCTCAAAGAGACATGGGTGGAAGAGATGAAGGTTACGATTGGTTGATT<br>ATGACACCCGGTGTGGGTTTAGATGACAAGGGAGATGCATTGGGTCAACAGTATAGAACCGTGGATGATGTGGTTTCTA<br>CAGGATCTGACATTATTATTGTTGGAAGAGGACTATTTGCAAAGGGAAGGGATGCTAAGGTAGAGGGTGAACGTTACAG<br>AAAAGCAGGCTGGGAAGCATATTTGAGAAGATGCGGCCAGCAAAACTAAGCTGGATCATCTGGATCTTTTTCGTCAAAA<br>ACCTGGAATCCATAATGGCTTCATCTAATGTGGACTCTTCTGAGCCCCGTATTTCCAGAGAAAGCTCTTTGAAGAGGTCT<br>CTATCTATTTCAAAGTCGTTAAAAGGGTTGTTCAAGTCAGGAGGGGGAAACGCCAATACTGGCCCTACTACTGCTGCTG<br>CTGCTGCTGCACCATCTAGTATCTTTTAAATTAATAATGTAATTCATATATTATAGTACTCGTCTCCGGGAGCTGCATGTG<br>TCAGAGGTTTTCACCGTCATCACCGAAACGCGTGAGACGAAAGGGCCTCGTGATACGCCTATTTTTATAGGTTAATGTCA<br>TGATAATAATGGTTTCTTAGACGTCAGGTGGCACTTTTCGGGGAAATGTGCGCGGAACCCCTATTTGTTTATTTTTCTAAA<br>TACATTCAAATATGTATCCGCTCATGAGACAATAACCCTGATAAATGCTTCAATAATATTGAAAAAGGAAGAGTATGAGTA<br>TTCAACATTTCCGTGTCGCCCTTATTCCCTTTTTTGCGGCATTTTGCCTTCCTGTTTTTGCTCACCCAGAAACGCTGGTGA<br>AAGTAAAAGATGCTGAAGATCAGTTGGGTGCACGAGTGGGTTACATCGAACTGGATCTCAACAGCGGTAAGATCCTTGA<br>GAGTTTTCGCCCCGAAGAACGTTTTCCAATGATGAGCACTTTTAAAGTTCTGCTATGTGGCGCGGTATTATCCCGTATTG<br>ACGCCGGGCAAGAGCAACTCGGTCGCCGCATACACTATTCTCAGAATGACTTGGTTGAGTACTCACCAGTCACAGAAAA<br>GCATCTTACGGATGGCATGA |
| pOP264 | 2*μ* plasmid for constitutive expression of PhiBT under RNR2 promoter. The plasmid includes a kanMX marker for | CCGACAGGACTATAAAGATACCAGGCGTTTCCCCCTGGAAGCTCCCTCGTGCGCTCTCCTGTTCCGACCCTGCCGCTTA<br>CCGGATACCTGTCCGCCTTTCTCCCTTCGGGAAGCGTGGCGCTTTCTCATAGCTCACGCTGTAGGTATCTCAGTTCGGT<br>GTAGGTCGTTCGCTCCAAGCTGGGCTGTGTGCACGAACCCCCCGTTCAGCCCGACCGCTGCGCCTTATCCGGTAACTA<br>TCGTCTTGAGTCCAACCCGGTAAGACACGACTTATCGCCACTGGCAGCAGCCACTGGTAACAGGATTAGCAGAGCGAG<br>GTATGTAGGCGGTGCTACAGAGTTCTTGAAGTGGTGGCCTAACTACGGCTACACTAGAAGGACAGTATTTGGTATCTGC<br>GCTCTGCTGAAGCCAGTTACCTTCGGAAAAAGAGTTGGTAGCTCTTGATCCGGCAAACAAACCACCGCTGGTAGCGGT<br>GGTTTTTTTGTTTGCAAGCAGCAGATTACGCGCAGAAAAAAAGGATCTCAAGAAGATCCTTTGATCTTTTCTACACTAGTC<br>GAAGCATCTGTGCTTCATTTTGTAGAACAAAAATGCAACGCGAGAGCGCTAATTTTTCAAACAAAGAATCTGAGCTGCAT<br>TTTTACAGAACAGAAATGCAACGCGAAAGCGCTATTTTACCAACGAAGAATCTGTGCTTCATTTTTGTAAAACAAAAATGC<br>AACGCGAGAGCGCTAATTTTTCAAACAAAGAATCTGAGCTGCATTTTTACAGAACAGAAATGCAACGCGAGAGCGCTATT<br>TTACCAACAAAGAATCTATACTTCTTTTTTGTTCTACAAAAATGCATCCCGAGAGCGCTATTTTTCTAACAAAGCATCTTAG<br>ATTACTTTTTTTCTCCTTTGTGCGCTCTATAATGCAGTCTCTTGATAACTTTTTGCACTGTAGGTCCGTTAAGGTTAGAAGA |

| | | |
|---|---|---|
| | selection in yeast and propagation in *E.coli*. | AGGCTACTTTGGTGTCTATTTTCTCTTCCATAAAAAAAGCCTGACTCCACTTCCCGCGTTTACTGATTACTAGCGAAGCTG<br>CGGGTGCATTTTTTCAAGATAAAGGCATCCCCGATTATATTCTATACCGATGTGGATTGCGCATACTTTGTGAACAGAAA<br>GTGATAGCGTTGATGATTCTTCATTGGTCAGAAAATTATGAACGGTTTCTTCTATTTTGTCTCTATATACTACGTATAGGAA<br>ATGTTTACATTTTCGTATTGTTTTCGATTCACTCTATGAATAGTTCTTACTACAATTTTTTTGTCTAAAGAGTAATACTAGAG<br>ATAAACATAAAAAATGTAGAGGTCGAGTTTAGATGCAAGTTCAAGGAGCGAAAGGTGGATGGGTAGGTTATATAGGGAT<br>ATAGCACAGAGATATATAGCAAAGAGATACTTTTGAGCAATGTTTGTGGAAGCGGTATTCGCAATATTTTAGTAGCTCGTT<br>ACAGTCCGGTGCGTTTTTGGTTTTTTGAAAGTGCGTCTTCAGAGCGCTTTTGGTTTTCAAAAGCGCTCTGAAGTTCCTAT<br>ACTTTCTAGAGAATAGGAACTTCGGAATAGGAACTTCAAAGCGTTTCCGAAAACGAGCGCTTCCGAAAATGCAACGCGA<br>GCTGCGCACATACAGCTCACTGTTCACGTCGCACCTATATCTGCGTGTTGCCTGTATATATATATACATGAGAAGAACGG<br>CATAGTGCGTGTTTATGCTTAAATGCGTATATGTGTTATGTATAGGTCTAGAGATCTGTTTAGCTTGCCTCGTCCCCGCC<br>GGGTCACCCGGCCAGCGACATGGAGGCCCAGAATACCCTCCTTGACAGTCTTGACGTGCGCAGCTCAGGGGCATGAT<br>GTGACTGTCGCCCGTACATTTAGCCCATACATCCCCATGTATAATCATTTGCATCCATACATTTTGATGGCCGCACGGCG<br>CGAAGCAAAAATTACGGCTCCTCGCTGCAGACCTGCGAGCAGGGAAACGCTCCCCTCACAGACGCGTTGAATTGTCCC<br>CACGCCGCGCCCCTGTAGAGAAATATAAAAGGTTAGGATTTGCCACTGAGGTTCTTCTTTCATATACTTCCTTTTAAAATC<br>TTGCTAGGATACAGTTCTCACATCACATCCGAACATAAACAACCATGGGTAAGGAAAAGACTCACGTTTCGAGGCCGCG<br>ATTAAATTCCAACATGGATGCTGATTTATATGGGTATAAATGGGCTCGCGATAATGTCGGGCAATCAGGTGCGACAATCT<br>ATCGATTGTATGGGAAGCCCGATGCGCCAGAGTTGTTTCTGAAACATGGCAAAGGTAGCGTTGCCAATGATGTTACAGA<br>TGAGATGGTCAGACTAAACTGGCTGACGGAATTTATGCCTCTTCCGACCATCAAGCATTTTATCCGTACTCCTGATGATG<br>CATGGTTACTCACCACTGCGATCCCCGGCAAAACAGCATTCCAGGTATTAGAAGAATATCCTGATTCAGGTGAAAATATT<br>GTTGATGCGCTGGCAGTGTTCCTGCGCCGGTTGCATTCGATTCCTGTTTGTAATTGTCCTTTTAACAGCGATCGCGTATT<br>TCGTCTCGCTCAGGCGCAATCACGAATGAATAACGGTTTGGTTGATGCGAGTGATTTTGATGACGAGCGTAATGGCTGG<br>CCTGTTGAACAAGTCTGGAAAGAAATGCATAAGCTTTTGCCATTCTCACCGGATTCAGTCGTCACTCATGGTGATTTCTC<br>ACTTGATAACCTTATTTTTGACGAGGGGAAATTAATAGGTTGTATTGATGTTGGACGAGTCGGAATCGCAGACCGATACC<br>AGGATCTTGCCATCCTATGGAACTGCCTCGGTGAGTTTTCTCCTTCATTACAGAAACGGCTTTTTCAAAAATATGGTATTG<br>ATAATCCTGATATGAATAAATTGCAGTTTCATTTGATGCTCGATGAGTTTTTCTAATCAGTACTGACAATAAAAAGATTCTT<br>GTTTTCAAGAACTTGTCATTTGTATAGTTTTTTTATATTGTAGTTGTTCTATTTTAATCAAATGTTAGCGTGATTTATATTTTT<br>TTTCGCCTCGACATCATCTGCCCAGATGCGAAGTTAAGTGCGCAGAAAGTAATATCATGCGTCAATCGTATGTGAATGCT<br>GGTCGCTATACTGCTGTCGATTCGATACTAACGCCGCCATCCAGTGTCGAAAACGAGCTCTCGAGAACCCTTAATGTCG<br>ACAGTCGAACAAGAAGCAGGCAAAGTTTAGAGCACTGCCCCTCCGCACTCAAAAAAGAAAAAACTAGGAGGAAAATAAA<br>ATTCTCAACCACACAAACACATAAACACATACAAATACAAATACAAGCTTATTTACTTGACATCGCGCGATCTTCCACTAT<br>TCAGCGCCGTCCGCCCTCTCTCGTGTTTTTTGTTTACGCGACAACTATGCGAAATCCGGAGCAACGGGCAACCGTTTGG<br>GGAAAGACCACACCCACGCGCGATCGCCATGGCAACGAGGTCGCACACGCCCCACACCCAGACCTCCCTGCGAGCGG<br>GCATGGGTACAATGTCCCCGTTGCCACAGACACCACTTCGTAGCACAGCGCAGAGCGTAGCGTGTTGTTGCTGCTGAC<br>AAAAGAAAATTTTTCTTAGCAAAGCAAAGGAGGGGAAGCACGGGCAGATAGCACCGTACCATACCCTTGGAAACTCGAA<br>ATGAACGAAGCAGGAAATGAGAGAATGAGAGTTTTGTAGGTATATATAGCGGTAGTGTTTGCGCGTTACCATCATCTTCT<br>GGATCTATCTATTGTTCTTTTCCTCATCACTTTCCCCTTTTTCGCTCTTCTTCTTGTCTTTTATTTCTTTCTTTTTTTTAATTG<br>TTCCCTCGATTGGCTATCTACCAAAGAATCCAAACTTAATACACGTATTTATTTGTCCAATTACCATGAGCCCATTCATTG<br>CACCTGATGTTCCAGAACATTTGCTTGATACGGTGAGAGTGTTTCTTTACGCTCGCCAATCTAAAGGTAGGTCGGATGGT<br>AGTGATGTCTCAACCGAAGCGCAGCTGGCCGCTGGAAGAGCTTTAGTTGCCTCCAGGAACGCACAAGGCGGTGCGAG<br>GTGGGTAGTAGCTGGTGAATTTGTTGACGTTGGTAGATCCGGTTGGGACCCTAACGTTACACGTGCCGACTTTGAAAGG<br>ATGATGGGTGAGGTCAGAGCTGGAGAGGGTGACGTTGTGGTCGTTAACGAATTGTCTCGGTTAACAAGGAAGGGCGCC<br>CATGATGCACTGGAGATTGATAATGAATTAAAAAAACACGGAGTCAGATTCATGTCTGTGTTAGAGCCCTTTTTGGATAC<br>GTCTACACCGATCGGTGTAGCAATCTTCGCTTTGATAGCTGCCCTTGCTAAGCAAGATAGTGATTTAAAAGCTGAGCGTT<br>TGAAGGGAGCCAAGGATGAAATAGCGGCGCTAGGGGGTGTTCATTCTTCCTCTGCTCCTTTTGGTATGCGTGCAGTTCG<br>GAAAAAGGTAGATAATCTTGTTATTTCTGTTCTCGAGCCAGATGAAGATAACCCAGACCACGTCGAACTGGTTGAGAGGA<br>TGGCTAAGATGAGTTTTGAAGGTGTATCAGATAATGCAATAGCAACAACTTTTGAAAAGGAGAAGATCCCGTCTCCCGGT<br>ATGGCGGAAAGACGAGCCACCGAGAAGAGATTAGCAAGTATCAAAGCTCGTCGTTTAAACGGGGCTGAAAAACCAATTA<br>TGTGGAGAGCGCAAACCGTAAGATGGATTCTAAATCATCCAGCCATTGGTGGCTTTGCTTTCGAAAGAGTTAAGCATGG<br>AAAAGCTCATATCAACGTAATACGACGCGACCCAGGGGGCAAACCTTTAACTCCACATACTGGTATACTATCAGGCTCAA<br>AATGGTTGGAGCTACAAGAAAAAAGGAGCGGTAAGAATTTAAGCGACAGAAAACCCGGCGCTGAAGTGGAACCTACACT<br>ACTGTCTGGGTGGAGGTTTTTGGGCTGCAGAATTTGTGGAGGTTCTATGGGTCAATCACAAGGAGGAAGAAAACGCAAT<br>GGTGATCTTGCTGAAGGCAATTACATGTGTGCCAATCCTAAAGGTCATGGGGGATTAAGTGTTAAAAGGTCCGAGTTGG<br>ATGAGTTTGTAGCAAGTAAGGTCTGGGCCAGATTAAGAACCGCAGACATGGAAGACGAACACGATCAAGCTTGGATTGC<br>TGCAGCTGCTGAACGATTTGCATTGCAGCACGATTTAGCCGGAGTCGCAGACGAAAGAAGAGAACAACAGGCACATCTT<br>GACAATGTGAGACGTTCGATTAAGGACCTACAAGCAGATAGAAAGGCCGGGTTGTATGTAGGTAGAGAAGAACTAGAAA<br>CTTGGAGATCAACGGTTTTGCAGTATCGTTCGTATGAAGCTGAGTGTACTACTCGATTGGCCGAATTAGATGAAAAAATG<br>AACGGCTCGACAAGGGTGCCTAGCGAGTGGTTCAGCGGTGAAGATCCGACTGCAGAAGGTGGAATATGGGCTTCCTG<br>GGACGTCTATGAAAGAAGAGAGTTCTTATCCTTCTTCTTGGATTCAGTCATGGTTGATAGAGGCAGGCACCCAGAAACTA<br>AGAAATACATTCCACTCAAAGATAGAGTGACACTTAAATGGGCCGAATTGCTCAAAGAAGAGGATGAAGCATCAGAGGC<br>TACCGAAAGAGAATTGGCAGCGCTGTAAATCCGCTCTAACCGAAAAGGAAGGAGTTAGACAACCTGAAGTCTAGGTCCC<br>TATTTATTTTTTTATAGTTATGTTAGTATTAAGAACGTTATTTATATTTCAAATTTTTCTTTTTTTTCTGTACAGACGCGTGTA<br>CGCATGTAACATTATACTGAAAACCTTGCTTGAGAAGGTTTTGGGACGCTCGAAGCCGCGGTGAGCAAAAGGCCAGCAA<br>AAGGCCAGGAACCGTAAAAAGGCCGCGTTGCTGGCGTTTTTCCATAGGCTCCGCCCCCCTGACGAGCATCACAAAAAT<br>CGACGCTCAAGTCAGAGGTGGCGAAAC |
| pOP266 | Plasmid for genertaion of donor DNA for PhiBT-compatible p1 integration in yeast. After linearization with ScaI, this plasmid produces donor DNA encoding the fluorescent protein mKate as a reporter with a *LEU2* auxotrophic selection marker, flanked by *attP* recombination sites for PhiC31. This | TTGAGATCCTTTTTTTCTGCGCGTAATCTGCTGCTTGCAAACAAAAAAACCACCGCTACCAGCGGTGGTTTGTTTGCCGG<br>ATCAAGAGCTACCAACTCTTTTTCCGAAGGTAACTGGCTTCAGCAGAGCGCAGATACCAAATACTGTTCTTCTAGTGTAG<br>CCGTAGTTAGGCCACCACTTCAAGAACTCTGTAGCACCGCCTACATACCTCGCTCTGCTAATCCTGTTACCAGTGGCTG<br>CTGCCAGTGGCGATAAGTCGTGTCTTACCGGGTTGGACTCAAGACGATAGTTACCGGATAAGGCGCAGCGGTCGGGCT<br>GAACGGGGGGTTCGTGCACACAGCCCAGCTTGGAGCGAACGACCTACACCGAACTGAGATACCTACAGCGTGAGCTAT<br>GAGAAAGCGCCACGCTTCCCGAAGGGAGAAAGGCGGACAGGTATCCGGTAAGCGGCAGGGTCGGAACAGGAGAGCG<br>CACGAGGGAGCTTCCAGGGGGAAACGCCTGGTATCTTTATAGTCCTGTCGGGTTTCGCCACCTCTGACTTGAGCGTCG<br>ATTTTTGTGATGCTCGTCAGGGGGGCGGAGCCTATGGAAAAACGCCAGCAACGCGGCCTTTTTACGGTTCCTGGCCTTT<br>TGCTGGCCTTTTGCTCACATGTTCTTTCCTGCGTTATCCCCTGATTCTGTGGATAACCGTATTACCGCCTTTGAGTGAGC<br>TGATACCGCTCGCCGCAGCCGAACGACCGAGCGCAGCGAGTCAGTGAGCGAGGAAGCGGAAGAGCGCCCAATACGCA<br>AACCGCCTCTCCCCGCGCGTTGGCCGATTCATTAATGCAGCTGGCACGACAGGTTTCCCGACTGGAAAGCGGGCAGTG<br>AGCGCAACGCAATTAATAGTACTATCTATTTTTTCAGTACATAATTTAATTAAATCATCATGACCTTTATCTAATTTTACATC<br>TCTTCTACCAAGACCTAATTCTTTATTAATACTATCAAAATCTGTTATTCTATTGCCTTGAGGATCTTTGAAACACCAATCT<br>GAGTCTGAGGTGCTGGGTTGTTGTCTCTGGACAGTGATCCATGGGAAACTACTCAGCAGATGACCTATACATAGGAAGA<br>TCTATAGAAACAAAAAGATTAATAACTTTCAAATATCAGAAAAATGTAGAAACATGTGATAAGCTCATAGACATGTAAAATG<br>CATGTGAGCGAGCTGATTAAGGAGAACATGCACATGAAGCTGTACATGGAGGGCACCGTGAACAACCACCACTTCAAGT<br>GCACATCCGAGGGCGAAGGCAAGCCCTACGAGGGCACCCAGACCATGAGAATCAAGGCGGTCGAGGGCGGCCCTCTC<br>CCCTTCGCCTTCGACATCCTGGCTACCAGCTTCATGTACGGCAGCAAAACCTTCATCAACCACACCCAGGGCATCCCCG<br>ACTTCTTTAAGCAGTCCTTCCCCGAGGGCTTCACATGGGAGAGAGTCACCACATACGAAGACGGGGGCGTGCTGACCG<br>CTACCCAGGACACCAGCCTCCAGGACGGCTGCCTCATCTACAACGTCAAGATCAGAGGGGTGAACTTCCCATCCAACG<br>GCCCTGTGATGCAGAAGAAAACACTCGGCTGGGAGGCCTCCACCGAGACCCTGTACCCCGCTGACGGCGGCCTGGAA<br>GGCAGAGCCGACATGGCCCTGAAGCTCGTGGGCGGGGGCCACCTGATCTGCAACTTGAAGACCACATACAGATCCAA<br>GAAACCCGCTAAGAACCTCAAGATGCCCGGCGTCTACTATGTGGACAGAAGACTGGAAAGAATCAAGGAGGCCGACAA<br>AGAGACCTACGTCGAGCAGCACGAGGTGGCTGTGGCCAGATACTGCGACCTCCCTAGCAAACTGGGGCACAGATAAG<br>CATGCTTTTTCGCCGGAGTCAATTAGGTCATACTTTTCTATATAATCCAAATCCCAAAAATCAATTGAATGATTCTTAATAT<br>GATTTAATAGTTTATGATTATAAATGCACGTTCCTAAGAAGATCGTCGTTTTGCCAGGTGACCACGTTGGTCAAGAAATCA<br>CAGCCGAAGCCATTAAGGTTCTTAAAGCTATTTCTGATGTTCGTTCCAATGTCAAGTTCGATTTCGAAAATCATTTAATTG<br>GTGGTGCTGCTATCGATGCTACAGGTGTTCCACTTCCAGATGAGGCGCTGGAAGCCTCCAAGAAGGCTGATGCCGTTTT<br>GTTAGGTGCTGTGGGTGGTCCTAAATGGGGTACCGGTAGTGTTAGACCTGAACAAGGTTTACTAAAAATCCGTAAAGAA<br>CTTCAATTGTACGCCAACTTAAGACCATGTAACTTTGCATCCGACTCTCTTTTAGACTTATCTCCAATCAAGCCACAATTT<br>GCTAAAGGTACTGACTTCGTTGTTGTCAGAGAATTAGTGGGAGGTATTTACTTTGGTAAGAGAAAGGAAGACGATGGTG<br>ATGGTGTCGCTTGGGATAGTGAACAATACACCGTTCCAGAAGTGCAAAGAATCACAAGAATGGCCGCTTTCATGGCCCT<br>ACAACATGAGCCACCATTGCCTATTTGGTCCTTGGATAAAGCTAATGTTTTGGCCTCTTCAAGATTATGGAGAAAAACTG |

| | | |
|---|---|---|
| | plasmid contains AmpR marker for propagation in *E. coli.* | TGGAGGAAACCATCAAGAACGAATTCCCTACATTGAAGGTTCAACATCAATTGATTGATTCTGCCGCCATGATCCTAGTT<br>AAGAACCCAACCCACCTAAATGGTATTATAATCACCAGCAACATGTTTGGTGATATCATCTCCGATGAAGCCTCCGTTAT<br>CCCAGGTTCCTTGGGTTTGTTGCCATCTGCGTCCTTGGCCTCTTTGCCAGACAAGAACACCGCATTTGGTTTGTACGAA<br>CCATGCCACGGTTCTGCTCCAGATTTGCCAAAGAATAAGGTCAACCCTATCGCCACTATCTTGTCTGCTGCAATGATGTT<br>GAAATTGTCATTGAACTTGCCTGAAGAAGGTAAGGCCATTGAAGATGCAGTTAAAAAGGTTTTGGATGCAGGTATCAGAA<br>CTGGTGATTTAGGTGGTTCCAACAGTACCACCGAAGTCGGTGATGCTGTCGCCGAAGAAGTTAAGAAAATCCTTGCTTA<br>ATGCTGAGTAGTTTCCCATGGATCTTTGTCCAGAGACAACAACCCAGCAAATCCATAATGGCTTCATCTAATGTGGACTC<br>TTCTGAGCCCCGTATTTCCAGAGAAAGCTCTTTGAAGAGGTCTCTATCTATTTCAAAGTCGTTAAAAGGGTTGTTCAAGT<br>CAGGAGGGGGAAACGCCAATACTGGCCCTACTACTGCTGCTGCTGCTGCTGCACCATCTAGTATCTTTTAAATTAATAAT<br>GTAATTCATATATTATAGTACTCGTCTCCGGGAGCTGCATGTGTCAGAGGTTTTCACCGTCATCACCGAAACGCGTGAGA<br>CGAAAGGGCCTCGTGATACGCCTATTTTTATAGGTTAATGTCATGATAATAATGGTTTCTTAGACGTCAGGTGGCACTTTT<br>CGGGGAAATGTGCGCGGAACCCCTATTTGTTTATTTTTCTAAATACATTCAAATATGTATCCGCTCATGAGACAATAACCC<br>TGATAAATGCTTCAATAATATTGAAAAAGGAAGAGTATGAGTATTCAACATTTCCGTGTCGCCCTTATTCCCTTTTTTGCG<br>GCATTTTGCCTTCCTGTTTTTGCTCACCCAGAAACGCTGGTGAAAGTAAAAGATGCTGAAGATCAGTTGGGTGCACGAG<br>TGGGTTACATCGAACTGGATCTCAACAGCGGTAAGATCCTTGAGAGTTTTCGCCCCGAAGAACGTTTTCCAATGATGAG<br>CACTTTTAAAGTTCTGCTATGTGGCGCGGTATTATCCCGTATTGACGCCGGGCAAGAGCAACTCGGTCGCCGCATACAC<br>TATTCTCAGAATGACTTGGTTGAGTACTCACCAGTCACAGAAAAGCATCTTACGGATGGCATGACAGTAAGAGAATTATG<br>CAGTGCTGCCATAACCATGAGTGATAACACTGCGGCCAACTTACTTCTGACAACGATCGGAGGACCGAAGGAGCTAACC<br>GCTTTTTTGCACAACATGGGGGATCATGTAACTCGCCTTGATCGTTGGGAACCGGAGCTGAATGAAGCCATACCAAACG<br>ACGAGCGTGACACCACGATGCCTGTAGCAATGGCAACAACGTTGCGCAAACTATTAACTGGCGAACTACTTACTCTAGC<br>TTCCCGGCAACAATTAATAGACTGGATGGAGGCGGATAAAGTTGCAGGACCACTTCTGCGCTCGGCCCTTCCGGCTGG<br>CTGGTTTATTGCTGATAAATCTGGAGCCGGTGAGCGTGGGTCTCGCGGTATCATTGCAGCACTGGGGCCAGATGGTAA<br>GCCCTCCCGTATCGTAGTTATCTACACGACGGGGAGTCAGGCAACTATGGATGAACGAAATAGACAGATCGCTGAGATA<br>GGTGCCTCACTGATTAAGCATTGGTAACTGTCAGACCAAGTTTACTCATATATACTTTAGATTGATTTAAAACTTCATTTTT<br>AATTTAAAAGGATCTAGGTGAAGATCCTTTTTGATAATCTCATGACCAAAATCCCTTAACGTGAGTTTTCGTTCCACTGAG<br>CGTCAGACCCCGTAGAAAAGATCAAAGGATCTTC |

**Table S3. Key strains used in this study.**

| Yeast | Description | Genome |
|---|---|---|
| BJ5465 | Parent strain | MATa ura3-52 trp1 leu2-delta1 his3-delta200 pep4::HIS3 prb1-delta1.6R can1 GAL |
| yOP109 | Parent strain | MATα his3Δ1 leu2Δ0 met15Δ0 lys2Δ0 ura3Δ0 Can1::WT DNA-Hyg; Landing pad p1-MET15 |
| yYY20 | Strain that contained landing pad p1 with *attB* recombination sites and *URA3* selectable marker | MATα his3Δ1 leu2Δ0 met15Δ0 lys2Δ0 ura3Δ0 Can1::WT DNAP1-Hyg; Landing pad p1-MET15::Landing pad p1-*attB*-URA3 |
| yYY22 | Strain that contains 1) landing pad p1 with *attB* recombination sites and *URA3* selectable marker; and 2) a 2*μ* plasmid for constitutive expression of TP901 integrase. | MATα his3Δ1 leu2Δ0 met15Δ0 lys2Δ0 ura3Δ0 Can1::WT DNAP1-Hyg; Landing pad p1-*attB*-Ura3; pYY10 |
| yYY64 | Strain that contained landing pad p1 with *attB* recombination sites and *TPR1* selectable marker | MATαlpha his3Δ1 leu2Δ0 met15Δ0 lys2Δ0 ura3Δ0 SSD1Δ0 CAN1::PSP2 WT NAT HO::Rad27 Lys2 Ade2Δ Trp1Δ; Landing pad p1-MET15::Landing pad p1-*attB*-TRP1 |
| yYY206 | β-estradiol inducible yeast surface display strain with landing pad p1 containing *attB* recombination sites and *TPR1* selectable marker | MATa ura3-52 trp1Δ leu2Δ1 his3-Δ200 pep4::HIS3 prb1-Δ1.6R can1 GAL kar1Δ15 lyp1::WT DNAP1-URA3-CAN1 Aga1p::pBeta estradiol aga1p – synTF-HygR; Landing pad p1-*attB*-TRP1 |
| yYY235 | β-estradiol inducible yeast surface display strain with 1) landing pad p1 containing *attB* recombination sites and *TPR1* selectable marker; and 2) a 2*μ* plasmid for constitutive expression of TP901 integrase | MATa ura3-52 trp1Δ leu2Δ1 his3-Δ200 pep4::HIS3 prb1-Δ1.6R can1 GAL kar1Δ15 lyp1::WT DNAP1-URA3-CAN1 Aga1p::pBeta estradiol aga1p-synTF-HygR; Landing pad p1-*attB*-TRP1; pYY10 |
